# Regulation of stereotypic chromatin conformations at enhancers by CLOCK:BMAL1

**DOI:** 10.1101/2024.04.24.590818

**Authors:** Xinyu Y. Nie, Neha Ahuja, Amber Hsieh, Garima Abbi, Yuchao Jiang, Jerome S. Menet

## Abstract

Cooperation between the circadian transcription factor (TF) CLOCK:BMAL1 and other TFs at *cis*-regulatory elements (CREs) is critical to daily rhythms of transcription. Yet, the mechanisms underlying this cooperation are unclear. Here, we analyzed the co-binding of multiple TFs on single DNA molecules in mouse liver using single molecule footprinting (SMF). We found that SMF reads clustered in stereotypic chromatin states that reflect distinguishable organization of TFs and nucleosomes, and that were remarkably conserved between all samples. DNA protection at CLOCK:BMAL1 binding motif (E-box) varied between CREs, from E-boxes being solely bound by CLOCK:BMAL1 to situations where other TFs competed with CLOCK:BMAL1 for E-box binding. SMF also uncovered CLOCK:BMAL1 cooperative binding at E-boxes separated by 250 bp, which structurally altered the CLOCK:BMAL1-DNA interface. Importantly, we discovered multiple nucleosomes with E-boxes at entry/exit sites that were removed upon CLOCK:BMAL1 DNA binding, thereby promoting the formation of open chromatin states that facilitate DNA binding of other TFs and that were associated with rhythmic transcription. These results demonstrate the utility of SMF for studying how CLOCK:BMAL1 and other TFs regulate stereotypical chromatin states at CREs to promote transcription.

## Introduction

Cooperation between TFs at CREs (defined here as enhancers and promoters) is critical for transcription activation^1–11^. Such cooperativity can involve direct TF-TF interaction, but it can also occur without direct contact through a mechanism called nucleosome-mediated TF cooperation^6,8,12^. Because most TFs preferentially bind naked DNA^13^, nucleosomes act as a physical barrier that limits TF-DNA binding. Since TFs occupy much smaller footprints on DNA than histone octamers (< 30 bp *vs* 147 bp)^14^, binding of a TF on DNA can not only disrupt nucleosome compaction but also facilitate the binding of additional TFs nearby^6,12,15–20^. However, how TFs cooperate to compete with histones, access DNA, and regulate CRE activity remains unclear and is a central question in the field of gene expression.

By switching from an active to inactive state over the 24-hour cycle, circadian TFs provide a unique system to study the role of TF cooperativity in transcriptional regulation. In mammals, nearly every cell harbors a circadian clock that drives daily rhythms in gene expression, aligning biological processes with the appropriate time of day. Given the large number of rhythmically expressed genes, most transcriptional programs and biological pathways show oscillatory activity across the day, as for example those controlling metabolic functions in anticipation of the 24-hour feed-fasting rhythm^21,22^. The mammalian circadian clock is initiated by the heterodimeric basic-helix-loop-helix (bHLH) TF CLOCK:BMAL1. During the day, CLOCK:BMAL1 binds DNA and activates the transcription of *Period* (*Per1*, *Per2*, and *Per3*) and *Cryptochrome* (*Cry1* and *Cry2*), which upon translation form a repressive PER-CRY complex that decreases CLOCK:BMAL1 DNA binding at night and inhibits CLOCK:BMAL1-mediated transcription^23–25^. Beyond regulating core clock gene transcription, the circadian clock also initiates the rhythmic expression of thousands of genes to control daily biological processes^21,22,26^.

Accumulating evidence indicates that CLOCK:BMAL1 alone is not sufficient to generate active CREs, and that the activity of CLOCK:BMAL1-bound CREs rather depends on the cooperation between CLOCK:BMAL1 and other TFs^27–29^. CLOCK:BMAL1 has been identified as a pioneer-like transcription factor^30^. At CLOCK:BMAL1-bound CREs, nucleosome occupancy measured by MNase-Seq is rhythmic, with lower signal at the time of peak CLOCK:BMAL1 binding. A recent cryo-electron microscopy study revealed that CLOCK:BMAL1 can bind mononucleosomes when its E-box binding motif is positioned at the nucleosome entry-exit site^31^. These findings suggest that CLOCK:BMAL1 modulates chromatin dynamics by promoting nucleosome removal, thereby enhancing cooperativity with other TFs and enabling transcription of target genes.

Characterizing TF cooperative binding across entire CREs at single DNA molecule resolution has been challenging. While techniques like chromatin immunoprecipitation coupled to high-throughput sequencing (ChIP-Seq) have helped map TF binding genome-wide in diverse species and tissues, they lack the ability to assess nucleosome occupancy and TF cooperation on the same DNA molecule. MNase-seq and ATAC-seq provide base-pair resolution and can infer TF cooperation and nucleosome footprints, but they cannot directly reveal TF co-binding on single DNA molecules. The development of SMF, which is based on the foundational NOMe-seq approach^32,33^, has overcome some of these limitations by enabling detection of TF and nucleosome footprints at CREs with single-molecule resolution over a range of ∼500bp^6^. Here, we adapted SMF to 41 mouse liver CREs *in vivo*. Combined with our custom-made computational pipeline, this approach revealed that CREs *in vivo* exhibit stereotypical chromatin conformations reflecting dynamic changes in TF binding and nucleosome positioning. Further dissection of how CLOCK:BMAL1 regulates these distinct chromatin states shed light on how SMF can be leveraged to determine how TFs structurally and hierarchically remodel chromatin, evict/shift nucleosomes, and promote the recruitment of other TFs to regulate transcription.

## Results

### CREs exhibit stereotypic chromatin conformations that are conserved between samples

To characterize the modality of CLOCK:BMAL1 cooperation with other TFs, we applied SMF to liver nuclei from wild-type (WT) mice collected at ZT06 (midday) and ZT18 (midnight) and from clock-deficient *Bmal1^-/-^* mice (BMKO) at ZT06 (n = 3 per group), representing conditions exhibiting strong, weak, and no CLOCK:BMAL1 DNA binding, respectively^34,35^. SMF relies on treating nuclei with the exogenous methyltransferase M.CviPi to methylate exposed cytosines at GpC dinucleotides, followed by bisulfite or enzymatic conversion of unmodified cytosines to uracil (Fig. 1a). SMF therefore labels every GpC across the genome according to DNA protection by chromatin-associated proteins and identifies simultaneous binding of TFs and nucleosomes on single DNA molecules. We carried out SMF at a total of 41 CREs (see methods) that were PCR-amplified to increase sequencing depth to a few thousand DNA molecules and sequenced as paired-end 250 bp (PE250) reads to characterize DNA protection across full-length CREs (Fig. 1a, S1). Footprint analysis was performed using a custom bioinformatics pipeline that reconstructed PE250 reads for each CRE and animal, removed PCR duplicates, and output SMF footprint signals across CREs as binary strings with protected GpCs labeled as 1 and exposed GpCs labeled as 0 (see methods for details; Fig. 1a). Conversion efficiency of cytosines at HCH sites (H = A, C, or T) was high, with 96.81 +/- 2.02% (mean +/-SD) of cytosines sequenced as thymine (Fig. S2A).

**Figure 1.**
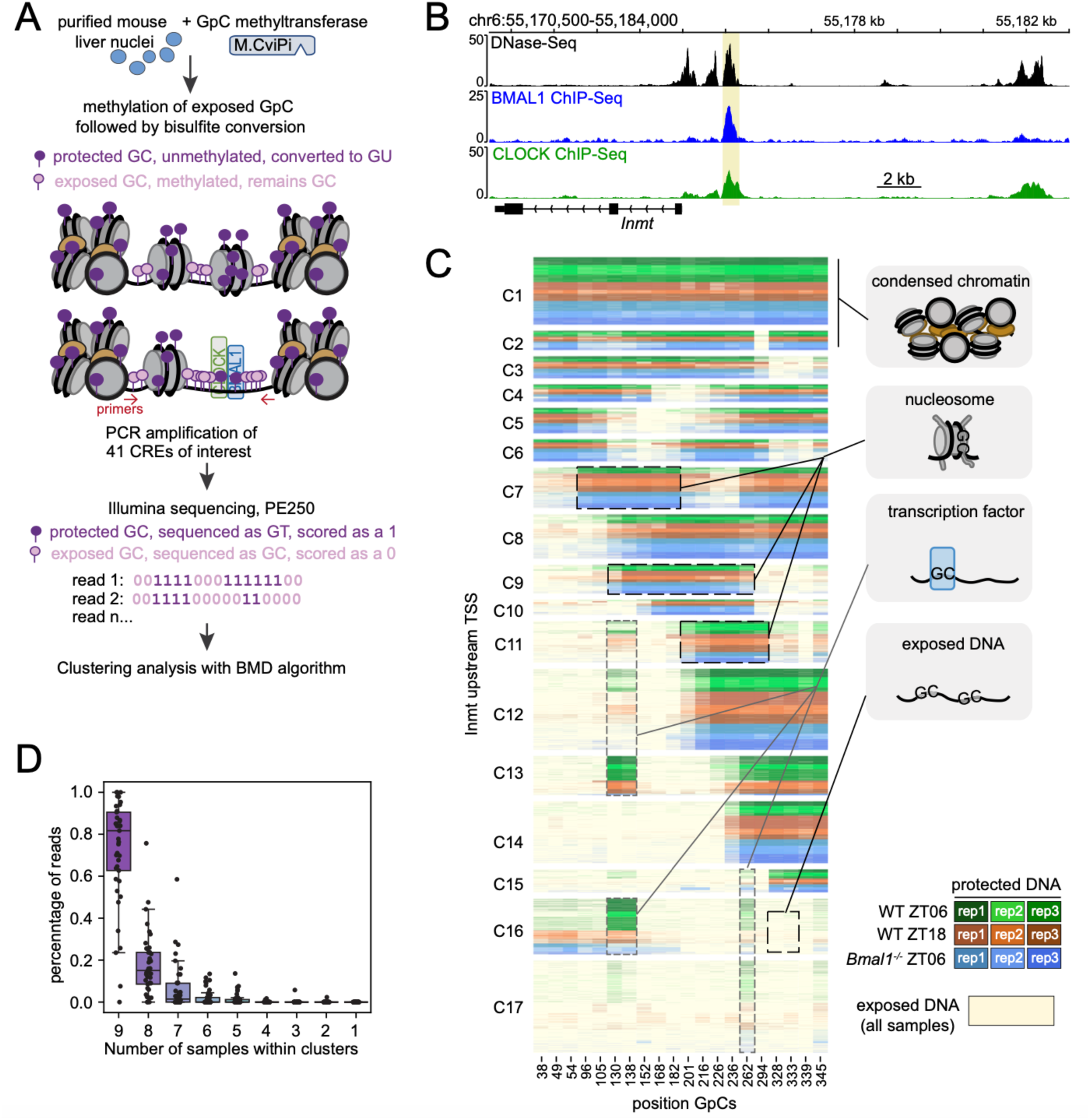
CREs exhibit stereotypic chromatin conformations. **(A)** Experimental design and bioinformatic analysis of single molecule footprinting (adapted from Sonmezer et al., 2021^6^). **(B)** DNase-Seq, CLOCK ChIP-seq and BMAL1 ChIP-seq signals at an enhancer 1kb upstream *Inmt* TSS (chr6:55152994-55153301; highlighted in yellow). **(C)** Heatmap illustrating SMF signal at the enhancer upstream *Inmt* TSS in mouse liver. Each line represents a GpC protection event on a single DNA molecule, with exposed/methylated cytosines colored in yellow, and protected/unmethylated cytosines colored in green (WT ZT06), brown (WT ZT18) or blue (BMKO ZT06). Shades of green, brown, and blue distinguish biological replicates. Reads from all nine animals (3,385 per sample) were clustered by the BMD clustering algorithm in 17 clusters, and each column illustrates protection at single GpCs (range of 308 bp). Boxes and lines illustrate four major protection events. **(D)** Percentage of reads allocated to clusters comprising reads from 1 to 9 samples for each of the 41 CREs. Dot for each boxplot represents one of the 41 CREs.

We first sought to determine if DNA protection at CREs occurs at specific sites and/or follows distinct patterns. To test this, we binned an equal number of reads per CRE from all nine biological samples to avoid overrepresentation of some samples *vs*. others, and performed clustering using the Binary Matrix Decomposition (BMD) algorithm that specializes in clustering binary data^36^. Reads within each CRE could be clustered into 11-31 chromatin states reflecting distinguishable organization of TFs and nucleosome signals (Fig. S3A). For example, reads clustered into 17 distinct states at an enhancer upstream *Inmt* transcription start site (TSS), which is targeted by CLOCK:BMAL1 and used hereafter as a prototypical CRE with accessible chromatin as assessed by DNase-seq^37^ (Fig. 1B,C). Interestingly, each cluster revealed a different protection profile. Cluster 1 (C1) showed high protection at nearly all GpCs, likely reflecting a condensed chromatin state. Clusters 2 to 8 had several regions displaying continuous protection over at least 75 bp, likely reflecting chromatin states bound by one or more nucleosomes (Fig. 1C). Considering that most TFs have less access to nucleosomal DNA than to free DNA^13^, these clusters could reflect transcriptionally inactive states. In contrast, clusters 11 to 15 showed less nucleosome protection and increased protection at one or two consecutive GpCs that likely correspond to a TF bound to DNA. When all nucleosomes are evicted, CREs can reach a fully accessible chromatin state, as in clusters 16 and 17 (Fig. 1C). Together, this clustering analysis suggests that SMF detects the multiple configurations that CREs adopt through dynamic changes in TF binding and nucleosome positioning.

Given prior evidence of decreased nucleosome occupancy upon CLOCK:BMAL1 binding^30^, we anticipated that CREs might exhibit distinct chromatin conformations in BMKO samples. However, these conformations were strikingly conserved across all nine biological samples, including between WT and BMKO (Fig. 1C). At the enhancer upstream *Inmt* TSS, all clusters contained reads from at least eight animals, and 14 clusters included reads from all nine animals (Fig. 1C, S3B-C). Thus, nucleosome positioning at CREs is not random but genetically encoded by DNA sequence and/or TF binding, as proposed elsewhere^38,39^. Similar results were found at other CREs (Fig. S3B). On average, 96.37% of all reads fell into clusters common to 7-9 animals, and clusters defined by reads from 6 or fewer samples were, for the most part, minor clusters encompassing less than 3.63% reads (Fig. 1D, Fig. S3B,D). This indicates that chromatin conformations are highly conserved among samples and largely unaffected by genotype or timepoint.

To ensure these findings were not biased by the pipeline, we performed additional analyses. First, BMD clustering of each sample individually revealed that clusters largely overlapped across animals, and hierarchical clustering grouped clusters into five major categories of nucleosome protection (Fig. S4A–B). Importantly, four of these categories contained clusters from all animals. Second, pairwise comparisons of ∼15,000 reads at enhancer upstream *Inmt* TSS (115 million comparisons total) showed no segregation by genotype or time (Fig. S4C–D). Finally, principal component analysis of all reads likewise revealed no sample-specific distributions (Fig. S4E).

Taken together, these data indicate that the chromatin states we identified represent biologically relevant chromatin conformations, that are distinguishable based on histone and TF protection, and conserved across timepoints and genotypes. Given that transcription occurs in burst^40–42^, it is tempting to speculate that the multiple chromatin conformations detected at each CRE reflect cell-to-cell heterogeneity in transcriptional activity.

### Chromatin states are conserved across tissues

To investigate whether different cell types contribute to the emergence of multiple chromatin states in the liver, we first performed single-nucleus RNA-seq (snRNA-seq) in two biological replicates using nuclei purified with the same protocol as for SMF. The analysis identified 10 distinct cell types, each expressing cell type-specific canonical marker genes (Fig. S5A-D). Consistent with previous findings^43^, hepatocytes accounted for 60-64% of captured liver cells. Because 80-90% of adult hepatocytes in C57BL/6 mice are polyploid^43^, hepatocytes are estimated to contribute to ∼80% of the liver’s DNA content^44^. Thus, the large majority of the stereotypic chromatin states identified by SMF likely originates from hepatocytes.

To explore this further, we performed SMF on nuclei from five additional tissues (kidney, lung, spleen, skeletal muscle, and cortex) across three conditions (WT ZT06, WT ZT18, and BMKO ZT06) with two biological replicates for each condition and tissue. We reasoned that if chromatin states were driven by cell-type-specific organization, different tissues should show distinct conformations. These tissues and samples differ in function, cell type composition, and CLOCK:BMAL1 binding, providing sufficient variability to test this hypothesis. Following the same experimental procedure, we carried out SMF at *Por* distal enhancer and a CRE located in *Nr1d1* 1^st^ intron, and clustered reads from all 36 samples together using BMD (one kidney WT ZT6 sample was excluded from the 36 samples for *Nr1d1* 1^st^ intron clustering due to insufficient number of reads). Surprisingly, chromatin states were highly conserved across all six tissues, with each state found in all tissues (Fig. S6A, B). Notably, nucleosome positioning patterns were remarkably consistent across tissues, with the primary difference reflected in the proportions of chromatin states and TF binding events, as exemplified by the *Por* enhancer in the spleen (Fig. S6A-B). This suggests that the chromatin states characterized in the liver are unlikely to be driven by cell-type clustering, but rather reflect an organization of nucleosomes and TF binding events that prevails in most cell types. We also note that the target genes of CREs included in this study were, with the exception of clock genes, exclusively expressed in hepatocytes, suggesting that hepatocytes may have more active states than other cells.

Taken together, these results further support the hypothesis that chromatin states are genetically encoded, either by DNA sequence and/or ubiquitous chromatin remodeler/TFs, and that cell-type or tissue-specific TF binding and cooperative events modulate the proportion of chromatin states to regulate CRE activity.

### SMF defines the multiple positions of nucleosomes at CREs

The numerous chromatin conformations that we identified by SMF clustering analysis (Fig. 1C) were largely linked to the different positions of long stretches of DNA protection that we interpreted as nucleosome protection. Because SMF-based analysis of nucleosome positioning may reveal how TFs dynamically remodel chromatin at an unprecedented resolution, we sought to determine formally whether these long stretches reflect bona fide nucleosome signal. To this end, we first developed an algorithm that computationally identified within each cluster long tracts of GpC protection as nucleosomes and calculated the percentage of nucleosome protection at each GpC for all CREs (see methods). We then compared this SMF-derived nucleosome signal with the current gold-standard for nucleosome mapping, *i.e*., sequencing of ∼150bp DNA fragments obtained from chromatin digestion with micrococcal nuclease (MNase-Seq)^45^. Using a public mouse liver MNase-seq dataset comprising nearly 1 billion nucleosomes from 47 biological replicates of WT and BMKO mice^30^, we found strong agreement, with SMF nucleosome protection signal at all 1,137 GpCs being positively correlated with MNase-seq signal (Pearson correlation coefficient: 0.74, p-value = 1.42 x 10^-200^) (Fig. 2A, S7). Importantly, SMF also resolved complex profiles of MNase-Seq signal that are difficult to interpret. For example, the lack of clear nucleosome positioning by MNase-Seq at the enhancer upstream *Inmt* TSS (Fig. 2B) can be explained by the half-dozen positions that nucleosomes occupy at that enhancer (Fig. 1C). Moreover, events where MNase-Seq indicates that dyads of two adjacent nucleosomes are spaced by less than 140 bp apart (*e.g*., as if nucleosomes were overlapping) are clarified by SMF showing well-positioned nucleosomes on distinct DNA molecules and parsed into different clusters (Fig. 2C, S8A). Thus, SMF clustering improves the resolution of nucleosome positioning across CREs and enables the quantification of the fraction of alleles comprising a nucleosome at specific positions.

**Figure 2.**
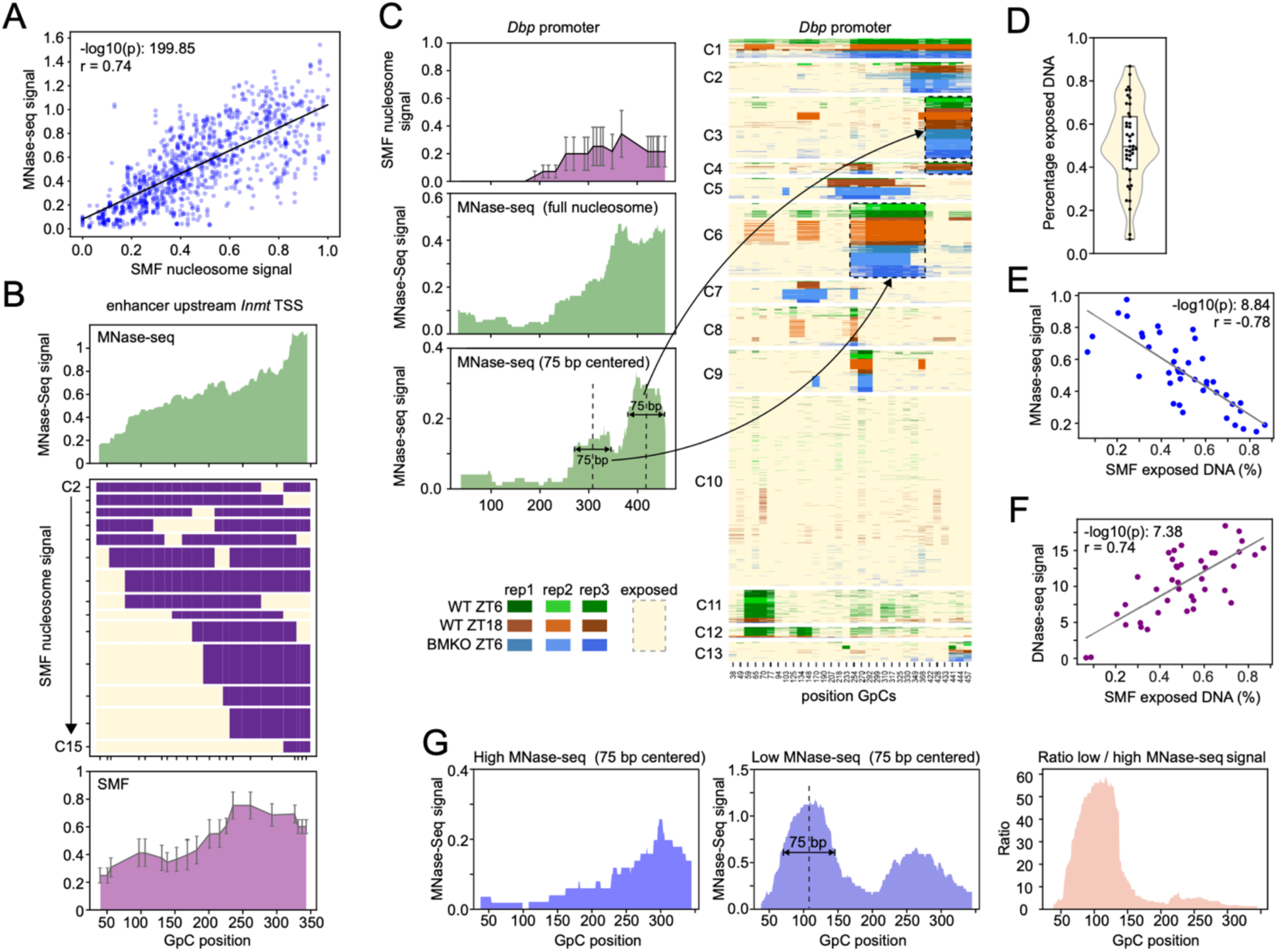
SMF improves the resolution of nucleosome positioning. **(A)** Pearson correlation between SMF nucleosome signal and MNase-seq signal at 1137 distinct GpCs located in 41 CREs. MNase-Seq datasets comprise 978,740,598 nucleosomes originating from 47 WT or BMKO mouse livers. **(B)** Nucleosome signal at *Inmt* TSS upstream enhancer. Top: MNase-seq signal; Middle: panel representing the definition of each GpC as either nucleosome (purple) or accessible chromatin (naked DNA or bound by TFs; yellow) in chromatin states C2 to C15 (see Figure 1C); Bottom: SMF nucleosome signal (equivalent to percentage of nucleosome protection at each GpC). **(C)** SMF signal at *Dbp* promoter (chr7:45354206-45354625). Top left panel: SMF nucleosome signal; Middle and bottom left panels: MNase-seq signals displayed with nucleosome signal calculated for the full-length 147 bp nucleosome (middle left) or calculated for 75 bp centered on the nucleosome dyad (bottom left; commonly used to improve the resolution of nucleosome positioning by MNase-Seq). Right: heatmap displaying SMF profile as in Figure 1C. Reads from all nine animals (1,715 reads for each sample) were clustered in 13 clusters, and each column illustrates protection at a single GpC (range of 420 bp). **(D)** Percentage of exposed DNA for each of the 41 CREs analyzed in this study, and calculated using SMF profiles and the computationally-defined nucleosome position. **(E)** Pearson correlation between SMF exposed DNA signal and MNase-seq signal using signals at 41 CREs. **(F)** Pearson correlation between SMF exposed DNA signal and DNase-seq signal using signals at 41 CREs. **(G)** MNase-seq nucleosome signals at *Inmt* TSS upstream enhancer, obtained from paired-end sequencing of high (left) and low (middle) MNase digested mouse liver nuclei. The ratio of low over high MNase-seq signal (right) was calculated to represent nucleosome fragility at *Inmt* TSS upstream enhancer.

Because nucleosomes represent a physical barrier for most TF binding^13^, we next leveraged our data to determine the percentage of exposed DNA at each CRE. We calculated the proportion of exposed DNA per cluster, which was then aggregated for each CRE (Fig. S8B). This analysis revealed that the extent to which DNA is exposed at CREs varied extensively. Excluding two CREs with little DNase-seq peak/signal and containing more than 90% of nucleosomal DNA (used as controls), nucleosomal DNA proportions ranged from as low as 13.2% to as high as 79.5% across CREs (Fig. 2D). Such variability implies CRE-specific regulation of transcriptional activity with, for example, increased dependence on chromatin remodelers at CREs with less exposed DNA. As expected, the percentage of exposed DNA at each CRE was negatively correlated with MNase-Seq signal (Fig. 2E) and positively correlated with DNase-seq signal (Fig. 2F).

Nucleosomal DNA can partially unwrap from the histone octamer, and when this transient conformation is stabilized, nucleosomes become more susceptible to MNase digestion and are called fragile nucleosomes^46^. CREs are enriched in fragile nucleosomes, and fragility can be increased without histone eviction in response to environmental changes that poise genes for activation^47,48^. Given that SMF detects discrete chromatin states and resolves nucleosome positioning, we asked whether fragile nucleosomes associate with specific chromatin states by analyzing public high and low MNase-Seq datasets from mouse liver^49^. At the enhancer upstream *Inmt* TSS, we observed a peak present in low but not high MNase-Seq datasets, indicating a fragile nucleosome at that location (dyad located ∼105bp downstream of the 5’ end of SMF reads) (Fig. 2G). Examination of DNA protection (Fig. 1C) revealed a nucleosome at position 38-168 bp in cluster C4, matching the low MNase-Seq peak at position ∼50-150bp (Fig. 2G). Analyses of other CREs (Fig. S8C-E) confirmed that combining SMF with low vs. high MNase-Seq can uncover chromatin states enriched for fragile nucleosomes, suggesting that the complementary usage of these assays could shed light on the mechanisms underlying the formation of fragile nucleosomes.

### Protection at E-boxes is variable between CREs

CLOCK:BMAL1 binds DNA rhythmically, yet it remains unclear whether other TFs bind the same E-boxes when CLOCK:BMAL1 DNA affinity is minimal at night. To address this, we analyzed the 30 CREs showing rhythmic BMAL1 ChIP-Seq signal among the 41 CREs studied (Fig. S9A). Specifically, we quantified DNA protection at all canonical and degenerate E-boxes (up to one mismatch to CACGTG motif) when at least one GCH was located within 5 bp of the motif (CLOCK:BMAL1 footprint on DNA is ∼18 bp^28,50^; Fig. S9B), reasoning that binding of CLOCK:BMAL1 and/or other TFs to E-boxes should prevent GpC methylation by M.CviPi (sequences in Table S1). We calculated protection at 158 GCH sites representing 107 E-boxes, using reads where E-boxes were exposed and not protected by a nucleosome. Protection was significantly higher in WT ZT06 and lower in BMKO, consistent with CLOCK:BMAL1 DNA binding (Fig. S9C).

At the level of individual E-boxes, variability was substantial. Among 107 E-boxes analyzed, 75 carried the CANNTG sequence preferred by bHLH TFs^51^. Of these, 24 had greater protection in WT ZT06 (35 GpCs located in 14 CREs; Fig. 3A), a proportion higher than at non-CANNTG motifs (2/32; Fig. S9D). Similar trends were observed at additional E-boxes (Fig. 3A, S9D), though some did not reach significance likely due to variability between samples. Many GpCs at the 24 significantly different E-boxes showed minimal protection in BMKO mice. For example, 11/35 GpCs had <10% protection in BMKO mice, and BMKO ZT06 had the lowest protection for 32/35 GpCs, indicating that CLOCK:BMAL1 is primarily responsible for protecting these sites (Fig. 3A, S9C-D).

**Figure 3.**
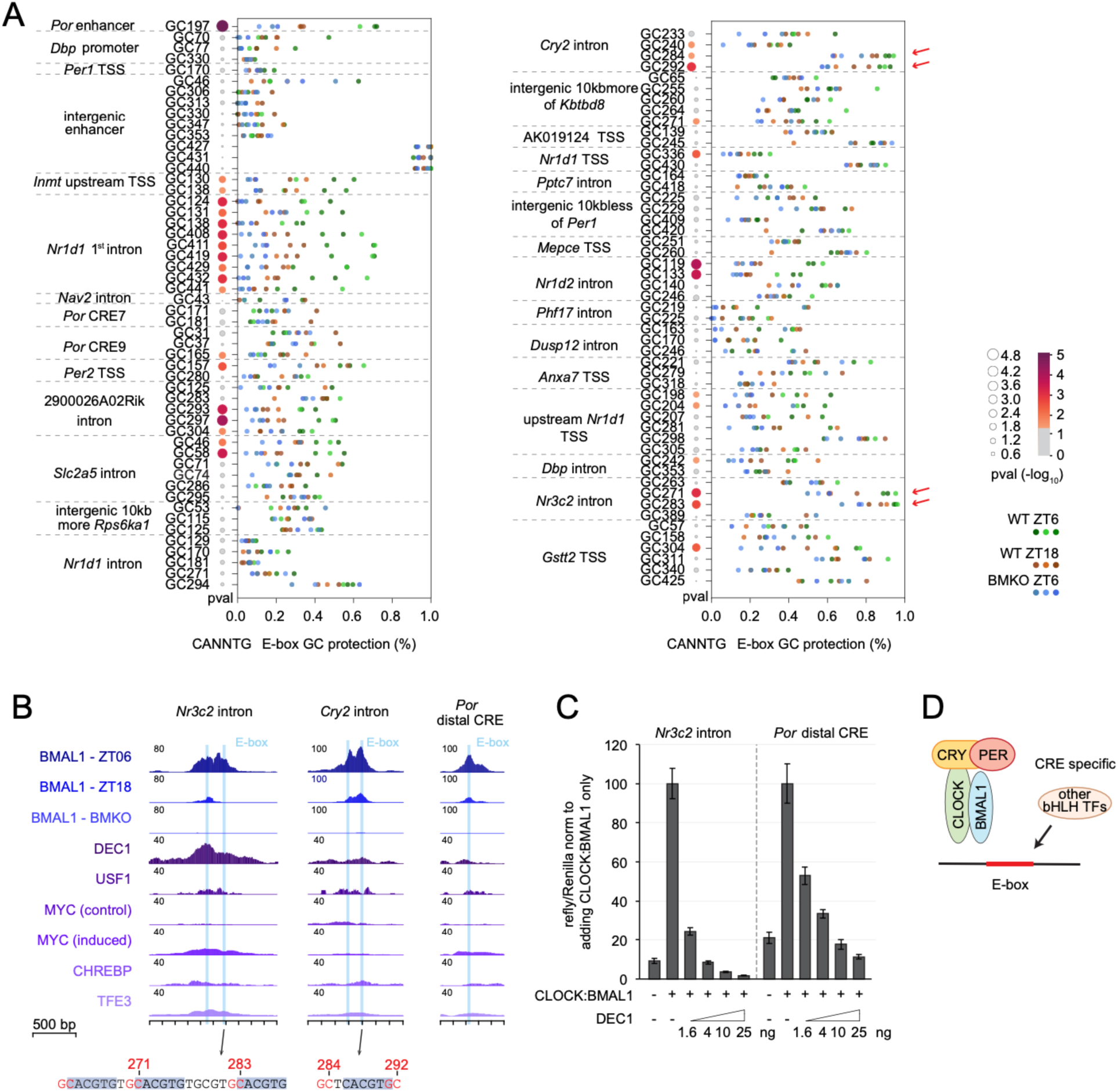
Patterns of protection at E-boxes differ between CREs. **(A)** Scatter plot of the percentage of protection at the 103 E-box GpCs for all 9 samples, using Fig. 1C color code. The size and color of the bubble plot (left) depict the p-value obtained by one-way ANOVA (n = 3 samples/group). Circles are colored in grey if p>0.05. The four red arrows point to GpCs with p-value &<= 0.05 and that retained high protection in BMKO mice. **(B)** ChIP-seq tracks for BMAL1 and five additional bHLH TFs. Blue shadings mark E-box motifs. Black arrows point to the positions of E-boxes and GpCs corresponding to the red arrows in panel A. **(C)** Repression assay of CLOCK:BMAL1 by DEC1 using reporters containing *Nr3c2* intron or *Por* distal CRE. HEK293T cells were co-transfected with *Clock* and *Bmal1* expression vectors (50 ng each) and increasing amounts of *Dec1* expression vector (0, 1.6, 4, 10, and 25 ng). The empty pGL3-promoter vector (pGL3) served as a negative control. Firefly/renilla luciferase activity of each condition and each replicate was normalized to the mean. Error bars represent the S.E.M. of three biological replicates. Each biological replicate corresponds to the mean of two technical replicates. **(D)** Schematic depicting how other bHLH TFs compete with CLOCK:BMAL1 when its binding is inhibited.

Notable exceptions were seen at two CREs located in *Cry2* and *Nr3c2* introns, where E-box protection exceeded 50% in BMKO, suggesting that other bHLH TFs bind when CLOCK:BMAL1 DNA binding is low (Fig. 3A-B; red arrows). Analysis of public mouse liver ChIP-seq datasets for five bHLH TFs other than CLOCK:BMAL1 showed strong DEC1 signal at *Nr3c2* intron, supporting this possibility (Fig. 3B). To test whether DEC1 competes with CLOCK:BMAL1 at *Nr3c2* intron, we performed a luciferase-based dose-response competition reporter assay in HEK293T cells. While CLOCK:BMAL1 alone strongly increased *Nr3c2* intron-driven luciferase activity, DEC1 potently repressed reporter activity even at low plasmid concentration (Fig. 3C). By contrast, the *Por* distal CRE, a negative control site with low protection in BMKO mice and no DEC1 ChIP-seq signal, required a ∼6-fold higher DEC1 for comparable repression. These results therefore indicate that the *Nr3c2* intron is more sensitive to DEC1 than the *Por* distal enhancer, consistent with SMF-detected competition event between CLOCK:BMAL1 and other bHLH TFs. Although strong binding was not detected for the five TFs tested at *Cry2* intron, its high protection in BMKO suggests targeting by another member of the bHLH TF family, which comprises over 100 TFs^51^.

Some CREs also showed similar levels of protection between groups, or even higher protection in WT ZT18. For example, at an intergenic CRE, three GpCs at E-box 3 showed 80% protection in BMKO ZT06 (GpCs 427, 431, 440; Fig. 3A, S9E). Given that open chromatin states at this CRE are enriched in WT ZT06 (Fig. S9E-F; see below), this suggests that CLOCK:BMAL1 binds DNA during the day, and other TFs compete with CLOCK:BMAL1 for E-box binding especially when CLOCK:BMAL1 affinity decreases. Based on public ChIP-Seq datasets, these TFs may include USF1, DEC1, or TFE3 (Fig. S9G).

In summary, SMF uncovered a wide range of protection patterns at CLOCK:BMAL1-bound E-boxes. Many sites showed protection profiles that directly mirror CLOCK:BMAL1 binding strength, dropping to near-background levels in BMKO mice indicative of exclusive CLOCK:BMAL1 occupancy. In contrast, other E-boxes retained partial or even full protection without CLOCK:BMAL1. These findings highlight that other bHLH TFs can compete with CLOCK:BMAL1 for E-box access, but crucially, this competition is CRE-specific (Fig. 3D).

### Cooperative DNA binding between CLOCK:BMAL1 molecules across >250bp

The resolution of SMF also enabled examination of cooperative CLOCK:BMAL1 binding between multiple E-boxes. We focused on a CRE located in the first intron of the clock gene *Nr1d1* that contains 5 E-boxes (Fig. 4A). E-boxes 1 and 2 are separated by 6 bp, forming a dual E-box considered as a preferred CLOCK:BMAL1 binding site^52^. Intriguingly, we observed CLOCK:BMAL1 binding on E-box 2 but not E-box 1, and protection at E-box 2 was instead associated with protection at E-box 3, which are separated by 9 bp (Fig. 4A). Protection at E-boxes 4 and 5, which are 16 bp apart, was also strongly correlated (Fig. 4A). To quantify cooperative binding, we adapted a published approach that computes a “normalized extent of co-binding” (N.EOC) between protection events at two GpCs^5^ (see Fig. S10A and methods for details). This analysis confirmed strong CLOCK:BMAL1 cooperative binding between E-boxes 2/3 and 4/5 (Fig. 4B). Cooperation at E-boxes 2/3 occurred only in WT ZT06, whereas cooperation at E-boxes 4/5 was similar in both WT ZT06 and WT ZT18. Examination of bHLH TF ChIP-Seq datasets revealed strong USF1 signal at E-boxes 4/5 (Fig. S10B), potentially accounting for their protection in WT ZT18 (Fig. 3A). Notably, we also observed a higher N.EOC between the last four E-boxes, *i.e.*, between GpCs 124/131 (E-boxes 2/3), GpCs 411/419 (E-box 4) and GpCs 429/432/441 (E-box5), suggestive of a cooperative binding of CLOCK:BMAL1 over ∼280 bp (Fig. 4B). Importantly, this cooperative binding was unique to WT ZT06. A second multi-E-box CRE in the *Dbp* promoter similarly showed cooperative binding across >250 bp, again stronger in WT ZT06 (Fig. S10C).

**Figure 4.**
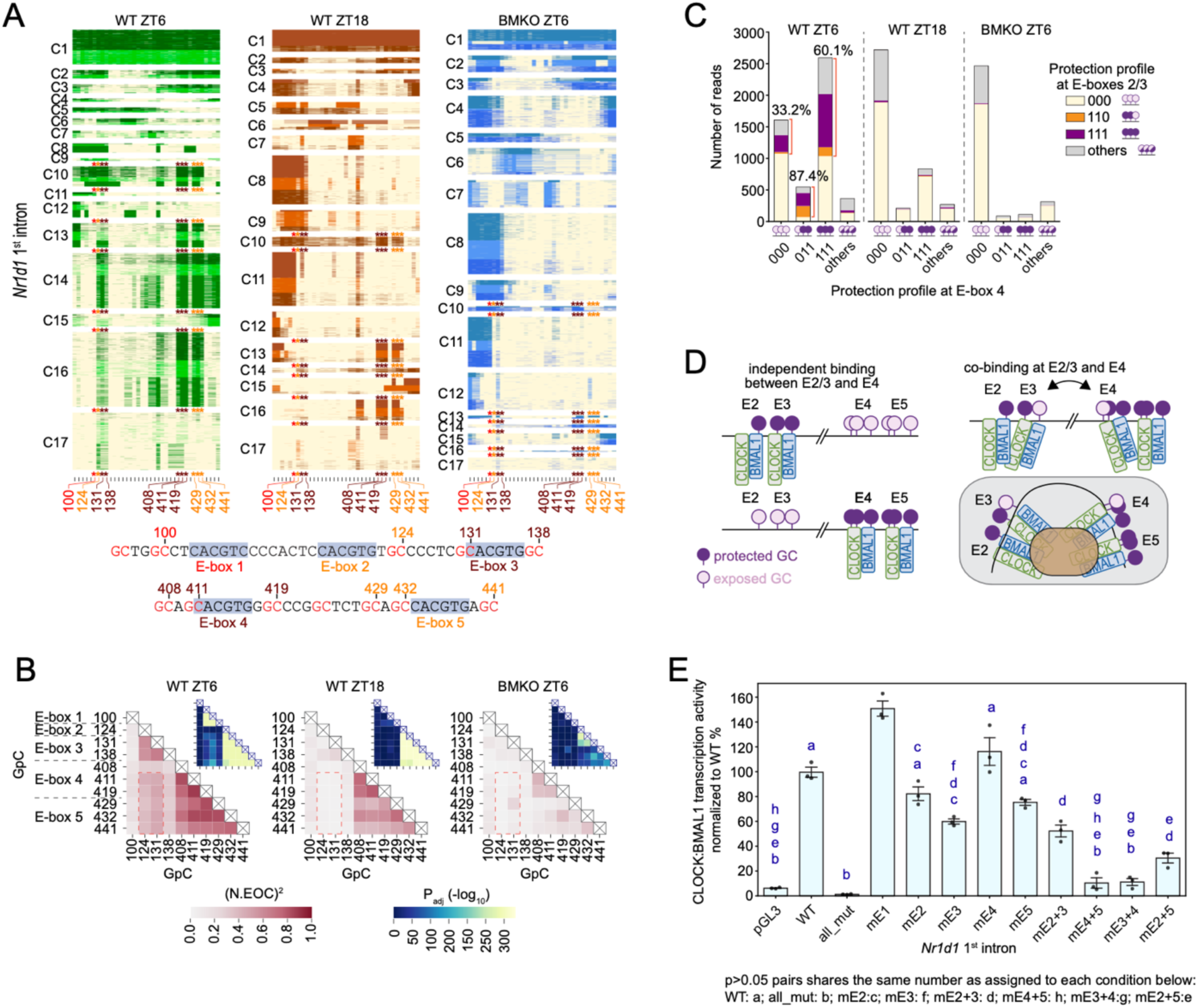
Long-range cooperative CLOCK:BMAL1 binding *in vivo*. **(A)** SMF profile at an enhancer located in *Nr1d1* 1^st^ intron (chr11:98664610-98665043) in mouse liver. The heatmap displays protection from GpC methylation as in Fig. 1C (n = 2,368 reads for each sample; range of 434 bp). Asterisks highlight GpCs at five E-boxes, with sequences provided below the heatmaps. **(B)** Normalized extend of co-binding (N.EOC) calculated between every GpC located within 5 bp of an E-box at *Nr1d1* 1^st^ intron. Fisher’s exact test followed by Benjamini-Hochberg correction. Squared N.EOC values were plotted in heatmap. **(C)** Number of reads based on the protection profiles at E-box4 and E-box 2/3 for each group. Protection events at E-box 4 (GpCs 408/411/419) were separated in 4 categories: all GpCs exposed (000), only GpC 408 exposed (011), all GpCs protected (111), and all other protection events (others). Protection events at E-boxes 2/3 (GpCs 124/131/138) were similarly separated in 4 categories: all exposed (000, yellow), only exposed at GpC 138 (110, orange), all protected (111, purple), and others (grey). **(D)** Schematic illustrating changes in GpC protection at E-box2/3/4/5 based on CLOCK:BMAL1 DNA binding. **(E)** Luciferase reporter assays using a CRE in *Nr1d1* 1st intron or variants containing combinations of E-box mutations. mE1, mE2, mE3, mE4, and mE5 indicate reporters with mutations at E-box 1, 2, 3, 4, and 5, respectively. Similarly, mE2+3 indicate a reporter with both E-boxes 2 and 3 mutated. pGL3 served as a negative control. Error bars represent the S.E.M. of three biological replicates, with each biological replicate corresponding to the mean of two technical replicates. One-way ANOVA between the 12 groups (p &<= 0.05) was performed and followed by a post-hoc test using Tukey’s Honestly Significant Difference (HSD) test (FWER=0.05). Pairs with the same letter were not significant (p > 0.05).

Closer inspection of *Nr1d1* first intron CRE revealed that although CLOCK:BMAL1 protects GpC 138 (E-box 3) and GpC 408 (E-box 4), their direct co-binding was much lower than co-binding between GpCs 124/131/411/419 (Fig. 4B). We hypothesized that CLOCK:BMAL1 may bind differently when engaged in long-range interactions. To test this, we categorized protection at E-box 4 (GpCs 408/411/419) in four classes: all GpCs exposed (000), only GpC 408 exposed (011), all GpCs protected (111), and all other protection events (others) (Fig. 4C). We then counted, for each E-box 4 class, reads exhibiting specific patterns at E-boxes 2/3 (GpCs 124/131/138): all exposed (000, yellow), only GpC 138 exposed (110, orange), all protected (111, purple), and others (grey). At ZT06, when CLOCK:BMAL1 is high, protection at E-boxes 2/3 was twice as frequent when E-box 4 was protected vs. unprotected (60.1% vs. 33.2%), confirming long-range cooperativity across 280 bp. Strikingly, the proportion was even higher when GpC 408 was exposed (87.4% vs. 60.1% of reads), with many reads showing exposure of GpC 138 at E-box 3, suggesting that CLOCK:BMAL1 binds GpC138 and GpC408 differently when engaged in a long-range interaction (Fig. 4C). At ZT18 and in BMKO mice, protection at E-boxes 2/3 was equivalent or lower when E-box4 was protected. Thus, CLOCK:BMAL1 cooperative binding between E-boxes 2/3/4/5 was only enriched during the day, and may be mediated by proteins that specifically interact with CLOCK:BMAL1 when DNA binding affinity is high (Fig. 4D).

To test whether this long-range cooperativity affects CRE activity, we expressed CLOCK and BMAL1 in HEK293T cells and measured luciferase activity using reporter constructs driven by the endogenous *Nr1d1* 1^st^ intron CRE or variants bearing different combinations of E-box mutations (Fig. 4E). CLOCK:BMAL1 induced strong luciferase activity that was completely blunted when all 5 E-boxes were mutated, indicating effective mutations without creation of new TF binding sites. Single E-box mutations had modest effect on luciferase activity, with only E-box 3 mutation leading to a significant decrease by about 40%. Mutation of the dual E-boxes 4/5 substantially decreased luciferase activity by ∼90%, a far larger effect than mutating either alone. By contrast, mutation of the dual E-boxes 2/3 produced a decrease similar to mutating E-box 3 alone. This difference in dual E-boxes cooperativity is unclear and warrants further investigation. Importantly, mutations of E-boxes separated by >250 bp (mutation of E-boxes 3 and 4, or of E-boxes 2 and 5) caused a much stronger reduction in luciferase activity than the additive effect of single mutations, demonstrating a synergistic effect (Fig. 4E). These results thus support SMF findings and demonstrate that CLOCK:BMAL1 cooperative binding can extend over distances exceeding 280bp, and that it substantially impacts CLOCK:BMAL1-mediated transcription.

In summary, CLOCK:BMAL1 not only engages in short-range dual E-boxes cooperativity but also in cooperativity across E-boxes separated by >280 bp *in vivo* to drive transcription. Because this longer distance cooperativity only occurs during the day and may involve site-specific DNA configuration changes, it likely relies on additional mediators, thus revealing a previously unrecognized mode of CLOCK:BMAL1 regulation.

### CLOCK:BMAL1 promotes rhythmic chromatin opening and cooperation with other TFs

A recent study demonstrated that CLOCK:BMAL1 binds the entry/exit site of nucleosomes, specifically when E-box(es) are located at superhelix locations +/- 5 to +/- 7 (Fig. 5A)^47^. To assess whether CLOCK:BMAL1 nucleosome binding is prevalent and contributes to chromatin remodeling, we analyzed our SMF data at 30 CLOCK:BMAL1-bound CREs and computationally identified 66 nucleosomes having an E-box located within 20 bp of the SMF-defined nucleosome edge (Fig. 5B,C, S11A; see methods for details). Of these, 20 contained a CANNTG motif and showed higher CLOCK:BMAL1 protections in WT ZT06. Among these, the SMF nucleosome signal for 11 nucleosomes was lower in WT ZT06 when CLOCK:BMAL1 DNA binding peaked. In the remaining 46 nucleosomes, a similar trend was observed, with lower SMF nucleosome signal at peak time CLOCK:BMAL1 DNA binding for 11 nucleosomes. Consistent with CLOCK:BMAL1-mediated nucleosome removal, many CLOCK:BMAL1-bound CREs exhibited higher DNA exposure in WT ZT06 than in WT ZT18 and BMKO (Fig. 5D, S11B). Importantly, such chromatin opening was not prevalent at non-CLOCK:BMAL1 CREs (Fig. 5D, S11B). Thus, CLOCK:BMAL1 binding at nucleosome entry/exit sites occurs at many CREs, promotes nucleosome removal and creates a more accessible chromatin environment.

**Figure 5.**
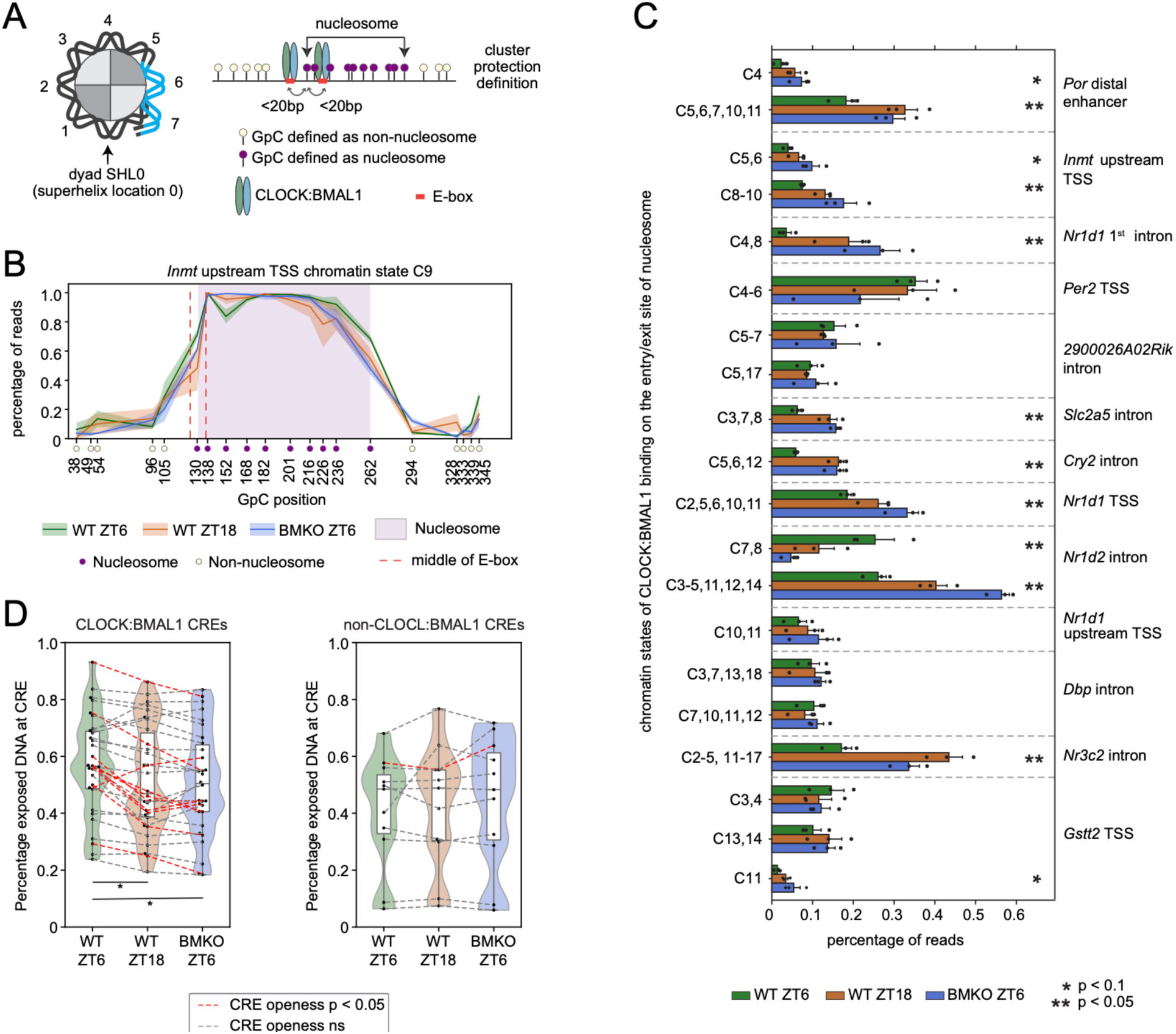
Many nucleosomes with an E-box at the entry/exit site are removed upon CLOCK:BMAL1 binding. **(A)** Schematic of CLOCK:BMAL1 preferential binding to nucleosome entry/exist site (left) and approach used to uncover E-box(es) at nucleosome entry/exit site. SHL = superhelix location. **(B)** Percentage of GpC protection at *Inmt* TSS upstream enhancer, chromatin state C9 (see Fig. 1C), showing a nucleosome (purple region; GpCs 130-262) with an E-box (red lines) at the entry/exist site. Shaded areas in green, brown, and blue represent the S.E.M. of 3 biological replicates. **(C)** Percentage of reads in clusters identified as having a nucleosome with CANNTG motif(s) at the entry/exist site, and where TF protection at E-box GpCs was significant between the three groups. Error bars correspond to the S.E.M. of 3 biological replicates. One-way ANOVA; * p&<0.1; ** p&<0.05. Clusters with nucleosomes at similar locations were combined. **(D)** Percentage of exposed DNA at CREs in WT ZT06, WT ZT18, and BMKO ZT06. CREs were separated based on CLOCK:BMAL1 binding by ChIP-seq. Repeated measures ANOVA between the three groups resulted in p&<0.05 for CLOCK:BMAL1 CREs but not non-CLOCK:BMAL1 CREs. * for groups between-subject pairwise T-test(s) p &< 0.05. One-way ANOVA between three groups of each CRE; p&<0.05, red lines; p>0.05, grey lines.

We next asked whether CLOCK:BMAL1-mediated increase in DNA accessibility enables other TFs to bind DNA. The *Por* distal enhancer and *Inmt* upstream TSS enhancer both contain dual E-boxes that exhibited higher protection in WT ZT06, consistent with BMAL1 ChIP-Seq (Fig. 3A, S3C). In specific chromatin states, both dual E-boxes were positioned at a nucleosome entry/exit site that showed lower occupancy at peak CLOCK:BMAL1 binding in WT ZT06. Interestingly, for both CREs, CLOCK:BMAL1 DNA binding was associated with nucleosomes progressively shifting to the right of the CREs (Fig. 6A, S3C). This increase in chromatin accessibility coincided with additional TF protection at motifs for TFs of the RFX family for both CREs (Fig. 6A, S11C). At *Por* distal enhancer, the percentage of reads exhibiting co-protection at E-box and RFX motifs (co-protection of GpCs 197, 301, and 307) was significantly higher in WT ZT06 (Fig. 6B), and the extent of TF co-binding on the same DNA molecules was higher in WT than in BMKO (Fig. 6C). Transcription of *Por* was rhythmic, peaking a few hours after the maximal CLOCK:BMAL1 DNA binding at ZT10 (Fig. 6D). At *Inmt* upstream TSS enhancer, we found that the extent of co-binding between CLOCK:BMAL1 and the protection event at RFX motif (GpC 262) was also higher in WT ZT06 than in the two other groups, corresponding to peak of *Inmt* pre-mRNA level (Fig. S11D, E). Additionally, RFX motif protection was significantly higher in WT ZT06 at *Inmt* upstream TSS (Fig. S11F), but not at *Por* distal enhancer (Fig. S11G).

**Figure 6.**
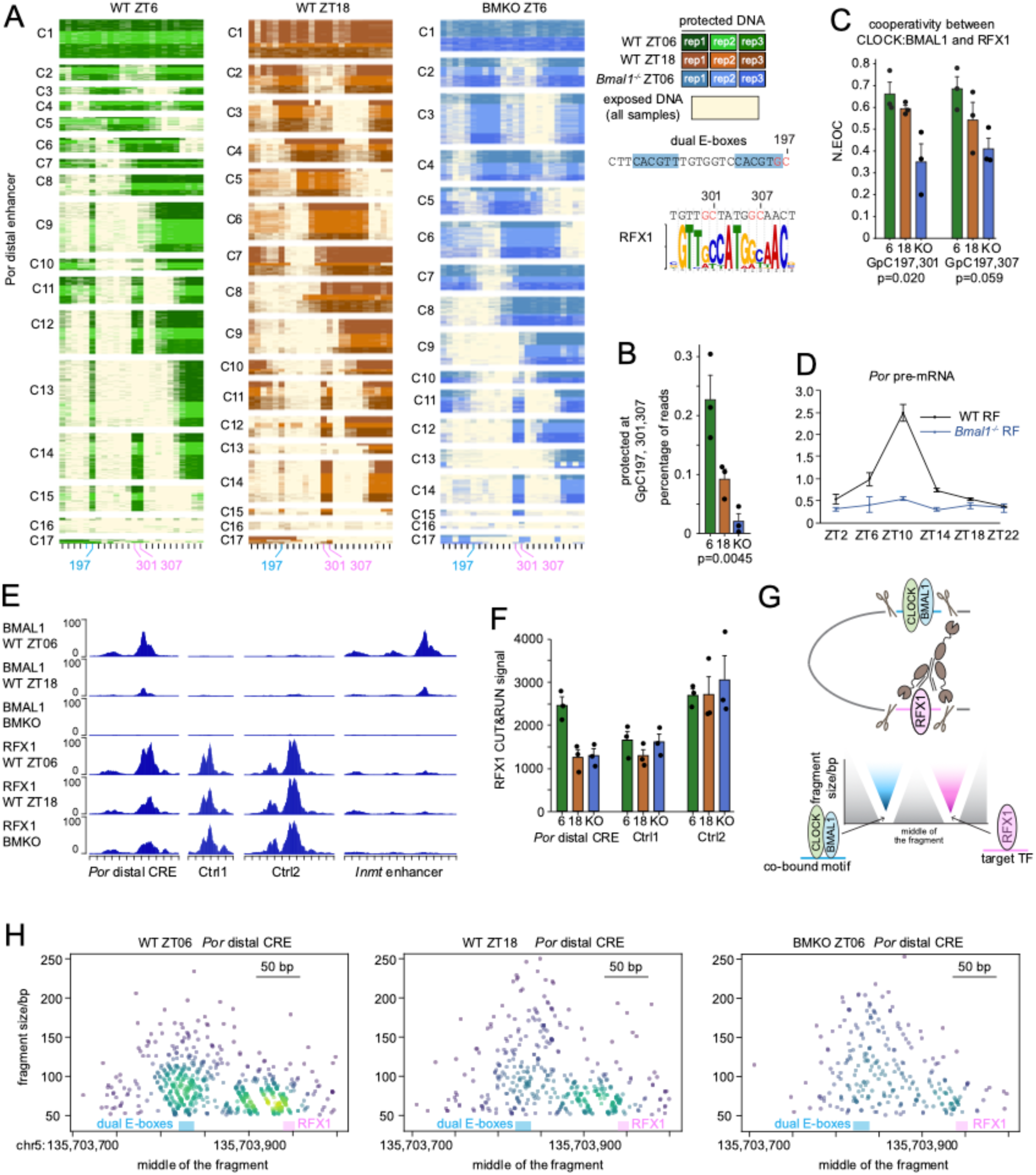
CLOCK:BMAL1-mediated chromatin opening facilitates the nearby binding of other TFs. **(A)** SMF profile at *Por* distal enhancer (chr5:135703727-135704053) in mouse liver parsed by each group (WT ZT06, WT ZT18, and BMKO ZT06). Each heatmap contains reads from three biological replicates, which are represented by the different shades of green, brown, and blue (1,833 reads in each sample). Color code is as in Figure 1C. GpC 197 reflects CLOCK:BMAL1 protection at the dual E-boxes. The consensus motif of a TF (RFX1) that may protect GpC 301/307 was found with TomTom tool using the sequence provided on the right of the heatmap. **(B)** Percentage of reads protected at the E-box and RFX1 motif (GpCs197, 301, and 307), in WT ZT06 (green), WT ZT18 (brown), and BMKO ZT06 (blue). **(C)** Normalized extend of co-binding between CLOCK:BMAL1 and the TF protection event at GpC 301/307 at *Por* distal enhancer in WT ZT06, WT ZT18 and BMKO ZT06. Each dot represents the value of each sample, and the error bars correspond to the SEM of 3 biological replicates. One-way ANOVA was performed between the three groups. **(D)** Mouse liver *Por* pre-mRNA level across the 24-hour day, from public mouse liver total RNA-Seq datasets (GSE73554) from Atger et al., 2015^81^. Black and blue line represent nighttime-fed WT and BMKO mice, respectively. **(E)** BMAL1 and RFX1 CUT&RUN signals at *Por* distal CRE, *Inmt* upstream TSS and two controls CREs without CLOCK:BMAL1 binding, in WT ZT06 (green), WT ZT18 (brown) and BMKO ZT06 (blue). **(F)** Quantification of RFX1 signals at *Por* distal CRE and the 2 control CREs from panel B. Error bars correspond to the SEM of 3 biological replicates. **(G)** Schematic of how CLOCK:BMAL1 and RFX1 co-binding would prevent cuts at E-boxes in RFX1 CUT&RUN experiments, and how it would be displayed by V-plot. **(H)** V-plots of RFX1 CUT&RUN at *Por* distal CRE in WT ZT06, WT ZT18, and BMKO ZT06. The middle of the chromosomal location of fragments is displayed against fragment size for each read.

To directly test whether CLOCK:BMAL1-mediated chromatin opening facilitates other TF binding, we performed CUT&RUN in the same three groups for BMAL1 and RFX1, an RFX family member expressed in mouse liver. As expected, BMAL1 CUT&RUN signal was strongest in WT ZT06, weaker at ZT18, and absent in BMKO mice. Consistent with the motif analysis, RFX1 CUT&RUN data showed a strong signal at *Por* distal enhancer. Strikingly, this signal was much higher in WT ZT06 than in WT ZT18 and BMKO ZT6, but not at other control CREs lacking CLOCK:BMAL1 binding, thereby further supporting that CLOCK:BMAL1 promotes RFX1 binding at *Por* distal CRE (Fig. 6E, F).

Analysis of MNase cuts, which are prevented by TF DNA binding, has been used to infer TF cooperation via V-plots that organize paired-end sequencing reads by genomic location and fragment length^5,53^ (Fig. 6G). At *Por* distal CRE, RFX1 CUT&RUN fragments displayed strong pattern differences around the E-box motif (Fig. 6G, H). In WT ZT06, RFX1 CUT&RUN cuts were almost exclusively located outside the dual E-boxes, whereas in BMKO ZT06 and even WT ZT18, cuts appeared more randomly distributed, i.e., MNase was able to cut within the dual E-boxes. These data therefore further support co-binding of RFX1 and CLOCK:BMAL1, and that this co-binding increases RFX1 binding in WT ZT06. RFX1 CUT&RUN signal was not detected at the enhancer upstream *of Inmt* TSS, likely reflecting binding by another TF.

Together, these data indicate that CLOCK:BMAL1 promotes rhythmic chromatin accessibility at many CREs, which can facilitate cooperative binding by other TFs and be associated with transcription activation.

## Discussion

By applying SMF to mouse liver *in vivo*, we demonstrate that CREs adopt a defined set of chromatin conformations that reflect stereotypic organization of nucleosomes and TFs. These chromatin states are remarkably conserved between individuals, strongly suggesting that they are genetically encoded. Carrying out SMF at CREs targeted by the pioneer-like circadian TF CLOCK:BMAL1 revealed that changes in CLOCK:BMAL1 DNA affinity did not generate activity-specific chromatin states, but rather altered their relative proportion. Specifically, at times of strong CLOCK:BMAL1 DNA binding, signal for nucleosomes with E-boxes at their entry-exit sites was decreased and chromatin states with greater accessibility were enriched. This increased accessibility coincided at some CREs with stronger footprints for other TFs and higher TF co-binding on the same molecules. Collectively, our data thus support a model wherein CLOCK:BMAL1 rhythmic DNA binding redistributes preconfigured chromatin states over the 24-hour cycle, thereby reorganizing local chromatin to prime daily rhythms in gene expression.

The remarkable conservation of chromatin states across genotypes, individuals, and tissues is striking. These states are unlikely to reflect differences among cell types, as hepatocytes contribute the majority of liver DNA (>80%) and the same states were conserved across multiple tissues with distinct biological functions. CREs remained open in BMKO mice and retained the same chromatin conformations than in WT mice, indicating that the absence of CLOCK:BMAL1 is insufficient to disrupt nucleosome positioning, and that other pioneer TFs or chromatin remodelers likely contribute to chromatin organization at CREs. These other factors are likely responsible for chromatin opening at any time of day and operate in parallel to CLOCK:BMAL1, since the loss of BMAL1 did not alter the chromatin opening. The functional significance of these stereotyped chromatin conformations remains unclear. Our clustering analysis showed that state identity was strongly influenced by nucleosome positioning across CREs. Although nucleosomes positions varied, their transitions occurred in reproducible and conserved patterns across tissues, suggesting they are genetically encoded. Nucleosome positioning has long been recognized as non-random, but the determinants of *in vivo* patterns remain debated. DNA sequence has been proposed as a major driver^54^. Yet, subsequent studies challenged this view, suggesting that sequence exerts only a modest influence and that chromatin remodelers and other DNA-binding proteins also play key roles^55–58^. Nucleosome positioning likely emerges from a complex interplay between multiple factors, *i.e*., the dynamic changes result in multiple states, and DNA sequence at CREs either limits the position of nucleosomes to specific locations, instructs where TFs bind to remodel chromatin, and/or a combination of both. Defining how these factors combine to drive transitions between chromatin states remains an important area for future investigation.

Transcription occurs in stochastic bursts interspersed with refractory periods of inactivity^40–42^. We propose that the different chromatin conformations identified with SMF reflect different biologically relevant states that represent various stages during transcriptional bursting, and that differ between different single cells or alleles. It remains unclear how chromatin states dynamically transition from one to another. Live-cell imaging study revealed substantial heterogeneity in transcriptional bursting between cells and even alleles within the same cell: some cells exhibited multiple bursts over several hours, whereas others displayed only a single burst or remained in a prolonged transcriptionally repressed state^59^. This raises the question of whether some cells are less dynamic, transitioning only between a few states, while active cells may experience most of the states. Future studies aiming at characterizing markers enriched in a specific state (e.g., active markers like Pol II, H3K27ac, and/or incorporation of histone variants that facilitate nucleosome destabilization) will help associating chromatin configuration with transcriptional dynamics.

CLOCK:BMAL1 binds DNA rhythmically across tissues, yet the extent to which other bHLH TFs bind the same E-boxes at times of low CLOCK:BMAL1 DNA binding affinity remains largely unknown. A few TFs including USF1^60^, DEC1/2 (aka BHLHE40/41)^61^, and MYC^62^ are known to compete with CLOCK:BMAL1 for E-box access, but competition may be more widespread since the mammalian bHLH TF family consists of over 100 members ^51,63^. So far, this question has been technically challenging to address, especially *in vivo,* where DNA is chromatinized. Using SMF at 30 CREs targeted by CLOCK:BMAL1, we found that levels of E-box protection varied between CREs. At 23 E-boxes located on 12 CREs, E-box protection was minimal in BMKO mice and mirrored CLOCK:BMAL1 ChIP-Seq signal, indicating that CLOCK:BMAL1 is the only bHLH TF member to target these E-boxes. In contrast, two CREs retained substantial protection in BMKO mice despite high protection in WT ZT6. This finding was also supported by the strong DEC1 ChIP-seq signal and the enhanced sensitivity to DEC1 repression at *Nr3c2* intron. Levels of protection at other E-boxes were more even between groups, including at an intergenic enhancer exhibiting very high protection even in BMKO mice. Given the overrepresentation of open chromatin states at that enhancer when CLOCK:BMAL1 ChIP-Seq is maximal, this suggests that CLOCK:BMAL1 binds DNA during the day and that other bHLH TFs bind at night. More generally, our data demonstrate that some (but not all) CLOCK:BMAL1-bound E-boxes are targeted by other TFs, and that the binding of these other bHLH TFs can vary substantially among E-boxes. Interestingly, changes in these TFs’ activity could alter how they compete with CLOCK:BMAL1, and thus affect CLOCK:BMAL1 DNA binding without altering its DNA binding affinity. Extending SMF to more CREs targeted by CLOCK:BMAL1 should help address this possibility.

SMF also provided valuable insights into the cooperation between CLOCK:BMAL1 molecules at CREs having multiple E-boxes. At *Nr1d1* 1^st^ intron, CLOCK:BMAL1 did not co-bind the two E-boxes of its preferred dual E-boxes motif, but rather engaged into co-binding events at E-boxes separated by 9 or more bp. Additionally, SMF revealed that *in vivo*, dual E-boxes are either unoccupied or bound simultaneously by two CLOCK:BMAL1 molecules. This contrasts with gel shift assays, where binding of CLOCK:BMAL1 to only one of the two E-boxes is observed^34^, and suggests that the priming binding event is a key regulatory step *in vivo*. Interestingly, a long-range interaction of 274 bp was only observed at ZT06 but not ZT18 and BMKO mice, and was associated with differences in E-boxes protection that imply structural changes in CLOCK:BMAL1 interface with DNA. Given the over 50-fold difference in *Nr1d1* transcription between ZT06 and ZT18^64^, and the low-level protection at E-boxes 4/5 at ZT18, this indicates that *Nr1d1* transcription is mostly initiated when CLOCK:BMAL1 is co-bound to E-boxes 2/3/4/5. To some extent, these data about cooperation between CLOCK:BMAL1 molecules also support recent findings showing that CLOCK:BMAL1-mediated transcription does not just rely on its binding to a single E-box, but rather on its cooperation with other TFs^27–29^. A recent study showed that TF DNA occupancy linearly correlates with gene expression^65^, suggesting that the cooperation among multiple CLOCK:BMAL1 proteins on this CRE may be important for *Nr1d1* expression in mouse liver.

CLOCK:BMAL1 was recently reported to bind the entry/exit site of nucleosomes and to interact with the H2A-H2B acidic patch^31^, a mechanism that would explain its pioneer-like activity^30^. Remarkably, we found 66 nucleosomes (out of the 30 CREs we analyzed in this study) harboring an E-box at their entry/exit sites, with 22 of them on 15 CREs showing decreased nucleosome signal in WT ZT06 compared to WT ZT18 or BMKO mice. CLOCK:BMAL1 binding at the entry/exit sites of nucleosomes is thus prevalent in the mouse genome, and it importantly promotes the removal and/or sliding of nucleosomes. This nucleosome remodeling is likely achieved by CLOCK:BMAL1-mediated recruitment of chromatin remodelers/modifiers^66^, and may also involve CLOCK histone acetyltransferase activity^67^. By generating a more open chromatin landscape, CLOCK:BMAL1 increases the capability of other TFs that preferentially bind exposed DNA to access their binding sites. We found that this nucleosome-mediated TF cooperation mechanism applies to at least two protection events that contain a binding motif for RFX TF family. Increased SMF resolution, which is currently limited to GpC dinucleotide, is likely to uncover additional TFs whose binding is increased by CLOCK:BMAL1-mediated chromatin opening.

In summary, our *in vivo* SMF study in mouse liver provides a mechanistic understanding of how CLOCK:BMAL1 DNA binding at CREs evicts/moves nucleosomes and increases DNA accessibility for other TFs that preferentially bind naked DNA to regulate gene expression across the 24-hour day. Our analysis pipeline also provides the framework to using SMF for mechanistic studies interrogating how TFs and other chromatin associated proteins remodel the chromatin environment at CREs to favor the overrepresentation of chromatin conformations that are permissive to transcription.

### Limitations of the study

While SMF provides precise, single-molecule maps of chromatin configurations and nucleosome positioning, it does not determine whether specific nucleosomes harbor distinct histone modifications and/or variants. Such information would be essential to establish the molecular basis of transitions between chromatin states and to link the structural configurations identified here to TF binding and, more broadly, transcriptional activity. Distinguishing whether particular configurations correspond to active, repressed, or poised states will be critical to fully understand how nucleosome remodeling influences transcriptional bursting. Integrating SMF with assays that profile histone marks or variants would help connect the spatial organization of nucleosomes to their biochemical identity and reveal whether specific modifications predispose a CRE to particular conformational transitions.

In addition, SMF cannot identify the TFs responsible for individual footprints, which must instead be inferred from motif sequence and available ChIP-Seq datasets. This limitation is especially relevant for motifs like E-boxes that are recognized by numerous members of the bHLH family, many of which can compete for the same binding sites. The development of approaches combining SMF with assays that profile TF binding will be necessary to dissect the hierarchy, interdependence, and temporal competition among these regulators at single-molecule resolution.

Finally, our study focused on a selected set of 30 CLOCK:BMAL1-bound CREs rather than the entire genome. Although these loci capture diverse regulatory configurations, a broader survey would be required to assess how representative these mechanisms are across tissues, environmental conditions, and genomic contexts. While technically challenging given our required sequencing depth for clustering analysis, extending this approach genome-wide will be important to evaluate the generality of the mechanisms we uncovered and to define how the interplay between circadian TFs like CLOCK:BMAL1, other TFs, and nucleosome remodeling contributes globally to the regulation of cycling transcriptomes.

## Acknowledgments

We thank members of the Menet lab for helpful discussions throughout the project; Alicia Michael, Paul Hardin, and Joseph Rodriguez for insightful suggestions and comments on the manuscript; and Ana Velasquez for her contribution at the early stage of this project. We thank Jared Bard for kindly providing HEK293T cells for the luciferase assay, and Diyun Sun for helping with the cell culture setup. We thank Audrey Jacq for helping collect brain samples and for discussions throughout the project. We also thank Michael Kladde for helpful discussions in the initial stage of this study. Portions of this research were conducted with the advanced computing resources provided by Texas A&M High Performance Research Computing. This work was supported by a NIH NIGMS grant (R01GM145737) and startup funds from Texas A&M University (JSM). JSM is also supported by a NIH NIDDK grant (R01DK128133), and YJ by a NIH NIGMS grant (R35GM138342).

## Author contributions

X.Y.N. and J.S.M. conceived and designed the research, analyzed the data, and wrote the manuscript. X.Y.N. performed the experiments and bioinformatic analyses. N.A. set up and performed CUT&RUN. A.H. N.A. and X.Y.N. set up and performed SMF in other mouse tissues. G.A. helped with the luciferase assays. J.S.M. supervised the study. All authors read and edited the manuscript.

## Conflict of interests

The authors have no competing interests to declare.

## Methods

### Mice

Experiments with mice were approved by the Texas A&M University Institutional Animal Care and Use Committee. Adult male wild type (WT; C57BL/6NCrl strain) and *Bmal1^-/-^* mice (BMKO; backcrossed to C57BL/6NCrl background for a minimum of 8 generations) were housed under 12 h light:12 h dark, and provided food and water *ad libitum*. Mice were euthanized by isoflurane anesthesia followed by decapitation in the middle of the day (ZT06; WT and BMKO) or night (ZT18; WT only), with three biological replicates per group. Liver, kidney, spleen, muscle, lung, and cortex were dissected quickly after euthanasia, briefly washed in ice-cold 1X PBS, snap-frozen in liquid nitrogen, and stored at -80°C until nuclei purification.

### Single molecule footprinting

The SMF protocol was adapted from Sönmezer et al., 2021^6^ for 6 mouse tissues (liver, kidney, spleen, muscle, lung, and cortex) included in this paper as described in Michael et al., 2023^31^. We adapted two protocols for nuclei purification. Nuclei from nine liver samples (WT ZT06, WT ZT18 and BMKO ZT06; 3 replicates of each) were purified as described in Menet et al., 2012^64^. For each reaction, 250,000 liver nuclei were washed (50 mM Tris pH 8.5, 50 mM NaCl, 10 mM DTT) and then resuspended in 1ml M.CviPi reaction buffer (50 mM Tris pH 8.5, 50 mM NaCl, 300 mM sucrose, 10 mM DTT). Because this isolation procedure did not work well with other tissues, we opted to the OMNI-ATAC procedure described in Corces et al. 2017^68^ to isolate nuclei from the cortex, kidney, spleen, muscle, and lung (WT ZT06, WT ZT18, and BMKO ZT06; 2 replicates of each). At the last step, nuclei collected from the gradient were mixed with 1x SMF wash buffer (1 mM HEPES pH 7.6, 0.5 mM KCl, 0.15 mM MgCl2, 0.1 mM EDTA; 3 to 4 times the volume of the collected nuclei). Nuclei were then pelleted at 5000g centrifuge for 15min at 4°C.

For each sample, M.CviPi methylation reactions were performed using 250,000 nuclei. Specifically, nuclei were resuspended in 1ml M.CviPi reaction buffer (50 mM Tris pH 8.5, 50 mM NaCl, 300 mM sucrose, 10 mM DTT), to which 18.75 µl of 32 mM SAM and 200U of M.CviPi were then added. After 7.5 min incubation in a 37°C water bath, the reaction was supplemented with 100U M.CviPI and 128 µmol of SAM for another round of 7.5 min incubation at 37°C. Then, 350 µl of SDS containing buffer (20 mM Tris, 600 mM NaCl, 1% SDS 10 mM EDTA) and 20 µL of Proteinase K (20 mg/ml) were added to stop the reaction, and samples were incubated overnight at 55°C. Genomic DNA (gDNA) was purified by phenol-chloroform extraction, and 2 µg of gDNA was used for bisulfite conversion using the Epitect bisulfite conversion kit (QIAGEN 59124), or 200 ng gDNA was used for enzymatic conversion (NEB E7125). 41 CREs were selected for this study, including 30 CLOCK:BMAL1 targeted CREs. The first 28 CREs were selected based on CLOCK and BMAL1 ChIP-seq peaks. The remaining 13 CREs included two closed chromatin regions (no DNase-seq peak) and 10 CREs chosen because they are located in two genes (*Por* and *Inmt*) studied with the original 28 CREs. A minimum of 10-30 ng of converted DNA was used to amplify each of the 41 CREs as in Sönmezer et al., 2021^6^. Each PCR product was individually purified with 1.5x SPRI beads and PCR products from the same liver sample were mixed. If enough converted DNA was available, 20-30 ng was used for PCR amplification with only 20 cycles, and all PCR products for one sample were mixed and purified using 1.5x SPRI beads. Purified PCR products were pulled for each sample and used to generate sequencing libraries with NEBNext Ultra II Kit (amplification for 12 cycles if starting from 200ng, or 15 cycles if starting from less than 50ng). Libraries from the nine liver samples were pooled together, and libraries from tissues other than the liver were pooled together. Libraries were sequenced using either a MiSeq v2 Nano Reagent kit or a MiSeq v2 Reagent kit. (paired-end 250 bp).

### Sequence alignment

The python package and parameters used for alignment were described in Michael et al 2023^31^, and provided as supplementary file 1. Briefly, *pairwiseAligner* function from *Bio.Align* python package were used to align reads from fastq files, using parameters of matched, mismatched, and gapped score set as 1.0, -0.2 and -0.5 respectively. An alignment score was calculated based on the sum of alignment score divided by the length of query sequence, with the maximum score of 1.0. First, a pre-selection was performed to select reads that have an alignment score of 0.8 when aligning both forward and reverse primer sequences to the beginning ∼25nt sequences in the paired-end fastq files. The full sequence of pre-selected pair-end reads were then aligned to bisulfite converted target CRE sequences (HCH replaced by HTH, GC replaced by GY, CG replaced by YG, with Y= pyrimidine and H = not-G). Based on the alignment results at each nucleotide, the full-length query sequence was then reconstituted for reads with both paired-end sequence alignment scores higher than 0.7, and nucleotides having higher quality scores were used for overlapping region reconstitution between the paired-end read. Reads having a cytosine of a GCH position not sequenced as GC or GT were removed, along with PCR duplicates.

### UMI barcode

Amplification of nine CREs was performed using primers having no UMI barcodes, including the first 7 CREs in Figure S3A, *Nr1d2* intron, and *Srebf1* intron. The remaining 32 CREs were amplified using primers having UMI barcodes added in both forward and reverse primers. To avoid biased incorporation of UMIs mapping perfectly to the amplicon, UMI barcodes contained 4 mismatched nucleotides directly upstream of the priming sequence and 4 random nucleotides (N) in 5’. The four mismatched nucleotides were designed to avoid annealing with the template nucleotide, *e.g*., if the template was an adenine (A), then a letter B (= any nucleotide other than an A, i.e., C, G, or T) was incorporated in the UMI barcode. If the template had a GC or CG, then a letter R (= A or G) was incorporated in the UMI. Because of the 250bp sequencing length limitation for paired-end reads and our goal of sequencing GpCs in the middle of the amplicon, *Po*r CRE8 has 6 N plus 4 mismatches in the forward primer, *Por* CRE7 and CRE10 has 6 N plus 5 mismatches in the forward primer, and none of the reverse primers for these 3 CREs had an UMI. Sequences of primers are provided in Table S1, and details about which CREs were sequenced in each fastq file are provided in Table S2.

Sequence alignment of CREs amplified with UMI-containing primers was performed as above with slight modifications. First, reads were called unique if they had a unique barcode in both forward and reverse primers. Then, if several reads had the same UMI combinations, then the read with the highest total quality score was kept. This score was calculated by adding the quality score (QC) at each position if QC ≥ 15, which is similar to samtools (as in *DuplicateScoringStrategy* class in samtools open-source code)^69^. A random read was chosen if multiple reads had the same maximum total quality score.

### BMD Clustering

Only reads that successfully passed the sequence alignment steps were used for downstream analysis. First, the sample with the lowest number of reads among the nine samples was identified for each CRE, and its number of reads was used to downsample by random selection the number of reads in the other eight samples such that all nine samples contained an equal number of reads for each CRE in subsequent steps. Conversion information was extracted at each GCH position (except for GCH positions in the PCR primers) for each read, using 0 or 1 to represent unprotected (sequenced as GC) or protected (sequenced as GT), respectively. In the *Nr1d1* 1^st^ intron, one GpC in the middle of the amplicon was not covered by the sequencing of either paired end read and was thus removed from analysis. Binary Matrix Decomposition (BMD) clustering was applied on randomly selected reads from all nine samples for each CRE^36^. The *generalBMD* function from *bmdcluster* python package was used, with the parameter b (number of points to initialize bootstrap seed points) set as 1% of the total number of reads, and parameter seed (the pseudorandom number generator) set as 1. The final cluster number was defined by increasing stepwise from 2 clusters until the number of reads in a cluster was less than 1% of all reads, and set such that no cluster contained less than 1% of the total number of reads. Once the number of clusters was set, reads were parsed based on their relative cluster and experimental group (WT ZT06, WT ZT18 and BMKO ZT06).

### Computational definition of nucleosome protection

A custom python function named *defineRegionCombinedSamples* (provided as supplementary file 2) was generated to automatically define the protection profile as either nucleosome or exposed DNA (bound or not by a TF) for each cluster. A figure illustrating the criteria used in this study and described below is provided as Fig. S12. First, if the percentage of protection at three or more consecutive GpCs was larger than 60% (each of them), or if the percentage of protection at four or more consecutive GpCs was larger than 50% (each of them), then the position of the first and last GpCs of this putative nucleosome was determined. If the distance between the first and last GpCs was over 75 bp, or if the distance between the first and last GpCs was between 60 bp and 75 bp but the distance between the GpC before the first and the GpC after the last was over 120 bp, the footprint was defined as nucleosome protection.

Because nucleosome protection signal may be truncated at the 5’ or 3’ end of each CRE/amplicon, we applied two other criteria for this specific situation. For the first criteria, a footprint was defined as nucleosome if the percentage of protection was larger than 50% at two or more consecutive GpCs starting from the first or last GpC of the PCR amplicon (first 5’ GpC or last 3’ GpC), and if the distance between the first and last GpC of this footprint was over 20bp (many TFs have a footprint < 20 bp^14^). The second criteria looked for patterns having continuous protection on the same DNA molecule starting from the first or last GpC of the PCR amplicon (first 5’ GpC or last 3’ GpC). In the 5’ to 3’ direction, if the protection of the first GpC was over 50% and the protection of the next GpC was over 40%, then, we calculated whether the two protection events occurred on the same DNA molecule. This step was called “*continuity value*” (*e.g*., in the script provided in supplementary file 2), and is defined as the number of reads being protected at two consecutive GpCs on the same DNA molecule divided by the number of protected reads at the first of the two GpCs. If the resulting “*continuity value*” was over 70%, the process was repeated for the next GpC(s) until the value was less than 70% or the GpC percentage of protection was lower than 40%. Finally, we required for this second criteria that the distance between the first and last protected GpCs to be over 15bp to be defined as nucleosome. The same procedure was carried out starting from the 3’ end of the CRE amplicon, with criteria applied in the 3’ to 5’ direction.

Another additional analysis was then carried out to define nucleosomes, and relied on the continuity of protection events on the same DNA molecules as described above with what we called the “*continuity value*” . Starting from the 5’ end of the CRE/amplicon, the algorithm identified GpC(s) having a percentage of protection over 40% and not defined as nucleosome. If the percentage of protection of the next GpC (in 3’) was over 30%, then the algorithm calculated the continuity value between the two GpCs, and continued computing this value with the next GpCs in 3’ until the continuity value was lower than 70% or the percentage of protection of the next GpC was lower than 30%. If 5 consecutive GpCs matched those criteria and the distance between the first GpC and the last GpC was over 75 bp, this footprint was defined as a nucleosome. The algorithm was also run in reverse orientation, using criteria starting from the 3’ end of the amplicon and running toward the 5’ end.

After defining nucleosomes using all criteria described above, the algorithm started checking the undefined GpCs that were next to the nucleosomal GpCs. The aim of this step was to avoid mixing signals that were likely nucleosomes and being called as TF signals. At the GpC adjacent to the first GpC or last GpC of a nucleosome, if the percentage of protection was over 1%, then, the algorithm calculated the continuity value. If this value higher than 70%, the footprint at this GpC was defined as nucleosome. The algorithm then continuously looked at the next adjacent GpCs until the continuity value was lower than 70% or the percentage of protection at the GpC was less than 1%.

These criteria were applied all across our analyses, except in cluster 14 at *Nr1d1* 1^st^ intron for the calculation of TF cooperativity (Fig. 4B) and of levels of exposed DNA at CRE (Fig. 5D, S11B), where the definition at four GpCs between 424 to 441 was curated to exposed DNA based on visual inspection.

### Analysis of protection profiles by simple matching coefficient

A total of 1,692 reads for each sample were randomly selected for the CRE at *Inmt* TSS upstream enhancer (number of reads corresponding to half of the reads of the sample with the lowest number of reads). Simple matching coefficient was calculated between each pair of reads using the formula below:

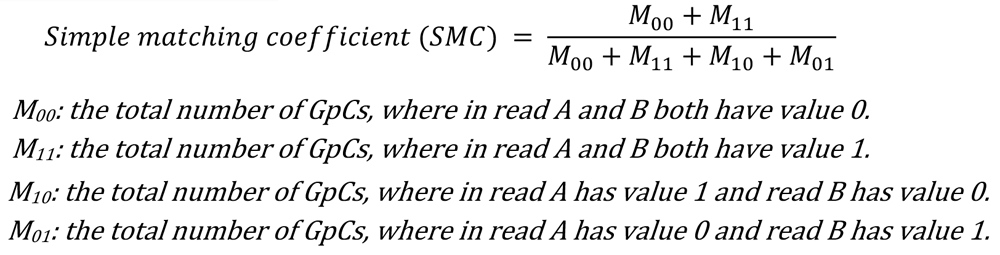

A hierarchical clustering was then performed on the matrix of SMC between each pair of reads (total of 15,228 reads and over 115 million pairwise comparisons), using *hierarchy* function (parameters of method and metric was set as ‘average’ and ‘euclidean’, respectively) from *scipy.cluster* python package. The heatmap illustrating the output of the hierarchical clustering of SMC was plotted with an entry of the source of each read. The percentage of reads from each sample was calculated for the first ten branches of the dendrogram.

### Analysis of protection profiles by PCA

A binary matrix with reads represented as vectors of binary values at GpCs was computed using reads at *Inmt* TSS upstream enhancer originating from all nine samples (GpCs output as a column, and each read corresponding to a row). PCA was performed using this binary matrix and the *PCA* function from *sklearn*.*decomposition* python package and the number of components set as the number of GCH within CRE. The output of the PCA analysis was then parsed for each sample, with PC1 plotted over PC2.

### BMD Clustering of individual samples

BMD clustering was performed for each individual sample with reads mapping *Inmt* TSS upstream enhancer. Nucleosome footprints were defined for every cluster of each sample using the custom-made function *defineRegionCombinedSamples* (see above). Outputs of this cluster definition (as either nucleosome or exposed DNA) were merged for all clusters of all nine samples and used to carry out hierarchical clustering. Five major hierarchical clusters were used to group the BMD clustering-based definition of chromatin states.

### Conversion efficiency

Conversion efficiency was performed on reads where cytosines at GCH position were sequenced as either GC or GT (but not GN, GA, or GG) and after PCR duplicates removal. Computation of bisulfite or enzymatic conversion efficiency was performed by calculating the percentage of cytosines at HCH positions sequenced as A, T, G, C or N at all for each CRE and for each sample, i.e., by dividing the number of A, T, G, C or N to the total number of HCH positions in all reads of each CRE per sample, as shown in Fig. S2.

### Conservation of chromatin states

A biological replicate was considered represented in a cluster/chromatin state if at least 1% of the reads of the cluster were assigned to that sample. This criterion was used to calculate the number of biological samples found in each cluster at each CRE, as reported in Fig. 1D and S3A, B.

### DNase-seq and ChIP-seq dataset analysis

Mouse liver DNase-seq (GSE37074^70^), and liver ChIP-seq datasets for BMAL1 (GSE39860^35^ and GSE110602^28^), CLOCK (GSE39860^35^ and GSE117488^71^), USF1 (GSE44609^60^), TFE3 (GSE160292^72^), MYC (GSE76042^73^), and bHLH40 (GSE207199^74^) were downloaded as fastq files and mapped to mouse genome version mm39 using bowtie2 (if read length ≥ 51 bp) or bowtie (of read length ≤ 50 bp). Duplicated reads were removed and bw files were generated by normalizing to the number of uniquely mapped reads to 10,000,000 reads for visualization.

### Motif analysis

TF motif analysis was performed using DNA sequences of ∼20 bp centered on the GpC(s) exhibiting TF protection using online Tomtom motif comparison tool (version 5.5.5).

### Calculation of exposed DNA level at CREs

BMD clustering was performed for each CRE with reads from all nine samples downsized to the sample with the lowest number of reads. Then, protection profile was defined at each GpC in each cluster of each CRE using the custom-made function *defineRegionCombinedSamples* (see computational definition of nucleosome protection section above), with each GpC ending up defined as either nucleosome (N) or exposed (E). This step located the two GpCs at the edge of nucleosomes for every cluster (GpCs defined as N). To improve the definition of nucleosome coverage and avoid an overestimation of exposed DNA at CREs, we then retrieved the position of the GpC before the first and after the last GpCs of a nucleosome (these GpCs being defined as E), and extended the nucleosome position to the middle position between the two linked N and E GpCs. Finally, nucleosome occupancy in each cluster was calculated by dividing the length of DNA protected by nucleosome in a cluster to the total length of the CRE. The nucleosome occupancy of each cluster was multiplied to the percentage of reads of this cluster, which were then summed between all clusters of a CRE to represent the total nucleosome exposure of a CRE. The exposed DNA level of a CRE was finally calculated by subtracting the total nucleosome exposure of a CRE to 1. Formulas are provided below:

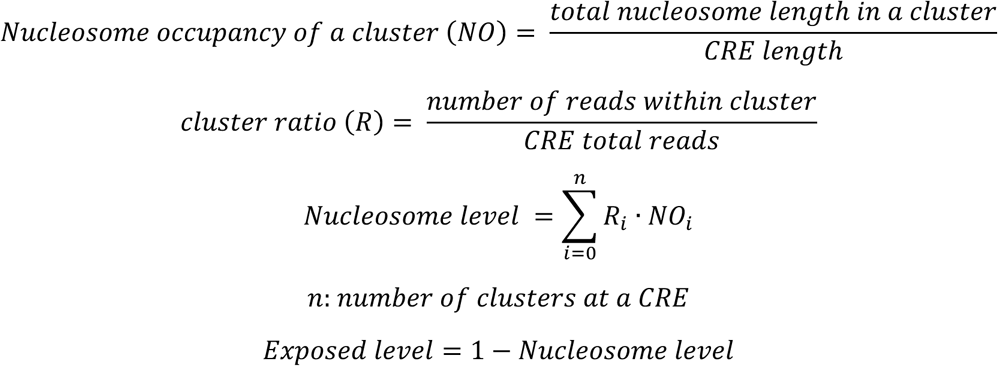

### Cooperativity between TFs

Calculation of the normalized extend of co-binding (N. EOC) between TFs at two GpCs was adapted from Rao et al., 2021^5^ with slight modification, and carried out in clusters where both GpCs were not defined as nucleosomes using the formula:

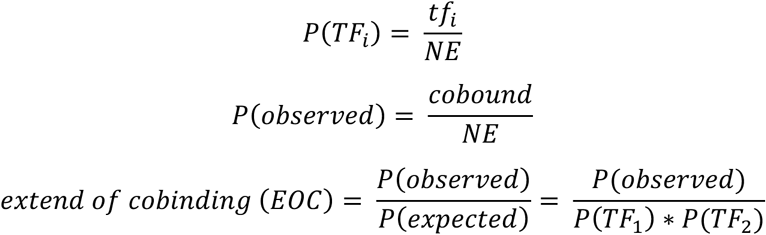

Where:

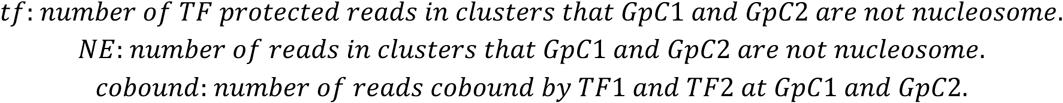

Then the normalized extend of co-binding was calculated using the formula:

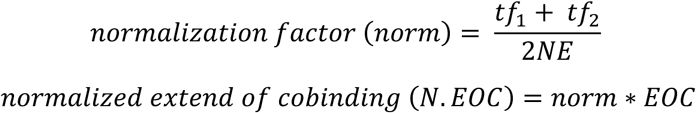

P-values were calculated by Fisher’s exact test (null hypothesis: protection at two GpCs are independent), followed by Benjamini/Hochberg correction to adjust the p value to multiple testing, using *fdr_correction* function from *mne.stats* python package, with the false discovery rate set as 0.05.

### MNase-seq analysis

24 WT and 23 BMKO single-end mouse liver MNase-seq datasets from Menet et al., 2014^30^ (GSE47145) were uploaded as fastq files and mapped to the mouse genome version mm39 using bowtie2. Reads located +/- 1 kb from each CRE were selected using samtools, and extended to 147 bp to map full length nucleosome. Sequences from all datasets were merged and sorted using bamtools and samtools, respectively. Finally, relative nucleosome signal was normalized to the total number of mapped reads from all 47 samples to generate bigwig (bw) files. A similar analysis was performed using sequence definition/position adjusted to cover 75 bp centered on the nucleosome dyad to better visualize nucleosome position.

Analysis of paired-end mouse liver low and high MNase-seq datasets (nuclei treated with low and high MNase concentration) from Iwafuchi-Doi et al., 2016^49^ (GSE57559) was performed as above with only slight differences. Selection of sequencing reads included an additional step, where only concordant paired reads with a size between 100 bp and 200 bp and a bam-file flag number of 99 or 163 were selected. The analysis was performed on four merged high MNase-seq datasets (2 replicates C3H WT mice and 2 replicates from C57BL6 WT mice), and on four merge low MNase-seq datasets (2 replicates C3H WT mice and 2 replicates from C57BL6 WT mice). The bw files were generated by normalizing MNase-Seq signal to the total number of mapped reads having a bam-file flag of 99 or 163 from the four high MNase-seq or low MNase-seq datasets.

### Comparison between SMF and MNase-seq nucleosome signals

SMF nucleosome signal for each sample at each GpC was calculated by adding the percentage of reads from clusters that were defined as nucleosome at that GpC. The averaged value from all nine samples was then calculated for each GpC of each CRE. The MNase-seq nucleosome signal at each GpC was extracted from the bw file generated from Menet et al., 2014 datasets as described above^30^, using reads extended to 147 bp for all 47 datasets. The MNase-seq nucleosome signal and averaged SMF nucleosome signal were calculated at a total of 1137 GpCs from 41 CREs, and used to carry out the Pearson correlation.

### Analysis of nucleosomes having an E-box at the entry/exit site

A custom python script was generated and applied to automatically select nucleosomes that had E-box(es) at the entry/exit site, followed by a limited curation based on visual inspection. Nucleosomes were initially selected if E-boxes were located 20 bp within a nucleosome as defined by our custom script *defineRegionCombinedSamples* (the edge of the nucleosome corresponding to a GpC location, see above), or 20 bp outside the nucleosome but without a GpC located between the E-box and the edge of the nucleosome. Then, some criteria were applied to further refine the selection. If the nucleosome location started at the first GpC in 5’ or 3’ of the CRE (i.e., first GpC amplified when carrying out the PCR of bisulfite converted DNA), and the E-box was also located on the 5’ or 3’ side, this nucleosome should protect over a range of 110 bp. Nucleosomes with protection over 200 bp were excluded (e.g., condensed chromatin) unless there was a dip in the percentage of protection between two observed nucleosomes. Nucleosomes representing less than 5% of all reads were excluded from downstream analysis.

### Transactivation assays

HEK293T cells were cultured under 5% CO_2_ at 37°C in Dulbecco’s Modified Eagles Medium (DMEM; Cytiva HyClone), containing 100U/ml streptomycin/penicillin antibiotics (ThermoFisher) and 10% fetal bovine serum (FBS; ThermoFisher), as complete DMEM medium. For transactivation assays, 1.4 x 10^5^ cells were seeded in 24-well plates and cultured for 24h prior to transfection. Two hours before transfection, the culture medium was replaced with Opti-MEM (ThermoFisher). Plasmids (500ng/well) were transfected to cells at ∼70% confluency using Lipofectamine 3000 (ThermoFisher) following the manufacturer’s protocol. 12h post-transfection, the medium was replaced with complete DMEM (with reduced FBS to 5%). Cells were harvested 24h later to measure luciferase activity using the Dual Luciferase Reporter Assay kit (Promega) on a luminometer (SpectraMax iD3; Molecule Devices). For each condition, DNA-lipid complexes were prepared as a 2-well amount (1 ug of DNA) and evenly distributed into two wells, serving as technical replicates. Three independent biological replicates were performed for each assay.

For *Nr1d1* reporter transfection assays, transfections were performed with and without CLOCK:BMAL1. For each transfection, we used 1ug of DNA consisting of 50ng of pGL3-promoter constructs (empty vector, or Nr1d1 1^st^ intron WT or E-box mutant; Firefly luciferase) and 50ng pcDNA3.1-Renilla (Renilla luciferase), and for CLOCK:BMAL1 groups, 300ng pcDNA3.1-mBMAL1 plus 300ng pcDNA3.1-mCLOCK. The remaining DNA was supplemented with pcDNA-LacZ to a total of 1ug for all conditions.

For DEC1 competition assays, the DNA-lipid complex was prepared similarly. The 1ug DNA consisted of 50ng of pGL3-promoter constructs (*Nr3c2* intron or *Por* distal enhancer; firefly luciferase) and 50ng pcDNA3.1-Renilla, 100ng pcDNA3.1-mBMAL1, 100ng pcDNA3.1-mCLOCK, and various amounts of pcDNA3.1-mDEC1 (0, 3.2, 8, 20, or 50ng). pcDNA-LacZ was added to bring the total DNA to 1ug. The condition containing the empty pGL3-promoter vector, without adding any TFs, served as the negative control. Three biological replicates were performed for each assay. The activity of each *Nr1d1* 1^st^ intron reporter was calculated by subtracting the firefly/renilla ratio of no CLOCK:BAML1 from that measured in their presence. Then the values for each replicate in each condition were normalized to the mean activity of the WT *Nr1d1* 1st intron reporter.

### Plasmid constructs

The pGL3-promoter vector was purchased from Addgene (plasmid cat#: 212939). *Nr1d1* 1st intron reporter constructs, *Nr3c2* intron, and *Por* distal enhancer reporter constructs, and DEC1 coding sequence were synthesized and cloned by GeneScript. WT and mutant enhancer sequences along with DEC1 sequence are listed in table S3.

### Single nucleus RNA-seq

#### Sample preparation

Six liver samples (WT ZT6) were crushed into powder, and the same volume of powder from three mice was mixed for each nuclei isolation as described in Menet et al., 2012^64^ and above in the SMF section. Pelleted nuclei were washed once (1 mM HEPES pH 7.6, 0.5 mM KCl, 0.15 mM MgCl2, 0.1 mM EDTA, 0.04% BSA). The snRNA-seq was performed using 10x Genomics GEM-X Flex Sample Preparation v2 Kit. 6 million nuclei were pelleted, and the wash buffer was completely removed. Following the manufacturer’s protocol in the Fixation of Cells & Nuclei section, nuclei were fixed at 4°C for 17h in fixation buffer B. On the same day that the fixation reaction was quenched, 500,000 fixed nuclei were loaded for probe hybridization using Chromium Mouse Transcriptome Probe Set v1.1.1. The library was prepared following the manufacturer’s protocol, and was sequenced using NextSeq 2000 P3 XLEAP kit for 100 cycles, 1 billion reads.

#### Data analysis

The filtered count matrix was generated following Cell Ranger (v9.0.0) pipeline for downstream pre-processing and cell type identification analysis using Scanpy^75^. For each replicate, we filtered cells that had less than 200 genes and removed genes that were expressed in less than ten cells, from which the top 2000 highly variable genes were used for doublet prediction using the Solo framework^76^ implemented in *scvi-tools*^77^. Cells predicted as doublets with a double and singlet score difference greater than 0.2 were removed from downstream analysis, resulting in no doublets in our data. A second round of filtration was performed after doublet removal. Cells were retained if they expressed fewer than the 98th percentile of genes per cell, <20% mitochondrial counts, and <2% ribosomal counts. Finally, cell type identities were assigned by transferring annotations from the reference liver cell atlas (LCA)^78^ using the Scanpy ingest function.

#### CUT&RUN

CUT&RUN was performed for BMAL1 and RFX1 in liver samples from WT ZT06, WT ZT18, and BMKO ZT06 (3 replicates of each), using a procedure adapted from established protocols^79,80^ and from Epicypher webiste. Nuclei were purified as previously described^64^. Briefly, ∼300 mg of liver powder was homogenized in 4 ml of 1x PBS containing 0.5% formaldehyde for 4 min at room temperature, then quenched by mixing with 25 ml of ice-cold quenching solution (2.2 M sucrose, 150 mM glycine, 10 mM Hepes pH 7.6, 15 mM KCl, 2 mM EDTA, 1 mM PMSF, 0.15 mM spermine, 0.5 mM spermidine, and 0.5 mM DTT). After 10min incubation on ice, the homogenate was carefully layered over a 10 ml ice-cold sucrose cushion (2.05 M sucrose, 125 mM glycine, 10 mM Hepes pH 7.6, 10% glycerol, 15 mM KCl, 2 mM EDTA, 1 mM PMSF, 0.15 mM spermine, 0.5 mM spermidine, and 0.5 mM DTT). Nuclei were pelleted by ultracentrifugation (45min, 100,000 g, at 4°C ; Beckman SW32Ti rotor), and washed once with wash buffer (20mM HEPES, 150mM NaCl, 0.5mM spermidine, 1× protease inhibitor EDTA-free). 400,000 nuclei were bound to activated CUTANA Concanavalin A conjugated paramagnetic beads (EpiCypher) and incubated overnight at 4°C with 1 μg antibody (anti-BMAL1: Abcam, AB3350; anti-RFX1: Bethyl, A303-043A) on a nutator, following the manufacturer’s protocol. The next day, nuclei were carefully washed twice with 200 µl Dig-wash (Digitonin-wash) buffer (5% digitonin, 20mM HEPES, 150mM NaCl, 0.5mM spermidine, 1× protease inhibitor EDTA-free) on a magnetic stand. 2.5 µl pAG-MNase (EpiCypher) was added to the nuclei in 50µl Dig-wash buffer, and incubated at room temperature for 15min. The reaction was initiated by adding 1 µl 100mM CaCl_2_ on ice, and incubated for 2h on a nutator at 4°C, which was then quenched by adding 33µl stop buffer (340mM NaCl, 20mM EDTA, 4mM EGTA, 50µg/ml RNase-A, 50ug/ml Glycogen, and 0.5 pg/ml *E.coli* Spike-in DNA) and incubated at 37°C for 20min. The supernatant that contained fragmented DNA was extracted by phenol: chloroform purification. Libraries were prepared using NEBNext ultra II DNA library Prep kit (NEB 7103). After the adaptor ligation step, the DNA was cleaned twice with 1.75x Ampure beads (Beckman). Libraries were amplified for 15 cycles. Fragments larger than 350bp were removed by taking the supernatant after incubation with 0.8x Ampure beads. The supernatant was then incubated with a final effective concentration of 1.2x Ampure beads to keep small insert fragments (∼50-200bp). Libraries were sequenced as paired-end using NextSeq 2000 P2 XLEAP kit for 2x50 cycles.

#### CUT&RUN and ChIP-Seq analysis

CUT&RUN reads were aligned to the mouse genome (version mm39) using bowtie2 with the parameters --end-to-end and --maxins 1500. PCR duplicates were removed and uniquely mapped reads were only considered for analysis. Visualization files were generated using bedtools and normalized to 10,000,000 reads. Files were processed individually and reads from biological replicates were merged as bam files using samtools. Signal at CREs of interest was normalized to 10,000,000 reads and retrieved using bedtools.

Public mouse liver ChIP-Seq datasets were downloaded as fastq files and processed following the same pipeline as for CUT&RUN. Reads were aligned to the mouse genome using either bowtie (read length <= 50; parameters -q -a -m 1) or bowtie2 (read length > 50; parameters --end-to-end and --maxins 1500). Datasets consisted of DEC1: GSE207199^74^, USF1: GSE44609^60^, MYC: GSE76042^73^, ChREBP: GSE207199^74^, TFE3: GSE160292^72^, and BMAL1: GSE39860^35^ and GSE110602^28^.

## Data availability

All sequencing data have been depositing in Gene Expression Omnibus (GEO), which are currently protected but accessible using the token provided below. They include: liver SMF datasets: GSE255510 (token: qxarkqksthgpnsj), SMF datasets from 6 tissues: GSE307333 (token: snqdqeekpjqthkx), snRNA-seq: GSE307334 (token:ydkrqqugzlatdap), and CUT&RUN datasets: GSE307335 (token: kfehewimvlothmt).

## Code availability

The script used to align paired-end reads is provided as supplementary file 1, and the code used to computationally define nucleosome protection (*defineRegionCombinedSamples*) is provided as supplementary file 2.

## Figure representation

Box-plot elements (Fig. 1D, 2D, 5D, S3D, S9C) were represented as follow: center line, median; box limits, upper and lower quartiles; whiskers, 1.5x interquartile range; points, all data values.

---

**Figure S1.**
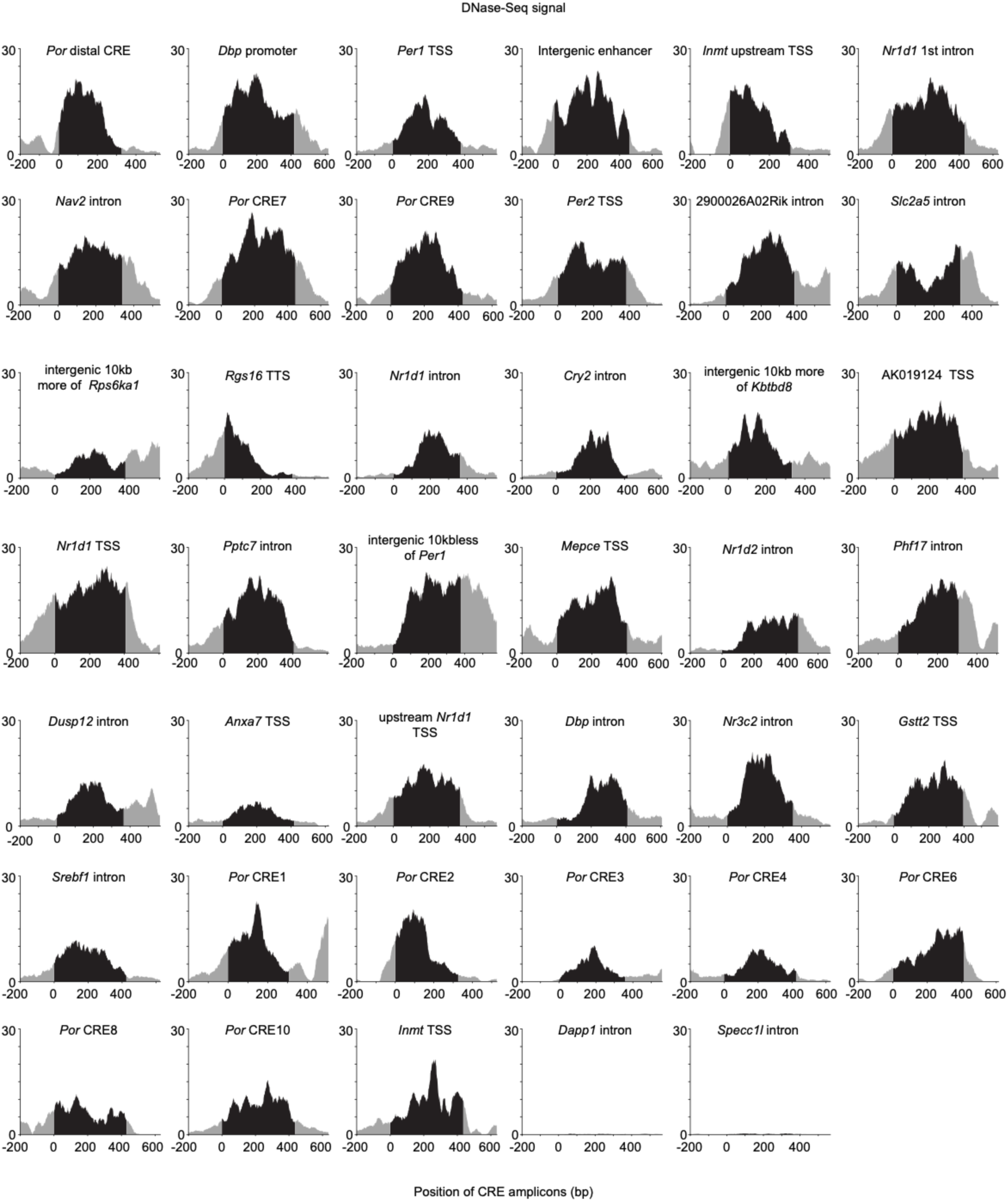
DNase-seq signal and SMF coverage at the 41 CREs investigated in this study. Mouse liver DNase-seq (from the Encode project^70^) signal at the 41 CREs selected for this study. The black regions represent the first to last GpC that were analyzed by SMF for each CRE. The grey regions are 200bp extended from upstream and downstream of the analyzed region.

**Figure S2.**
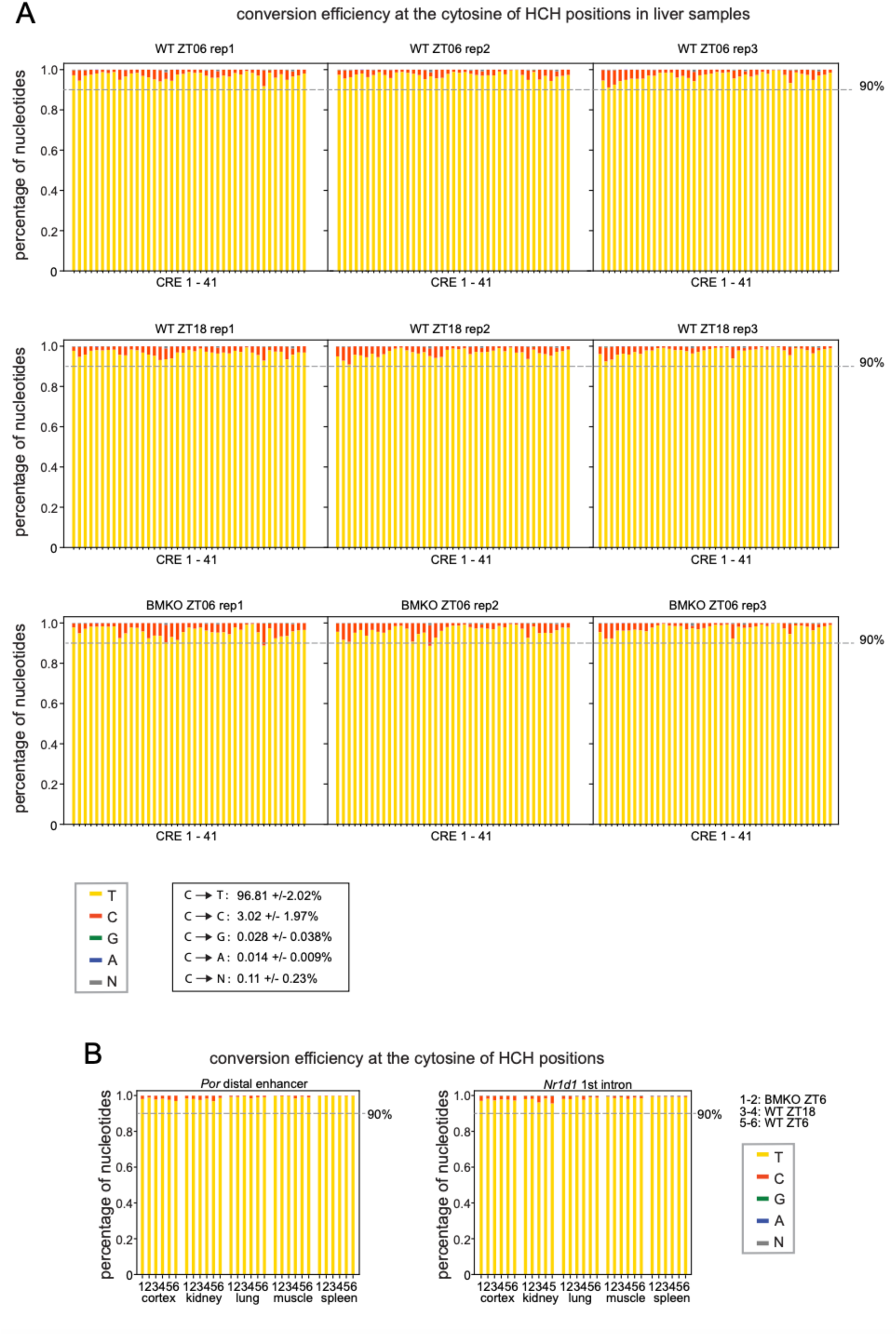
Conversion efficiency of cytosines at HCH positions. **(A, B)** Stacked bar plot representing the frequency of sequencing a cytosine as an adenine (blue), thymine (yellow), guanine (green), cytosine (red), or undefined nucleotide (N, grey) after bisulfite conversion or enzymatic conversion within each analyzed CRE for each animal. **(A)** 41 CREs amplified from liver samples collected from WT ZT06, WT ZT18, and BMKO ZT06 mice (three biological replicates per condition). Analysis was performed at HCH position (H = A,C,T), i.e., cytosines located in a CpG or GpC dinucleotide were excluded from this analysis. **(B)** *Por* distal CRE and a CRE located in *Nr1d1* 1^st^ intron were amplified from enzymatically converted DNA from cortex, kidney, kidney, lung, muscle and spleen that were collected from 2 replicates of BMKO ZT06, WT ZT06 and WT ZT18 mice.

**Figure S3.**
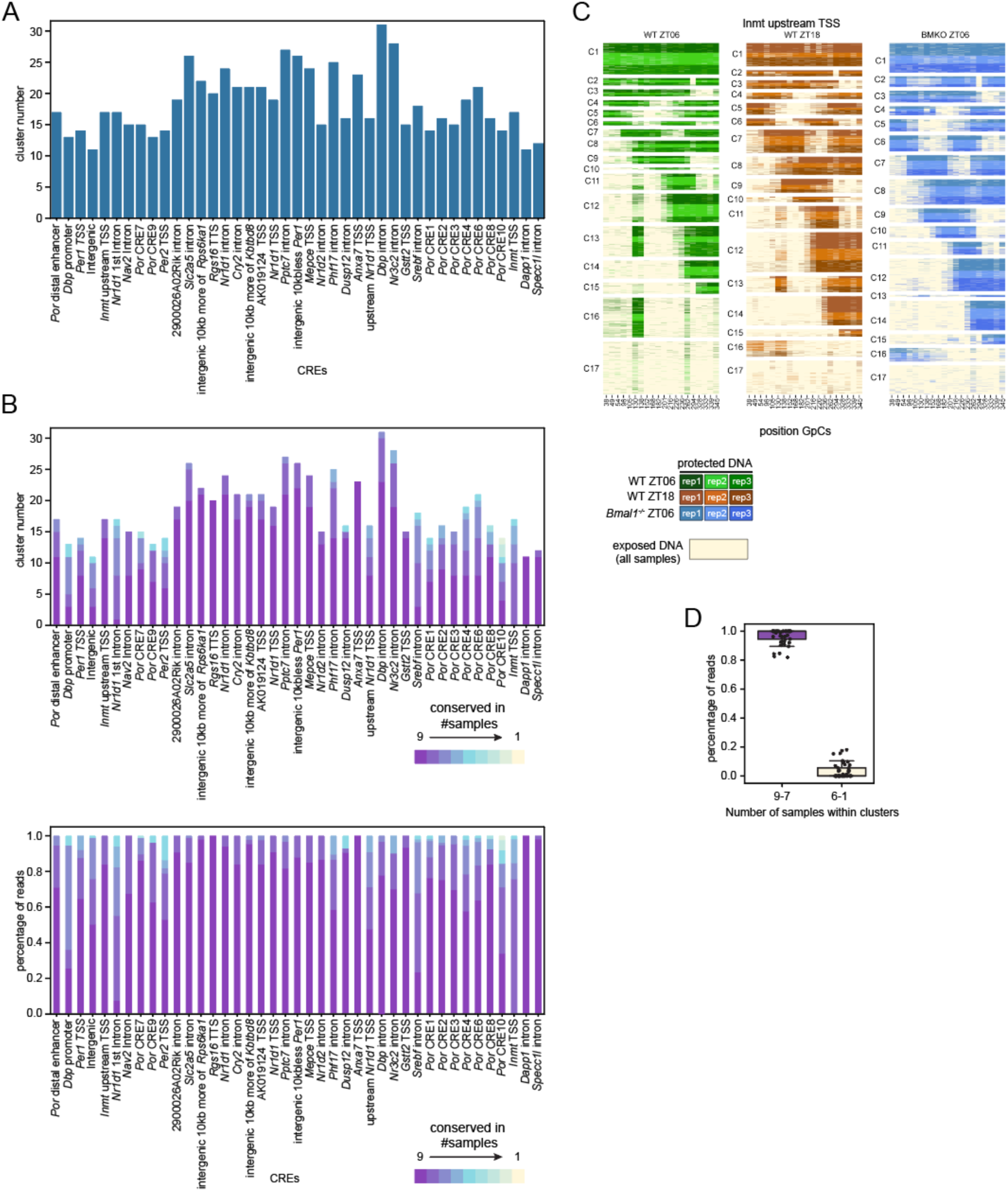
Summary of the BMD algorithm clustering analysis at all 41 CREs. **(A)** Number of clusters defined by BMD clustering analysis for the 41 CREs included in this study. **(B)** Top: number of clusters conserved between samples for each of the 41 CREs. Samples (i.e., biological replicates) were considered as not represented in a cluster if the percentage of reads within a cluster for that sample was less than 1%. Bottom: Percentage of reads in clusters found in 1-9 sample(s) for each CRE. Clusters found in 6 or less samples were generally small clusters representing less than 10% of all reads. **(C)** SMF profile at *Inmt* TSS upstream enhancer (chr6:55152994-55153301) in mouse liver parsed by group (WT ZT06, WT ZT18 and BMKO ZT06). Each heatmap contains reads from n=3 biological replicates that are shaded in different colors of green, brown, and blue. Colors are the same as in Figure 1C. **(D)** Percentage of reads in clusters conserved between 7, 8, and 9 samples (i.e., clusters contain reads from all groups) and between 1-6 samples. For each boxplot, a dot represents a CRE.

**Figure S4:**
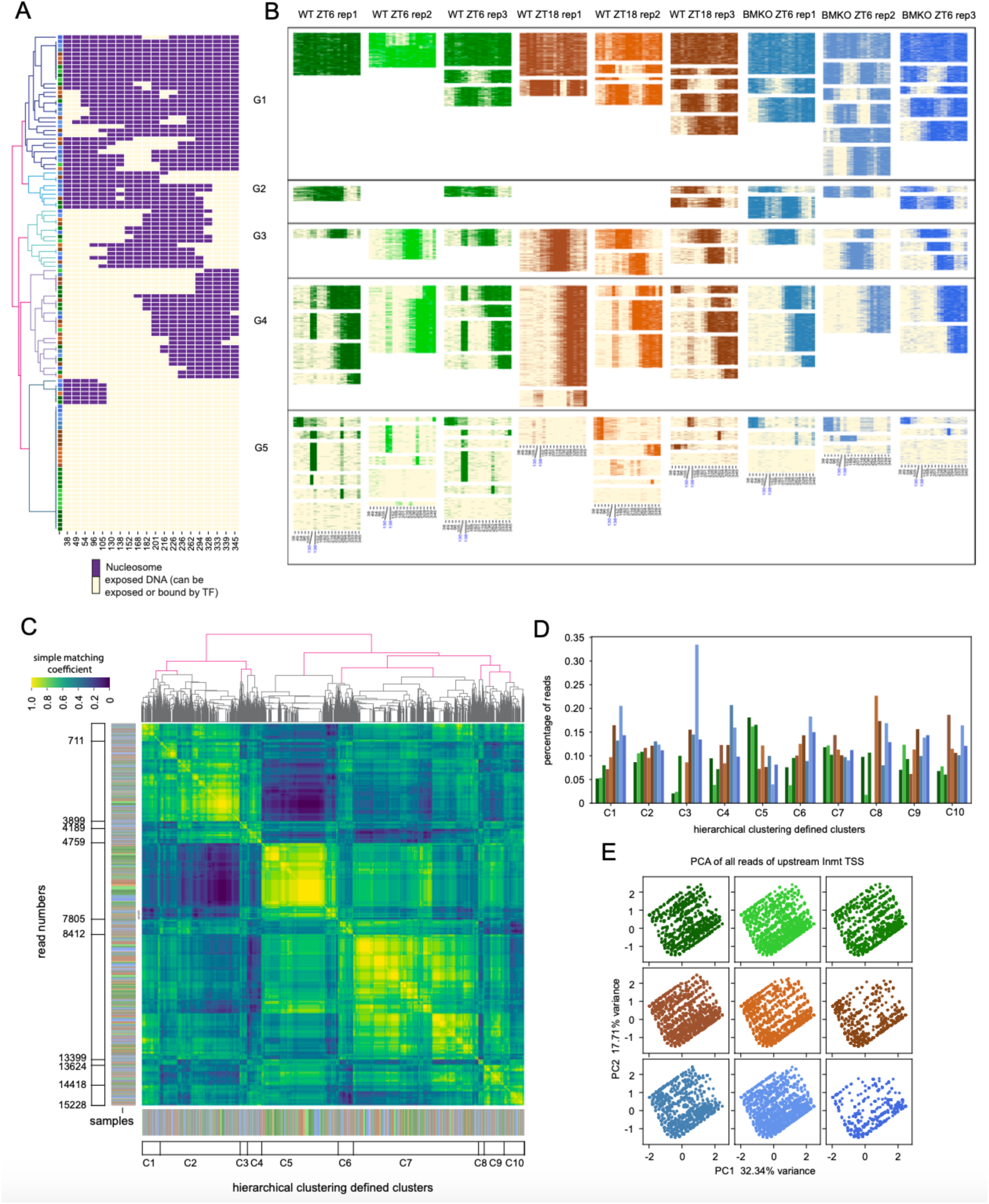
Analyses of conservation of chromatin states between samples. **(A)** BMD clustering analysis was performed separately for each individual sample (n=9) at *Inmt* TSS upstream enhancer, using the same set of reads as in Figure 1C. GpC protection was then computationally defined for every cluster as either nucleosome protected (purple) or exposed chromatin (bound or not by a TF; yellow), and this information was used to carry out a hierarchical clustering. The color of each square at the end of the dendrogram represents the biological source of the cluster using the same color code as in Figure 1C for each of the nine animals. Branches on the left of the dendrogram represent the five major hierarchical clusters. **(B)** After performing BMD clustering separately for each individual sample, SMF clusters were grouped based on the five major hierarchical clusters found in (a). **(C)** Heatmap of the simple matching coefficient calculated using GpC protection information at *Inmt* TSS upstream enhancer from a total of 15,228 reads (n = 1,692 reads per sample; n = 9 samples), and clustered by hierarchical clustering. The dendrogram at the top of the heatmap represents the hierarchical clusters. Colored lines on the outside left and bottom of the heatmap correspond to the biological source of each read (i.e., the originating biological sample) using the same color code as in Fig 1c, and the empty rectangles labelled C1 to C10 represent the 10 most different clusters as per the hierarchical clustering of the simple matching coefficient (SMC). **(D)** Bar plot representing the percentage of reads for each of the nine samples in each of the top 10 clusters obtained in panel (a) after hierarchical clustering of the SMC values. Most clusters contain reads from all nine samples (clusters C3 and C8, which are the two smallest clusters, show more variability). **(E)** Principal component analysis (PCA) of SMF signal at *Inmt* TSS upstream enhancer. Analysis was first performed using reads from all nine mouse livers, and then parsed for each sample. Each dot represents a DNA molecule, and individuals are represented by a single color as in Fig 1c. The distribution of the reads (dots) based on PC1 and PC2, which account for almost 50% of the variance, appears similar for all nine biological samples.

**Figure S5.**
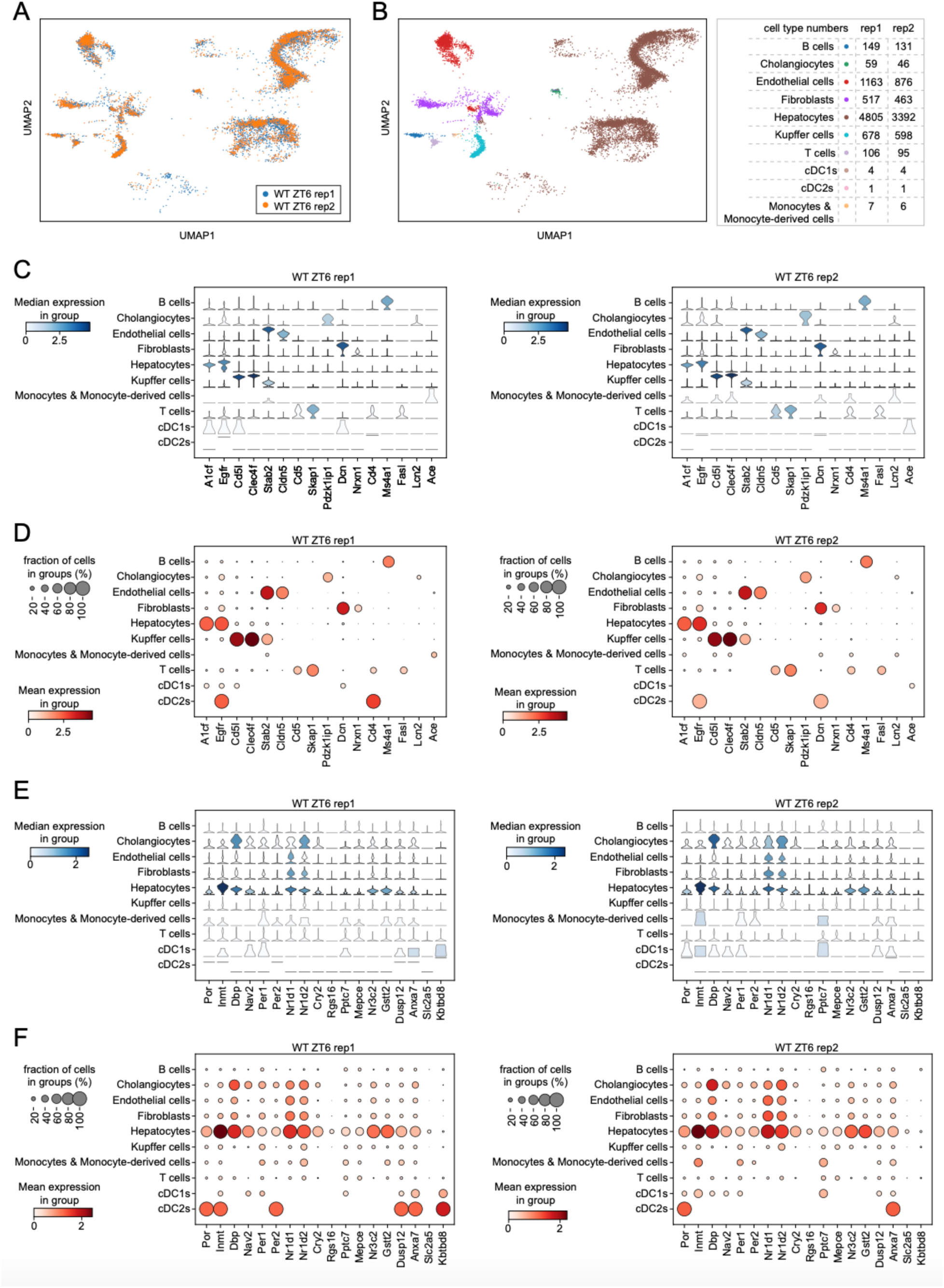
Single nuclei RNA-seq from two mouse livers. **(A-B)** UMAP of RNA expression from pre-processed snRNA-seq data from two mouse liver samples. A: UMAP color-coded by sample, demonstrating minimal batch effects. B: UMAP visualization of the same data, color-coded by 11 cell type clusters identified after transferring labels from a published liver cell atlas (LCA) dataset^78^. **(C, E)** Violin plots of the expression of cell-type specific marker genes (panel C) and genes targeted by CREs included in this paper (panel E) in each cell type. The color bar displayed the median expression. **(D, F)** Dot plots showing the expression of cell-type specific marker genes (panel D) and genes targeted by CREs included in this paper (panel F) in each cell type. The color bar represents the mean expression. The size of each dot represents the fraction of cells expressing the target genes.

**Figure S6.**
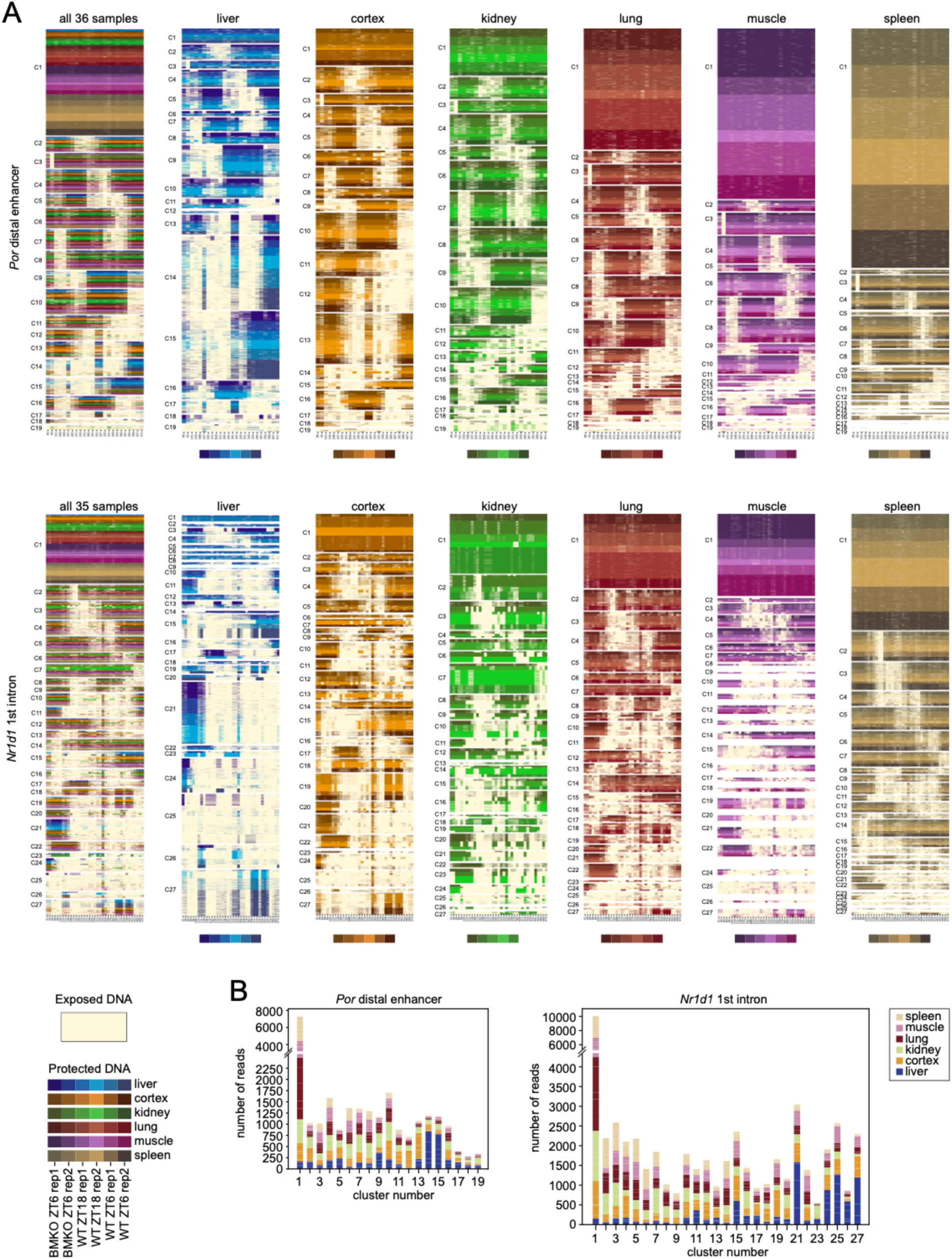
CREs exhibit stereotypic chromatin conformations across tissues. **(A)** SMF profile at *Por* distal enhancer (chr5:135703727-135704053) (top) and *Nr1d1* 1^st^ intron (chr11:98664610-98665043) (bottom) in 6 tissues, including liver, cortex, kidney, lung, muscle and spleen that were collected from three groups of mice (WT ZT06, WT ZT18 and BMKO ZT06; 2 replicates of each). The same number of reads was collected from each sample and were clustered by BMD clustering analysis. The heatmap on the very left contains reads from all samples (36 for *Por* distal enhancer and 35 for *Nr1d1* 1^st^ intron). Then, the heatmaps from left to right represent SMF data from the liver, cortex, kidney, lung, muscle, and spleen, with each heatmap containing reads from three groups. Color coding for protected reads in each sample are illustrated at the bottom. **(B)** Number of reads by each tissue in each cluster for *Por* distal enhancer (left) and *Nr1d1* 1^st^ intron (right).

**Figure S7.**
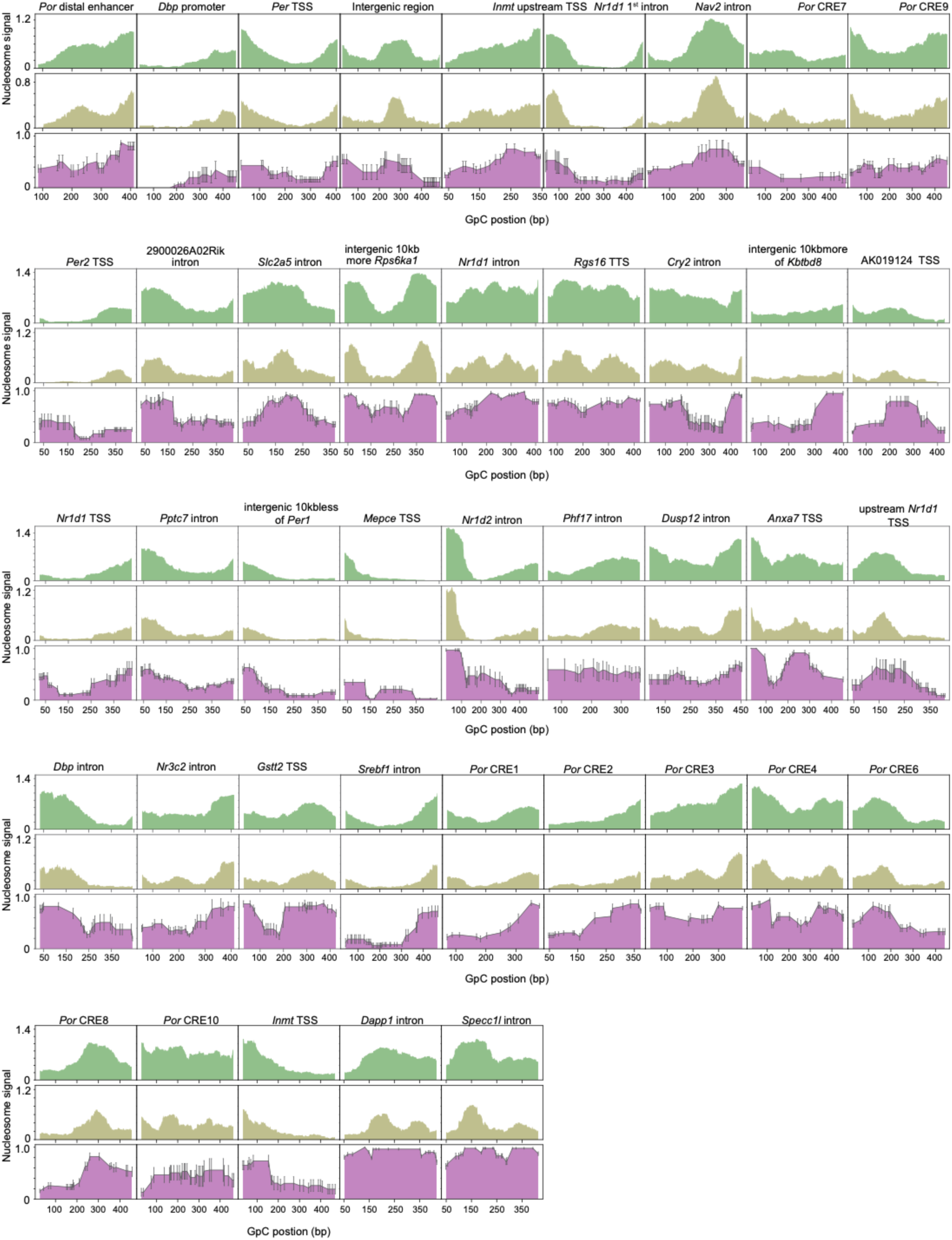
Comparison of SMF nucleosome signal with MNase-Seq signal. Nucleosome signals at the 41CREs investigated in this study. Top and middle panels: MNase-seq signals displayed with nucleosome signal calculated for the full-length 147 bp nucleosome (top) or calculated for 75 bp centered on the nucleosome dyad (middle; commonly used to improve the resolution of nucleosome positioning by MNase-Seq). Bottom panel: SMF nucleosome signal. MNase-Seq data was retrieved and reanalyzed from a public dataset (GSE47145; Menet et al., 2014^30^) representing almost 1 billion nucleosomes from 47 different mouse liver samples.

**Figure S8.**
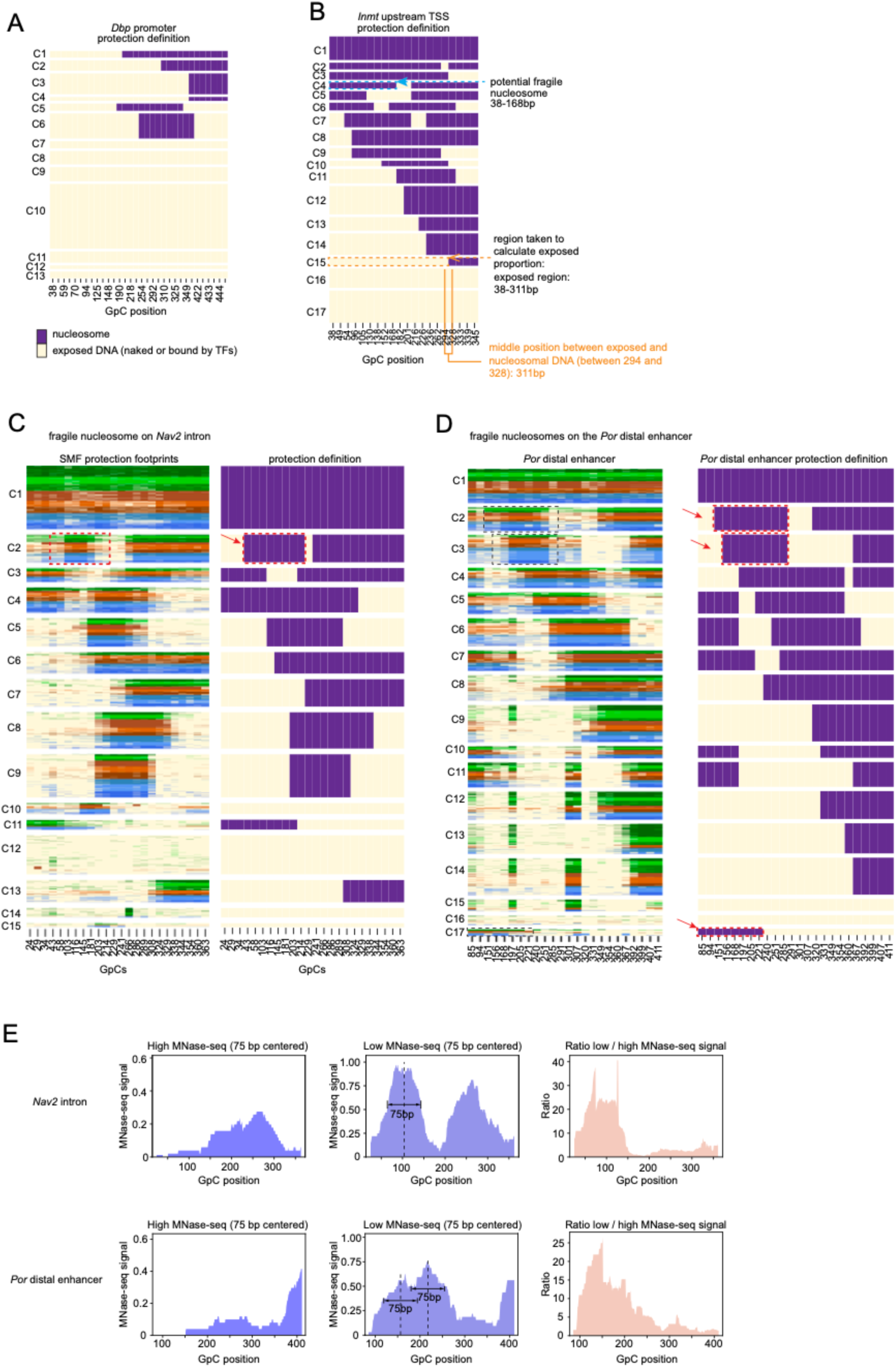
SMF detects fragile nucleosomes signal with MNase-Seq signal. **(A, B)** Computationally defined protection event at each GpC and for each cluster at *Dbp* promoter (A) and *Inmt* TSS upstream enhancer (B), with nucleosome signal colored in purple and DNA bound by TF or exposed in yellow. The algorithm used to computationally define nucleosome from SMF data is described in the method section. The nucleosome at GpCs 38-168 in C4 of *Inmt* TSS upstream enhancer (panel B) was characterized as a potential fragile nucleosome, see main text for details. This computationally defined nucleosome position was used to calculate the exposed level of a CRE. Specifically, the ratio of exposed DNA was first calculated for each cluster by dividing the length of exposed DNA for each cluster (e.g., 38 to 311 bp in C15 of *Inmt* TSS upstream enhancer) by the full length of the CRE (e.g., 345 - 38 = 307 bp for *Inmt* TSS upstream enhancer). The proportion of exposed DNA for each cluster was then multiplied by the percentage of reads of each cluster, which were then summed to return the final percentage of exposed DNA at a specific CRE. **(C)** Left panel: SMF profile at *Nav2* intron (chr7:49001617-49001956) in mouse liver. The heatmap displays protection from GpC methylation as in Figure 1C (n = 1761 reads for each sample, range of 340bp). Right panel: Computationally defined protection event at each GpC and for each cluster at *Nav2* intron. The box from GpCs 34-214 in C2 outlines a potential fragile nucleosome. **(D)** Left panel: SMF profile at *Por* distal enhancer (chr5:135703727-135704053) in mouse liver. The heatmap displays protection from GpC methylation as in Figure 1C (n = 1,833 reads for each sample; range of 327 bp). Right panel: Computationally defined protection event at each GpC and for each cluster at *Por* distal enhancer. The boxes from GpCs 151-285 in C2, 156-285 in C3, and 85-221 in C17 outline potential fragile nucleosomes. **(E)** MNase-seq nucleosome signals at *Nav2* intron (top) and *Por* distal enhancer (bottom), obtained from paired-end sequencing of high (left) and low (middle) MNase digested mouse liver nuclei. The ratio of low over high MNase-seq signal (right) was calculated to represent nucleosome fragility at *Por* distal enhancer.

**Figure S9:**
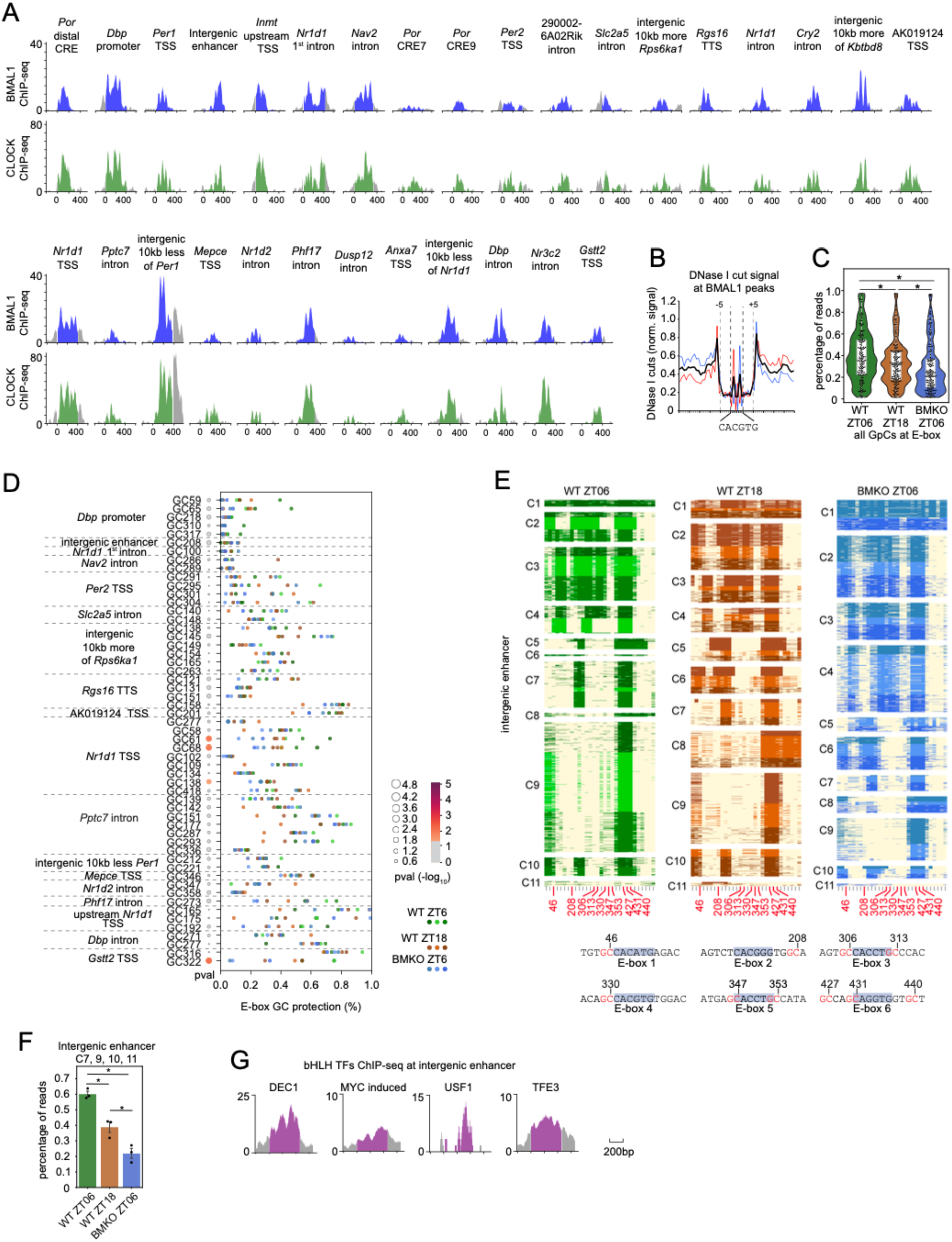
Analysis of DNA protection at E-boxes. **(A)** BMAL1 ChIP-seq (top, blue), and CLOCK ChIP-seq (bottom, green) at all 30 CREs bound by CLOCK:BMAL1. **(B)** DNase I cut signal centered on E-boxes (CACGTG only) at CLOCK:BMAL1 peaks. Signal corresponds to the average of DNase I cuts at a total of 1,912 E-boxes located within 3217 CLOCK:BMAL1 peaks (the list of peaks can be found in Trott and Menet, 2018^29^). The mouse liver DNase-Seq dataset was retrieved from the Encode project. **(C)** Percentage of protection at 158 GpCs located within 5 bp of E-boxes (CACGTG with up to one mismatch) for the 30 CREs bound by CLOCK:BMAL1 in WT ZT06 (green), WT ZT18 (brown), and BMKO ZT06 (blue). The percentage of protection was calculated using reads where GpCs were not protected by a nucleosome. Repeated measures ANOVA between the three groups. * p&<0.05. **(D)** Scatter plot of the percentage of protection at 55 degenerate E-boxes (not CACNTG or CANGTG) GpCs for all 9 samples, using Fig. 1C color code. The size and color of the bubble plot (left) depict the p-value obtained by one-way ANOVA (n = 3 samples/group). Circles are colored in grey if p>0.05. **(E)** Heatmaps illustrating the SMF profile at an intergenic enhancer region (chr17:29615647-29616108) in mouse liver. Reads from nine samples (533 reads for each sample) were clustered by BMD clustering, and parsed by group. Each heatmap contains reads from three replicates that are shaded in different colors of green, brown, and blue. Colors are as in Fig. 1c. **(F)** Percentage of reads in clusters C7, C9, C10, and C11 of the intergenic enhancer described in panel c. The bar graph represents the average +/- SEM of three biological replicates, and the dots correspond to the individual value for each of the nine sample. One-way ANOVA was performed between the three groups (p=2.8x10^-4^). **(G)** ChIP-seq of DEC1 (aka BHLHE40), USF1, MYC and TFE3. The purple areas are corresponding to the region as plotted in the heatmap.

**Figure S10.**
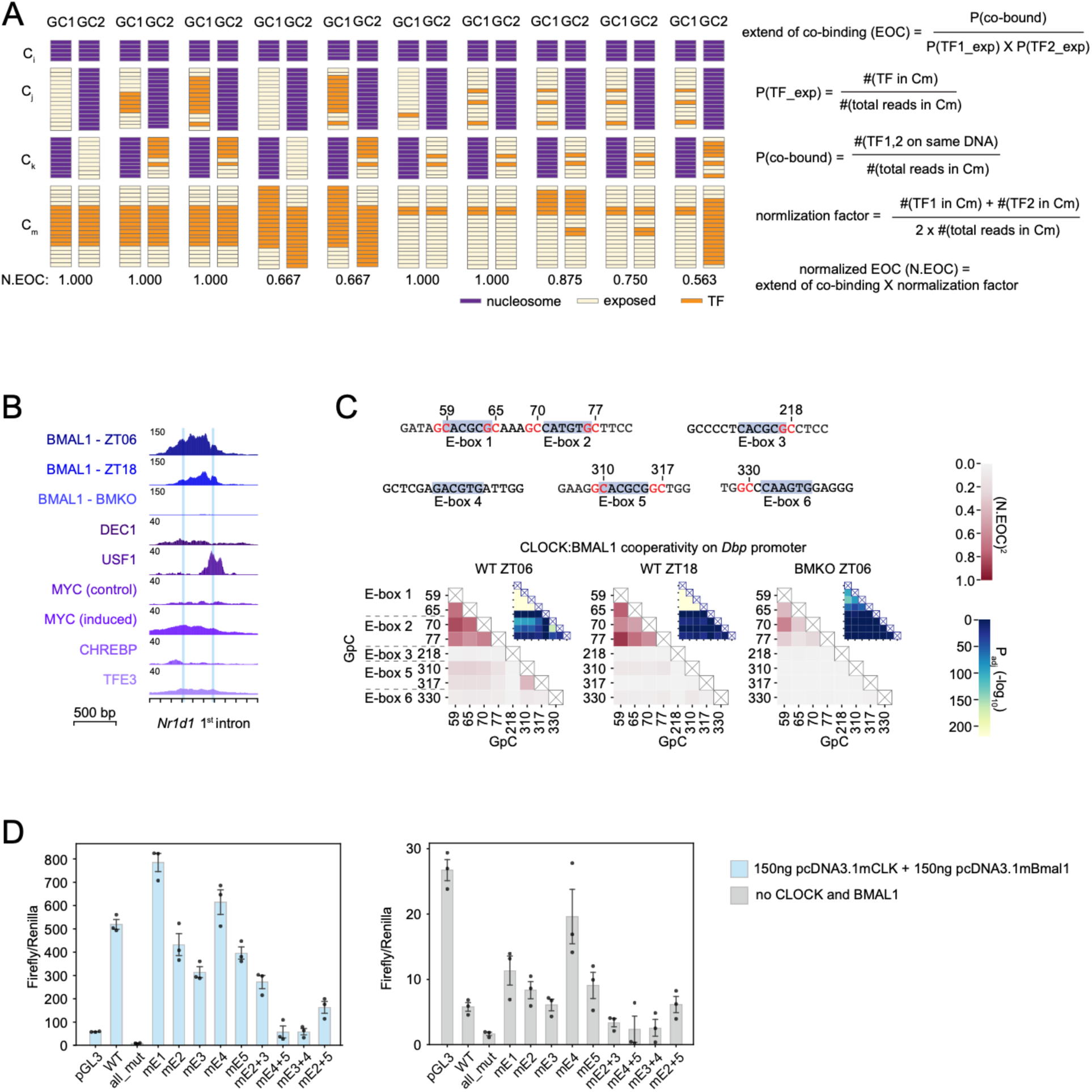
Long-range CLOCK:BMAL cooperative binding. **(A)** Formulas used for the calculation of the normalized extend of co-binding (N.EOC), which were adapted from Rao et al., 2021^5^. Panels on the left illustrate 10 different scenarios of co-binding events between two GpCs (GpC 1 and GpC 2, labeled GC1 and GC2, respectively) along with the associated N.EOC. **(B)** Liver BMAL1 CUT&RUN signals at *Nr1d1* 1^st^ intron in WT ZT6, WT ZT18, and BMKO ZT6. ChIP-seq signals of 5 bHLH TFs at *Nr1d1* 1^st^ intron. **(C)** N.EOC calculated between every pair of GpCs located in an E-box at *Dbp* promoter. Squared N.EOC values were plotted in heatmap. Sequences on the left correspond to the 5 E-boxes and their associated 8 GpCs. Fisher’s exact test was performed between each pair of E-box GpCs, followed by Benjamini-Hochberg correction. **(D)** Luciferase reporter assays using a CRE located in *Nr1d1* 1^st^ intron or variants containing different combinations of E-box(es) mutations. pGL3 served as a negative control. Transcriptional activity was calculated as the ratio firefly/renilla luciferase for all 12 reporters, and measured in the absence (right) or in the presence (left) of *Clock* and *Bmal1* expression vectors (150 ng each).

**Figure S11.**
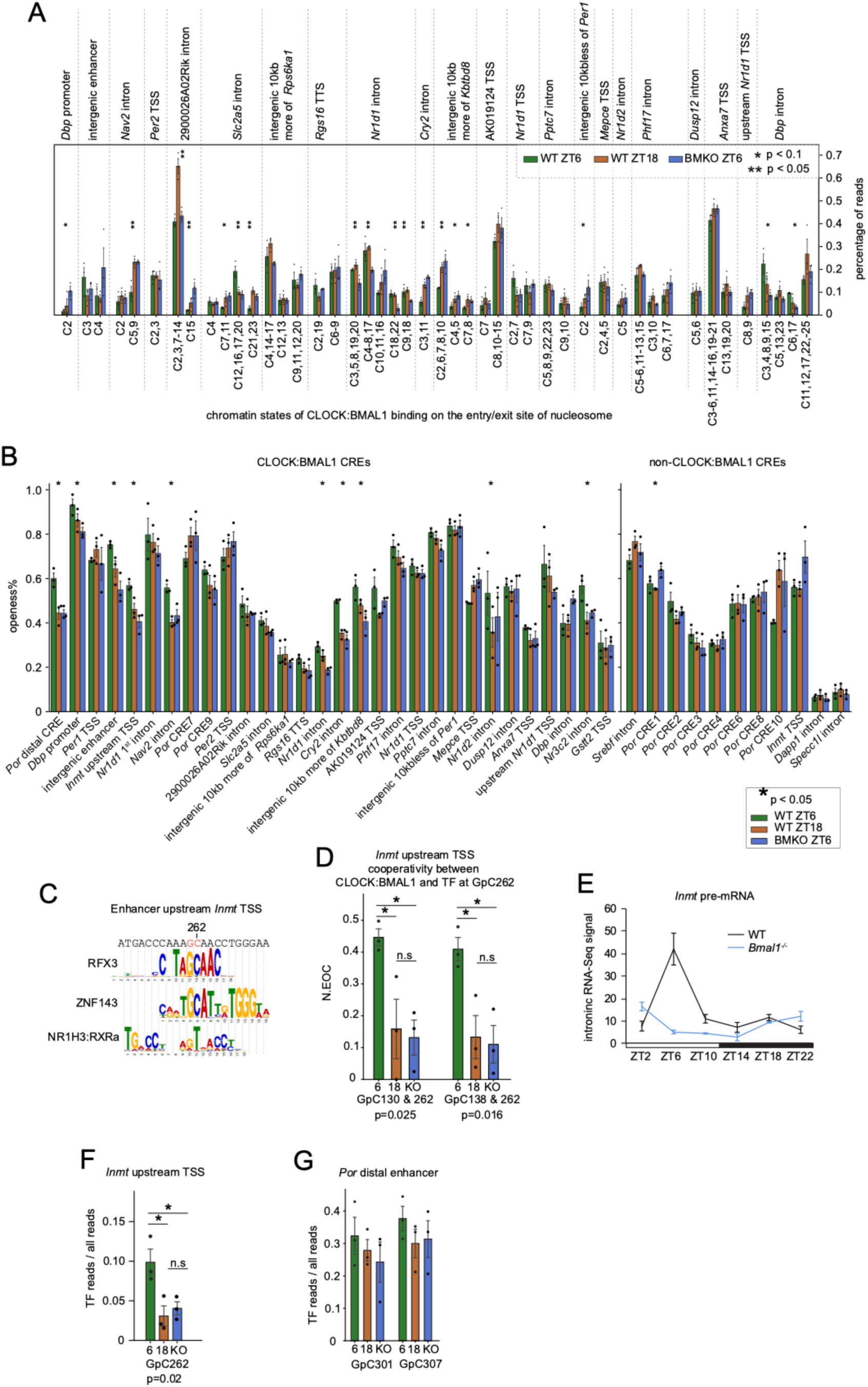
Cooperation between CLOCK:BMAL1 and other TFs. **(A)** Percentage of reads in clusters identified as containing a nucleosome with E-box(es) at the entry/exit site. These E-box motifs include variants that are not CACNTG or CANGTG motifs with GpCs that were not significant among the three groups. Error bars correspond to the S.E.M. of 3 biological replicates. One-way ANOVA; * p&<0.1; ** p&<0.05. Clusters with nucleosomes at similar locations were combined. **(B)** Percentage of exposed DNA at CREs in the three groups (WT ZT06, WT ZT18 and BMKO ZT06). Bar graphs were separated based on whether the CRE is bound by CLOCK:BMAL1 (left panel) or not (right panel), based on ChIP-seq data. **(C)** TF motif analysis around GpC 262 at *Inmt* TSS upstream enhancer performed using TomTom. **(D)** Normalized extent of co-binding between CLOCK:BMAL1 and the candidate TF at GpC 262 of *Inmt* TSS upstream enhancer. Each dot represents the value of each sample, and the error bars correspond to the S.E.M. of 3 biological replicates. **(E)** Mouse liver *Inmt* pre-mRNA level across the 24-hour day, from public mouse liver total RNA-Seq datasets (GSE73554) from Atger et al., 2015^81^. Black and blue line represent nighttime-fed WT and BMKO mice, respectively. **(F, G)** Percentage of protection at GpC 301, 307 at *Por* distal enhancer (F) and GpC 262 at *Inmt* TSS upstream enhancer (G). Error bars correspond to the S.E.M. of 3 biological replicates.

**Figure S12:**
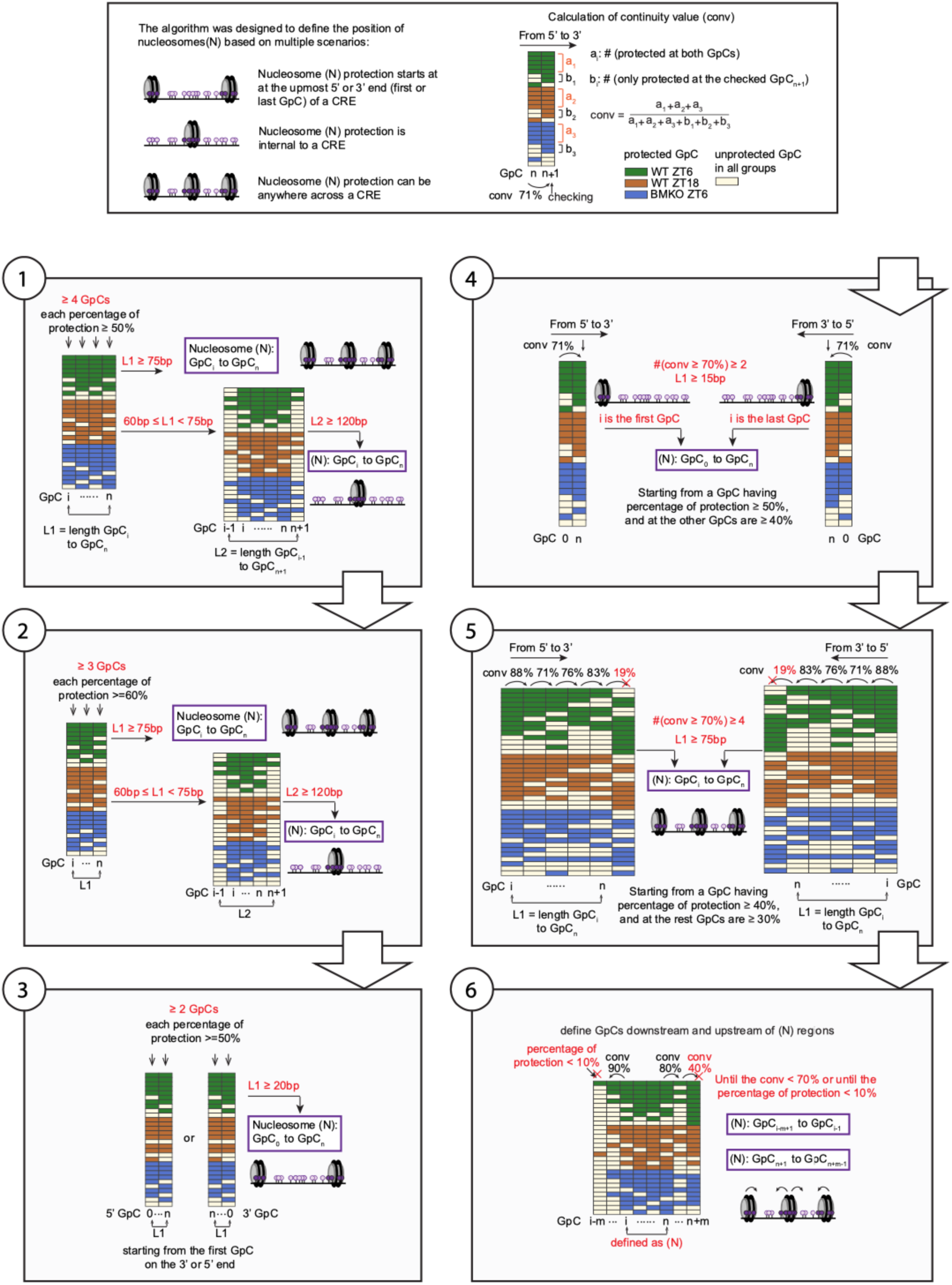
Schematic of computational definition of nucleosome protection. Schematic illustrating the logical workflow used in the algorithm to computationally define the position of nucleosomes using SMF data

## Notes

### Competing Interest Statement

The authors have declared no competing interest.

### Summary of Updates

The revised manuscript has been updated with more CREs and validated experiments, e.g luciferase assays to test TF cooperatively. More co-authors who helped with the experiments are updated in the revised manuscript.

