## Supplementary material for "Regulation of stereotypic chromatin conformations at enhancers by CLOCK:BMAL1": SMF_defineRegionCombinedSamples.py

```
import pandas as pd
import numpy as np
import sys
def defineRegionCombinedSamples(clustered,GCIndex):
    '''define TF(1), N(-1) taken sample together for each state (or cluster)
    can plot the GT% for each site for each cluster'''
    GT = [] # contains the percentage of GT at each site for each state.
    clusterNumber = len(clustered)
    clusterCombinedSample = []
    for st in range(clusterNumber):
        combined = clustered[st]
        GT_st = [] #GT percentage at each site.
        for site in range(len(GCIndex)):
            count = sum([combined[n][site] for n in range(len(combined))])
            GT_st.append(count/len(combined))
        clusterCombinedSample.append(combined)
        GT.append(GT_st)
    allReads = sum([len(clustered[n]) for n in range(clusterNumber)])

    condition = []
    # define >=3 consecutive >= 0.6 & range >= 75bp as N.
    # define site at head and tail >= 0.6 single site as N
    siteNumber = len(GCIndex)
    for st in range(clusterNumber):
        con = np.ones(siteNumber)
        percentage = GT[st]
        site = 0
        while site < siteNumber:
            rolling = 0
            while (site+rolling) < siteNumber and GT[st][site+rolling] >= 0.6:
                rolling += 1
            if rolling >= 3 and 1+GCIndex[site+rolling-1]-GCIndex[site] >= 75:
                con[site:site+rolling] = -1
            elif site > 0 and site + rolling < siteNumber and rolling >= 3 and 1+GCIndex[site+rolling-1]-GCIndex[site] >= 60:
                if 1+GCIndex[site+rolling]-GCIndex[site-1] >= 120:
                    con[site:site+rolling] = -1
            site = site + rolling + 1
        condition.append(con)
    # define >=4 consecutive >= 0.5 & range >= 75bp as N.
    for st in range(clusterNumber):
        percentage = GT[st]
        site = 0
        while site < siteNumber:
            rolling = 0
            while (site+rolling) < siteNumber and GT[st][site+rolling] >= 0.5:
                rolling += 1
            if rolling >= 4 and 1+GCIndex[site+rolling-1]-GCIndex[site] >= 75:
                condition[st][site:site+rolling] = -1
            elif site > 0 and site + rolling < siteNumber and rolling >= 4 and 1+GCIndex[site+rolling-1]-GCIndex[site] >= 60:
                if 1+GCIndex[site+rolling]-GCIndex[site-1] >= 120:
                    condition[st][site:site+rolling] = -1
            site = site + rolling + 1

    #if the edge 20bp are >60%, we call it as nucleosome
    for st in range(clusterNumber):
        site = 0
        while site < siteNumber and GT[st][site] >= 0.5:
            site+=1
        if site > 0 and 1 + GCIndex[site-1]-GCIndex[0] >= 20:
            condition[st][:site] = -1
        site = -1
        while site >= -siteNumber and GT[st][site] >= 0.5:
            site-=1
        if site < -1 and 1 + GCIndex[-1] - GCIndex[site+1] >= 20:
            condition[st][site+1:] = -1
    # consecutive 1 on the edge. (70%  (continuity_value) of 1 with a 1 at the previous site). This is to check the edge nucleosome only
    # continuity_value is 0.7
    for st in range(clusterNumber):
        state = clusterCombinedSample[st]
        site = 0
        consecutivePer = 0
        fuzzy = [[state[seq][site],state[seq][site+1]] for seq in range(len(state))]
        checking = [state[seq][site+1] for seq in range(len(state)) if state[seq][site+1] == 1]
        if len(checking) > 0:
            consecutivePer = fuzzy.count([1,1])/len(checking)
        while site < siteNumber-1 and GT[st][site] >= 0.5 and GT[st][site+1] >= 0.4 and consecutivePer >= 0.7:
            fuzzy = [[state[seq][site],state[seq][site+1]] for seq in range(len(state))]
            checking = [state[seq][site+1] for seq in range(len(state)) if state[seq][site+1] == 1]
            if len(checking) > 0:
                consecutivePer = fuzzy.count([1,1])/len(checking)
            else:
                consecutivePer = 0
            if consecutivePer>=0.7:
                site += 1
        if site > 0 and 1 + GCIndex[site]-GCIndex[0] >= 15:
            condition[st][:site+1] = -1
        site = -1
        consecutivePer = 0
        fuzzy = [[state[seq][site],state[seq][site-1]] for seq in range(len(state))]
        checking = [state[seq][site-1] for seq in range(len(state)) if state[seq][site-1] == 1]
        if len(checking) > 0:
            consecutivePer = fuzzy.count([1,1])/len(checking)
        while site > -siteNumber and GT[st][site] >= 0.5 and GT[st][site-1] >= 0.4 and consecutivePer >= 0.7:
            fuzzy = [[state[seq][site],state[seq][site-1]] for seq in range(len(state))]
            checking = [state[seq][site-1] for seq in range(len(state)) if state[seq][site-1] == 1]
            if len(checking) > 0:
                consecutivePer = fuzzy.count([1,1])/len(checking)
            else:
                consecutivePer = 0
            if consecutivePer >= 0.7:
                site -= 1
        if site < -1 and 1 + GCIndex[-1] - GCIndex[site] >= 15:
            condition[st][site:] = -1

    # five consecutive 1 are N. (70% of 1 with a 1 at the previous site)
    for st in range(clusterNumber):
        site = 0
        state = clusterCombinedSample[st]
        while site <= siteNumber-5:
            rolling = 0
            consecutivePer = 0
            if GT[st][site] >= 0.4 and condition[st][site] != -1 and GT[st][site+1] >= 0.3:
                fuzzy = [[state[seq][site],state[seq][site+1]] for seq in range(len(state))]
                checking = [state[seq][site+1] for seq in range(len(state)) if state[seq][site+1] == 1]
                if len(checking) > 0:
                    consecutivePer = fuzzy.count([1,1])/len(checking)
            while consecutivePer >= 0.7 and site + rolling < siteNumber-1 and GT[st][site+rolling+1] >= 0.3:
                fuzzy = [[state[seq][site+rolling],state[seq][site+rolling+1]] for seq in range(len(state))]
                checking = [state[seq][site+rolling+1] for seq in range(len(state)) if state[seq][site+rolling+1] == 1]
                if len(checking) > 0:
                    consecutivePer = fuzzy.count([1,1])/len(checking)
                else:
                    consecutivePer = 0
                if consecutivePer >= 0.7:
                    rolling = rolling + 1
            if rolling >= 4 and 1+GCIndex[site+rolling]-GCIndex[site] >= 75:
                condition[st][site:site+rolling+1] = -1

            site = site + rolling + 1
    for st in range(clusterNumber):
        site = siteNumber - 1
        state = clusterCombinedSample[st]
        while site >= 4:
            rolling = 0
            consecutivePer = 0
            if GT[st][site] >= 0.4 and condition[st][site] != -1 and GT[st][site-1] >= 0.3:
                fuzzy = [[state[seq][site-1],state[seq][site]] for seq in range(len(state))]
                checking = [state[seq][site-1] for seq in range(len(state)) if state[seq][site-1] == 1]
                if len(checking)>0:
                    consecutivePer = fuzzy.count([1,1])/len(checking)
            while consecutivePer >= 0.7 and site - rolling > 0  and GT[st][site-rolling-1] >= 0.3:
                fuzzy = [[state[seq][site-rolling-1],state[seq][site-rolling]] for seq in range(len(state))]
                checking = [state[seq][site-rolling-1] for seq in range(len(state)) if state[seq][site-rolling-1] == 1]
                if len(checking) > 0:
                    consecutivePer = fuzzy.count([1,1])/len(checking)
                else:
                    consecutivePer = 0
                if consecutivePer >= 0.7:
                    rolling = rolling + 1
            if rolling >= 4 and 1+GCIndex[site]-GCIndex[site-rolling] >= 75:
                condition[st][site-rolling:site+1] = -1

            site = site - rolling - 1

    # check fuzzy nucleosome signal.
    for st in range(clusterNumber):
        con = condition[st]
        for c in range(len(con)):
            con = condition[st]
            if con[c] == -1:
                # check fuzzy
                state = clusterCombinedSample[st]
                if c >= 1 and con[c-1] != -1:
                    fuzzy = [[state[seq][c-1],state[seq][c]] for seq in range(len(state))]
                    checking = [state[seq][c-1] for seq in range(len(state)) if state[seq][c-1] == 1]
                    consecutivePer = 0
                    if len(checking) != 0 and GT[st][c-1]>=0.1:
                        consecutivePer = fuzzy.count([1,1])/len(checking)
                        if np.round(consecutivePer,2) >= 0.7:
                            condition[st][c-1] = -1
                    n = 1
                    while c-1-n >= 0 and consecutivePer >= 0.7:
                        fuzzy = [[state[seq][c-1-n],state[seq][c-n]] for seq in range(len(state))]
                        checking = [state[seq][c-1-n] for seq in range(len(state)) if state[seq][c-1-n] == 1]
                        if len(checking) != 0 and GT[st][c-1-n]>0.1:
                            consecutivePer = fuzzy.count([1,1])/len(checking)
                            if np.round(consecutivePer,2) >= 0.7:
                                condition[st][c-1-n] = -1
                        else:
                            consecutivePer = 0
                        n = n+1
                elif c <= len(con)-2 and con[c+1] != -1:
                    fuzzy = [[state[seq][c],state[seq][c+1]] for seq in range(len(state))]
                    checking = [state[seq][c+1] for seq in range(len(state)) if state[seq][c+1] == 1]
                    consecutivePer = 0
                    if len(checking) != 0 and GT[st][c+1]>0.1:
                        consecutivePer = fuzzy.count([1,1])/len(checking)
                        if np.round(consecutivePer,2) >= 0.7:
                            condition[st][c+1] = -1
                    n = 1
                    while c+1+n < len(con) and consecutivePer >= 0.7:
                        fuzzy = [[state[seq][c+n],state[seq][c+n+1]] for seq in range(len(state))]
                        checking = [state[seq][c+1+n] for seq in range(len(state)) if state[seq][c+1+n] == 1]
                        if len(checking) != 0 and GT[st][c+1+n]>0.1:
                            consecutivePer = fuzzy.count([1,1])/len(checking)
                            if np.round(consecutivePer,2) >= 0.7:
                                condition[st][c+1+n] = -1
                        else:
                            consecutivePer = 0
                        n = n+1
    return pd.DataFrame(data = condition, index = [n for n in range(0,len(condition))],
                             columns = GCIndex)

cmdargs = sys.argv
FILENAME = cmdargs[cmdargs.index('-f')+1] #processed file with clustered files
clusterdData = pd.read_csv(FILENAME, sep = '\t')
DHSnumber = np.unique(clusterdData['DHS_number'].values)
DHSnumber = [int(i) for i in DHSnumber]
try:
    e = cmdargs.index('-e') # the number has to match the number in your file (the DHS_number column).
    # for example (no space between comma): -e [1,2,3]
    enhancersToPlot = cmdargs[e+1]
    enhancersToPlot = enhancersToPlot[1:-1].split(',')
    enhancersToPlot = [int(i) for i in enhancersToPlot]
    print('will plot enhancer', enhancersToPlot)
except:
    enhancersToPlot = DHSnumber
    print('will plot all enhancers',enhancersToPlot)
try:
    ind = cmdargs.index('-o')
    outputFileName = cmdargs[ind+1]
except:
    outputFileName = FILENAME[:FILENAME.index('.clustered')]

GCIndex = []
TFconditionEachEnhancer = []
protectionPerEachEnhancer = []
for i in range(len(enhancersToPlot)):
    e = enhancersToPlot[i]
    sub = clusterdData[clusterdData['DHS_number']==e]
    name = sub['DHS_name'].values[0]
    gc = sub['GCIndex'].values[0]
    gc = gc[1:-1].split(',')
    gc = [int(m) for m in gc]
    GCIndex.append(gc)
    notSortedClstOrder = np.unique(sub['cluster_number'].values)
    clusterNumber = len(notSortedClstOrder)
    combineSampleEachEnClustered = []
    GT = [] # contains the percentage of GT at each site for each state.
    for st in range(clusterNumber):
        stdf = sub[sub['cluster_number'] == st]
        checked = stdf['checkGT'].values
        checkedEachSt = []
        for j in range(len(checked)):
            l = checked[j] # e.g: '[1,1,0,0,1]'
            l = l[1:-1].split(',')
            ll = [int(m) for m in l]
            checkedEachSt.append(ll)
        combineSampleEachEnClustered.append(checkedEachSt)
        GT_st = [] #GT percentage at each site.
        for site in range(len(gc)):
            count = sum([checkedEachSt[n][site] for n in range(len(checkedEachSt))])
            GT_st.append(count/len(checkedEachSt))
        GT.append(GT_st)
    TFcondition = defineRegionCombinedSamples(combineSampleEachEnClustered,gc)
    TFcondition.replace(0,-2,inplace=True) #there should be no 0 in the definotion. (here just in case)
    TFcondition.replace(1,0,inplace=True) #exposed
    TFcondition.replace(-1,1,inplace=True) #Nucleosome
    TFconditionEachEnhancer.append(TFcondition)
    protectionPerEachEnhancer.append(GT)
    indJ = 0
    for j in range(len(outputFileName)-1,-1,-1):
        if outputFileName[j] == '/':
            indJ = j
            break
    substring = outputFileName[indJ+1:]
    title = substring + 'DHS'+str(e)+'_'+name
    TFcondition.to_csv(outputFileName[:indJ+1]+title+'NuTFdefine.tsv',sep='\t') #in the output files, exposed:0, nucleosome:1
```
