## Supplementary material for "Regulation of stereotypic chromatin conformations at enhancers by CLOCK:BMAL1": SMFalignV3.py

```
import pandas as pd
import numpy as np
import random
from Bio.Align import PairwiseAligner
from collections import Counter
import sys

# this file can be submitted to terminal as the example below (R1fileName and R2fileName contains the fastq names to be aligned and have to be in the same folder):
# python SMFalignV3.py -r1 R1fileName.txt -r2 R2fileName.txt -p Primer_sequences.bed -t DHS_sequences.bed -o outputname &

def getReverseComplement (sequence):
    '''get reverse complement of atgc DNA sequences'''
    sequence_upper = sequence.upper()
    sequence_upper_reverse = sequence_upper[::-1]
    reverse_complement = ""
    for i in sequence_upper_reverse:
        if i == "T":
            reverse_complement = reverse_complement + "A"
        elif i == "A":
            reverse_complement = reverse_complement + "T"
        elif i == "G":
            reverse_complement = reverse_complement + "C"
        elif i == "C":
            reverse_complement = reverse_complement + "G"
        else:
            reverse_complement = reverse_complement + i
    return reverse_complement

def getRevPrimerRevComplement(reversePrimer):
    '''this function get the reverse compliment of reverse primer sequences.'''
    reversePrimer_revComplement = []
    for i in reversePrimer:
        reversePrimer_revComplement.append(getReverseComplement(i))
    return reversePrimer_revComplement

def getBisulfiteEnSeq(originalEnhancerSeq):
    ''' all single Cs are mutated to T. GC or CG changed to GY or YG.'''
    enhancerNumber = len(originalEnhancerSeq)
    newEnhancerSeq = [0]*enhancerNumber
    for i in range(enhancerNumber):
        newEnhancer = ''
        old = originalEnhancerSeq[i].upper()
        length = len(old)
        for j in range(0,length):
            if old[j] != 'C':
                newEnhancer = newEnhancer + old[j]
            else:
                if j < length - 1 and old[j+1] == 'G':
                    newEnhancer = newEnhancer + 'Y'
                elif j > 0 and old[j-1] == 'G':
                    newEnhancer = newEnhancer + 'Y'
                else:
                    newEnhancer = newEnhancer +'T'
        newEnhancerSeq[i] = newEnhancer
    return newEnhancerSeq

def getForwardReverseUMIList(forwardPrimer,reversePrimer):
    '''this function return a list of umi for all forward and reverse primers.'''
    UMI_F, UMI_R = [],[]
    for i in range(len(forwardPrimer)):
        f = forwardPrimer[i].upper()
        umif = '' # to collect umi seq
        for j in range(len(f)):
            if f[j] not in ['A','T','G','C']:
                umif+= f[j]
            if f[j+1] in ['A','T','G','C']:
                break
        r = reversePrimer[i].upper()
        umir = '' # to collect umi seq
        for j in range(len(r)):
            if r[j] not in ['A','T','G','C']:
                umir+= r[j]
            if r[j+1] in ['A','T','G','C']:
                break
        UMI_F.append(umif)
        UMI_R.append(umir)
    return [UMI_F,UMI_R]

def getQCnum(R1rawQC,R2rawQC):
    '''based on the AscII code, raw quality score are converted to lists of numbers'''
    animalNum = len(R1rawQC)
    R1_qc, R2_qc = [0]*animalNum, [0]*animalNum
    for i in range(0,animalNum):
        tempContent1 = R1rawQC[i]
        tempContent2 = R2rawQC[i]
        qc_1, qc_2  = [], []
        length = len(tempContent1)
        for j in range(length):
            content1 = tempContent1[j]
            content2 = tempContent2[j]
            score1 = []
            score2 = []
            for c in content1:
                score1.append(ord(c)-33)
            for c in content2:
                score2.append(ord(c)-33)
            qc_1.append(score1)
            qc_2.append(score2)
        R1_qc[i] = qc_1
        R2_qc[i] = qc_2
    return [R1_qc,R2_qc]

def Align(animalNum,R1_sequence,R2_sequence,R1_qc,R2_qc,forwardPrimer,reversePrimer,UMI_F,UMI_R,newEnhancerSeq):
    # using Align.pairwiseAligner to compare primer with the beginning len(primer)+4nt.
    # Then compare target enhancer to result (whole sequence).
    # included collecting qc score and keep nt with higher qc.
    # the newest way to align
    enhancerNumber = [0]*animalNum #contains number of which enhancer that sequence is
    allFullSeq = [0]*animalNum
    allWholeQC = [0]*animalNum
    aligner = PairwiseAligner(match_score=1.0)
    aligner.mismatch_score = - 0.2
    aligner.gap_score = -0.5
    pair1alignment_score = [0]*animalNum #this score is pairwiseAligner score
    pair2alignment_score = [0]*animalNum
    pair1_alignment = [0]*animalNum # forward sequence
    pair2_alignment = [0]*animalNum # reverse sequence
    pair1_qualityScore = [0]*animalNum
    pair2_qualityScore = [0]*animalNum

    for sample in range(0,animalNum):
        # # pair2 should not be the seq after taking reverse complement.
        #for loop to go through all sequences in file 1 and 2.
        sequence_1 = R1_sequence[sample]
        sequence_2 = R2_sequence[sample]
        sequence1Length = len(sequence_1)
        whichEnhancer = [-100]*sequence1Length
        pair1_score = [0]*sequence1Length
        pair2_score = [0]*sequence1Length
        # whichEnhancer: the order is corrspond to the sequence order,
        # with enhancer number in this list (1-9). -100 indicates a trashed sequence
        fullSeq = ['NA']*sequence1Length
        p1_aligned = [0]*sequence1Length
        p2_aligned = [0]*sequence1Length
        wholeQC = [0]*sequence1Length # the quality score of the full length sequence
        P1QC = [0]*sequence1Length
        P2QC = [0]*sequence1Length
        for s in range(sequence1Length):
            pair1 = sequence_1[s]
            pair2 = sequence_2[s]
            qc1 = R1_qc[sample][s]
            qc2 = R2_qc[sample][s]
            for i in range(len(newEnhancerSeq)):
                c, cc = 0, 0 # the alignment score that matches the forward & reverse primer.
                n, nn = 0, 0 # the alignment score of the pair1 & pair2 read
                pair1reverse = False
                umi_f = len(UMI_F[i])
                umi_r = len(UMI_R[i])
                p = forwardPrimer[i][umi_f:]
                sub = pair1[umi_f:len(p)+umi_f+2]
                c = aligner.score(p, sub)/(len(p))
                if c < 0.8:
                    # meaning pair1 can be reverse as well
                    c = 0
                    revP = reversePrimer[i][umi_r:]
                    sub_rev = pair1[umi_r : len(revP)+umi_r+2]
                    c = aligner.score(revP, sub_rev)/(len(revP))
                    if c >= 0.8:
                        pair1reverse = True
                # if one is forward the other pair is reverse.
                # if one is reverse the other pair is forward.
                if c >= 0.8 and pair1reverse == False:
                    revP = reversePrimer[i][umi_r:]
                    pair2Sub = pair2[umi_r:len(revP)+umi_r+2]
                    cc = aligner.score(revP, pair2Sub)/(len(revP))
                elif c >= 0.8 and pair1reverse == True:
                    p = forwardPrimer[i][umi_f:]
                    pair2Sub = pair2[umi_f:len(p)+umi_f+2]
                    cc = aligner.score(p, pair2Sub)/(len(p))
                # compare the full sequence of each pair with correct sequences. UMI is removed when aligning.
                alignments1 = 0
                alignments2 = 0
                if cc >= 0.8 and pair1reverse == False:
                    pair2 = getReverseComplement(pair2[umi_r:])
                    pair1 = pair1[umi_f:]
                    qc2 = qc2[umi_r:]
                    qc1 = qc1[umi_f:]
                    qc2.reverse()
                    correctSeq1 = newEnhancerSeq[i][:len(pair1)]
                    correctSeq2 = newEnhancerSeq[i][len(newEnhancerSeq[i])-len(pair2):]
                    alignments1 = aligner.align(correctSeq1,pair1)
                    alignments2 = aligner.align(correctSeq2,pair2)
                    n = alignments1.score/len(pair1)
                    nn = alignments2.score/len(pair2)
                    pair1_score[s] = n
                    pair2_score[s] = nn
                    whichEnhancer[s] = -(i+1) #this is to mark enhancer 1 as -1 but not 0
                elif cc >= 0.8 and pair1reverse == True:
                    pair1 = getReverseComplement(pair1[umi_r:])
                    pair2 = pair2[umi_f:]
                    qc1 = qc1[umi_r:]
                    qc2 = qc2[umi_f:]
                    qc1.reverse()
                    correctSeq1 = newEnhancerSeq[i][len(newEnhancerSeq[i])-len(pair1):]
                    correctSeq2 = newEnhancerSeq[i][:len(pair2)]
                    alignments1 = aligner.align(correctSeq1,pair1)
                    alignments2 = aligner.align(correctSeq2,pair2)
                    n = alignments1.score/len(pair1)
                    nn = alignments2.score/len(pair2)
                    pair1_score[s] = n
                    pair2_score[s] = nn
                    whichEnhancer[s] = -(i+1) #this is to mark DHS 1 as -1 but not 0

                if n >= 0.7 and nn >= 0.7:
                    whichEnhancer[s] = i+1  #to mark which DHS (DHS number starts from  1)
                    alignment1 = alignments1[0].aligned
                    alignment2 = alignments2[0].aligned

                    query_aligned1 = alignment1[1] # all tuples
                    query_aligned2 = alignment2[1] # all tuples
                    target_aligned1 = alignment1[0] # all tuples
                    target_aligned2 = alignment2[0] # all tuples

                    # to reconstruct pair1
                    query_tuple1 = query_aligned1[len(query_aligned1)-1] # the last query tuple
                    target_tuple1 = target_aligned1[len(target_aligned1)-1] # the last target tuple
                    start1 = query_tuple1[0]
                    end1 = query_tuple1[1]
                    target_start1 = target_tuple1[0]
                    target_end1 = target_tuple1[1]
                    reconstructedPair1 = pair1[start1:end1]
                    reconQC1 = qc1[start1:end1]
                    if len(query_aligned1)==1 and target_start1 > start1:
                        reconstructedPair1 = 'I'*(target_start1-start1) + reconstructedPair1
                        reconQC1 = [0]*(target_start1-start1) +  reconQC1
                    for tup in range(len(query_aligned1)-2,-1,-1):
                        query_tuple = query_aligned1[tup]
                        newstart = query_tuple[0]
                        newend = query_tuple[1]
                        target_tuple = target_aligned1[tup]
                        target_newstart = target_tuple[0]
                        target_newend = target_tuple[1]
                        if start1 > newend: # to remove insertion in result
                            reconstructedPair1 = pair1[newstart:newend] + reconstructedPair1
                            reconQC1 = qc1[newstart:newend] + reconQC1
                        elif target_newend < target_start1: # to fill in the gap
                            reconstructedPair1 = pair1[newstart:newend] + 'I'*(target_start1-target_newend) + reconstructedPair1
                            reconQC1 = qc1[newstart:newend] + [0]*(target_start1-target_newend) + reconQC1
                        if tup == 0 and target_newstart > newstart and pair1reverse == False:
                            reconstructedPair1 = 'I'*(target_newstart-newstart) + reconstructedPair1
                            reconQC1 = [0]*(target_newstart-newstart) + reconQC1
                        start1 = newstart
                        end1 = newend
                        target_start1 = target_newstart
                        target_end1 = target_newend
                    # to reconstruct pair2
                    query_tuple2 = query_aligned2[len(query_aligned2)-1] # the last query tuple
                    target_tuple2 = target_aligned2[len(target_aligned2)-1] # the last target tuple
                    start2 = query_tuple2[0]
                    end2 = query_tuple2[1]
                    target_start2 = target_tuple2[0]
                    target_end2 = target_tuple2[1]
                    reconstructedPair2 = pair2[start2:end2]
                    reconQC2 = qc2[start2:end2]
                    if len(query_aligned2)==1 and target_start2 > start2:
                        reconstructedPair2 = 'I'*(target_start2-start2) + reconstructedPair2
                        reconQC2 = [0]*(target_start2-start2) +  reconQC2
                    for tup in range(len(query_aligned2)-2,-1,-1):
                        query_tuple = query_aligned2[tup]
                        newstart = query_tuple[0]
                        newend = query_tuple[1]
                        target_tuple = target_aligned2[tup]
                        target_newstart = target_tuple[0]
                        target_newend = target_tuple[1]
                        if start2 > newend: # to remove insertion in result
                            reconstructedPair2 = pair2[newstart:newend] + reconstructedPair2
                            reconQC2 = qc2[newstart:newend] + reconQC2
                        elif target_newend < target_start2: # to fill in the gap
                            reconstructedPair2 = pair2[newstart:newend] + 'I'*(target_start2-target_newend) + reconstructedPair2
                            reconQC2 = qc2[newstart:newend] + [0]*(target_start2-target_newend) + reconQC2
                        if tup == 0 and target_newstart > newstart and pair1reverse == True:
                            reconstructedPair2 = 'I'*(target_newstart-newstart) + reconstructedPair2
                            reconQC2 = [0]*(target_newstart-newstart) + reconQC2
                        start2 = newstart
                        end2 = newend
                        target_start2 = target_newstart
                        target_end2 = target_newend
                    # to combine full sequence
                    overlap = len(reconstructedPair1) + len(reconstructedPair2) - len(newEnhancerSeq[i])
                    # the overlapping number. The recon seq doesn't have umi at this stage
                    full = ""
                    # the umi seqence and the QC of umi is add back to reconsitituted sequence and reconQC.
                    if pair1reverse == False:
                        pair1umiSeq = sequence_1[s][:umi_f]
                        pair2umiSeq = getReverseComplement(sequence_2[s][:umi_r])
                        reconstructedPair1 = pair1umiSeq + reconstructedPair1
                        reconstructedPair2 = reconstructedPair2 + pair2umiSeq
                        umiQC1 = R1_qc[sample][s][:umi_f]
                        umiQC2 = R2_qc[sample][s][:umi_r]
                        reconQC1 = umiQC1 + reconQC1
                        reconQC2 = reconQC2 + umiQC2[::-1]
                        p1_aligned[s] = reconstructedPair1
                        p2_aligned[s] = reconstructedPair2
                        P1QC[s] = reconQC1
                        P2QC[s] = reconQC2
                    else:
                        pair1umiSeq = getReverseComplement(sequence_1[s][:umi_r])
                        pair2umiSeq = sequence_2[s][:umi_f]
                        # to add back UMI sequence.The full length after reconstruction has umi
                        reconstructedPair1 = reconstructedPair1 + pair1umiSeq
                        reconstructedPair2 = pair2umiSeq + reconstructedPair2
                        umiQC1 = R1_qc[sample][s][:umi_r]
                        umiQC2 = R2_qc[sample][s][:umi_f]
                        # to add back UMI qc. The full length qc after reconstruction has umi
                        reconQC1 = reconQC1 + umiQC1[::-1]
                        reconQC2 = umiQC2 + reconQC2
                        p1_aligned[s] = reconstructedPair2
                        p2_aligned[s] = reconstructedPair1
                        P1QC[s] = reconQC2
                        P2QC[s] = reconQC1
                    if overlap >= 0:
                        overlapQC = []
                        overlapSeq = ''
                        if pair1reverse == False:
                            for ov in range(overlap):
                                if reconQC1[-overlap+ov] >= reconQC2[ov]:
                                    overlapQC.append(reconQC1[-overlap+ov])
                                    overlapSeq += reconstructedPair1[-overlap+ov]
                                else:
                                    overlapQC.append(reconQC2[ov])
                                    overlapSeq += reconstructedPair2[ov]
                            full = reconstructedPair1[:len(reconstructedPair1)-overlap] + overlapSeq + reconstructedPair2[overlap:]
                            fullQC = reconQC1[:len(reconQC1)-overlap] + overlapQC + reconQC2[overlap:]
                            P1QC[s] = reconQC1
                            P2QC[s] = reconQC2
                        else: #if pair1 is revers
                            for ov in range(overlap):
                                if reconQC2[-overlap+ov] >= reconQC1[ov]:
                                    overlapQC.append(reconQC2[-overlap+ov])
                                    overlapSeq += reconstructedPair2[-overlap+ov]
                                else:
                                    overlapQC.append(reconQC1[ov])
                                    overlapSeq += reconstructedPair1[ov]
                            full = reconstructedPair2[:len(reconstructedPair2)-overlap] + overlapSeq + reconstructedPair1[overlap:]
                            fullQC = reconQC2[:len(reconQC2)-overlap] + overlapQC +reconQC1[overlap:]
                            P1QC[s] = reconQC2
                            P2QC[s] = reconQC1
                    else: # M is used to fill up the poistion that was not covered in sequencing
                        if pair1reverse == False:
                            full = reconstructedPair1 + 'M'*(-1*overlap) + reconstructedPair2
                            fullQC = reconQC1 + [0]*(-1*overlap) + reconQC2
                            P1QC[s] = reconQC1
                            P2QC[s] = reconQC2
                        else:
                            full =  reconstructedPair2 + 'M'*(-1*overlap) + reconstructedPair1
                            fullQC = reconQC2 + [0]*(-1*overlap) + reconQC1
                            P1QC[s] = reconQC2
                            P2QC[s] = reconQC1
                    fullSeq[s] = full
                    wholeQC[s] = fullQC
                    break
        print('Aligned:',sample)
        enhancerNumber[sample] = whichEnhancer
        allFullSeq[sample] = fullSeq
        allWholeQC[sample] = wholeQC
        pair1_qualityScore[sample] = P1QC
        pair2_qualityScore[sample] = P2QC
        pair1_alignment[sample] = p1_aligned # the first half seq
        pair2_alignment[sample] = p2_aligned # the second half seq
        pair1alignment_score[sample] = pair1_score # to record alignment scores for all reads in R1
        pair2alignment_score[sample] = pair2_score # to record alignment scores for all reads in R2
    return [enhancerNumber, allFullSeq, allWholeQC, pair1alignment_score, pair2alignment_score, pair1_alignment, pair2_alignment]

def getGCIndex(forwardPrimer,reversePrimer,originalEnhancerSeq):
    '''This function return the index of C within each GCH. UMI not included.
    To be noticed: the original sequences don't have umi, but umi exists in some enhancer reads.
    In this case, UMIs have to be trimmed from sequencing reads, if using this index to take
    conversion profiles at GC.'''
    UMI = getForwardReverseUMIList(forwardPrimer,reversePrimer)
    UMI_F, UMI_R = UMI[0], UMI[1]
    GCIndex = [] # stores the site where there is a C after G (based on original enhancer sequence).
    for e in range(len(forwardPrimer)):
        seq = originalEnhancerSeq[e].upper()
        umi_f,umi_r = len(UMI_F[e]),len(UMI_R[e])
        fp,rp, ll = len(forwardPrimer[e])-umi_f,len(reversePrimer[e])-umi_r, []
        for i in range(fp-1,len(seq)-rp-1):
            if seq[i] == 'G' and seq[i+1] == 'C' and seq[i+2] != 'G':
                #GC in primers are not included. GCGs are not considered
                ll.append(i+1)
        GCIndex.append(ll)
    return GCIndex
def readLengthCounter(alignmentResp1,alignmentResp2,DHSNumberList,animalNum, enhancerNumber):
    '''return the most frequent length in first and second half of sequences being sequenced'''
    p1p2frequentLength = [0]*animalNum
    for j in range(animalNum):
        pair1_alignment = alignmentResp1[j]
        pair2_alignment = alignmentResp2[j]
        numberOfRd = len(pair1_alignment)
        DHSlist = DHSNumberList[j]
        counterDHS = [0]*enhancerNumber
        for e in range(enhancerNumber):
            p1, p2 = [], []
            for s in range(numberOfRd):
                dhs = DHSlist[s]
                if dhs == e + 1:
                    p1.append(len(pair1_alignment[s]))
                    p2.append(len(pair2_alignment[s]))
            if len(p1)*len(p2) == 0:
                continue
            else:
                counterDHS[e] = [Counter(p1).most_common(1)[0][0], Counter(p2).most_common(1)[0][0]]
        p1p2frequentLength[j] = counterDHS
    return p1p2frequentLength
def GCnotInclude(p1Length, p2Length,umi_f,umi_r,enhancerLength,GCIndex):
    '''Some gc may not be covered in sequencing, those gc index will be excluded'''
    cutoff1 = p1Length - umi_f
    cutoff2 = enhancerLength - (p2Length - umi_r)
    GCnotIncludeList = []
    for i in GCIndex:
        if cutoff1 <= i and i < cutoff2:
            GCnotIncludeList.append(i)
    return GCnotIncludeList
def checkGT(seq,index):
    '''In GC check list, converted GT is marked as 1 for protection. GC as 0 not GY as -1.
    UMI not included.'''
    checked = [-1]*len(index)
    for m in range(len(index)):
        c = index[m]
        if seq[c] == 'T' and seq[c-1] == 'G':
            checked[m] = 1
        elif seq[c] == 'C' and seq[c-1] == 'G':
            checked[m] = 0
    return checked

# -r1 to indicate R1 file name (.txt). Each file name is a line
# -r2 to indicate R2 file name for the same sample (.txt) Each file name is a line
# -p to indicate primer file name. sep = '\t'. The headers are as the following order: F_primer_name	F_primer_sequence	R_primer_name	R_primer_sequence
# -t to indicate target orinal enhancer sequences. sep='\t'. The headers are as the following order: name	chr	start	end	sequence	library_run
# -o for alignment outputfile with all the reconstructed full length sequences

# here is to take input file names from command lines
cmdargs = sys.argv
print ("Running python file: ", cmdargs[0])
R1FILE = cmdargs[cmdargs.index('-r1')+1]
R2FILE = cmdargs[cmdargs.index('-r2')+1]
PRIMER_FILE_NAME = cmdargs[cmdargs.index('-p')+1]
ENHANCER_SEQ = cmdargs[cmdargs.index('-t')+1]
R1fileName = pd.read_csv(R1FILE, header = None).iloc[:,0].values
R2fileName = pd.read_csv(R2FILE, header = None).iloc[:,0].values
animalNum = len(R1fileName) # the number of samples
pathOfFastq = ''
if '/' in R1FILE:
    for i in range(len(R1FILE)-1,-1,-1):
        if R1FILE[i] == '/':
            pathOfFastq = R1FILE[:i+1]
            break
print(pathOfFastq+R1fileName[0])
OUTPUT = cmdargs[cmdargs.index('-o')+1]
print('output will save as',OUTPUT+'.sorted.all.tsv')

print('Reading paired-end fastq files')
R1, R2 = [0]*animalNum, [0]*animalNum
R1_sequence, R2_sequence = [0]*animalNum, [0]*animalNum
R1rawQC, R2rawQC = [0]*animalNum, [0]*animalNum
for i in range(animalNum):
    temp   = pd.DataFrame(pd.read_csv(pathOfFastq + R1fileName[i], sep='\t', header=None).values.reshape(-1, 4),
                         columns=['read_id', 'seq', '+', 'qual'])
    temp2  = pd.DataFrame(pd.read_csv(pathOfFastq + R2fileName[i], sep='\t', header=None).values.reshape(-1, 4),
                         columns=['read_id', 'seq', '+', 'qual'])
    R1[i] = temp
    R2[i] = temp2
    R1_sequence[i] = R1[i].iloc[:,1].values
    R2_sequence[i] = R2[i].iloc[:,1].values
    R1rawQC[i] = R1[i].iloc[:,3].values
    R2rawQC[i] = R2[i].iloc[:,3].values
qcNum = getQCnum(R1rawQC,R2rawQC)
R1_qc, R2_qc = qcNum[0], qcNum[1]
numberOfRdEachSample = [len(R1_sequence[i]) for i in range(len(R1_sequence))]
totalNumOfRead = sum(numberOfRdEachSample)
print('total number of reads processing: ',totalNumOfRead)

primers = pd.read_csv(PRIMER_FILE_NAME,sep = '\t')
forwardPrimer = primers.iloc[:,1].values
reversePrimer = primers.iloc[:,3].values
reversePrimer_revComplement = getRevPrimerRevComplement(reversePrimer)
UMI = getForwardReverseUMIList(forwardPrimer,reversePrimer)
UMI_F, UMI_R = UMI[0], UMI[1]
print('loaded primer sequences: ',PRIMER_FILE_NAME)

originalEnhancerInfo = pd.read_csv(ENHANCER_SEQ,sep = '\t')
originalEnhancerSeq = originalEnhancerInfo.iloc[:,4].values
enhancerNames = originalEnhancerInfo.iloc[:,0].values
enhancerNumber = len(originalEnhancerSeq)
newEnhancerSeq = getBisulfiteEnSeq(originalEnhancerSeq)
GCIndex = getGCIndex(forwardPrimer,reversePrimer,originalEnhancerSeq)
print('loaded original enhancer sequences from file: ',ENHANCER_SEQ)


#it contains the file path and the name of your file but with out adding a file formt (eg.not adding .tsv)
# this script adds a .sorted.all.tsv at the end of the file name
# generates file of bisulfite converted target sequences
file = pathOfFastq+'bisulfiteNewEnhancerSeq.txt'
with open(file,'w') as tfile:
    tfile.write('\n'.join(newEnhancerSeq))
alignmentRes = Align(animalNum,R1_sequence,R2_sequence,R1_qc,R2_qc,
                     forwardPrimer,reversePrimer,UMI_F,UMI_R,newEnhancerSeq)

p1p2frequentLength = readLengthCounter(alignmentRes[5],alignmentRes[6],alignmentRes[0],animalNum,enhancerNumber)
print('summarizing alignment...')
alignedAll = []
gcExcludelog = ['']*enhancerNumber
for j in range(animalNum):
    numberOfRd = len(R1_sequence[j])
    DHS_name, Chr, start, end = ['NA']*numberOfRd,['NA']*numberOfRd,['NA']*numberOfRd,['NA']*numberOfRd
    R1R2qcraw, umiForwardSeq, umiReverseSeq = ['NA']*numberOfRd,['-']*numberOfRd,['-']*numberOfRd
    charQC, checked, containNotGY, gcPositions = ['NA']*numberOfRd, ['NA']*numberOfRd, ['NA']*numberOfRd, ['NA']*numberOfRd
    for s in range(numberOfRd):
        dhs = alignmentRes[0][j][s] # the dhs index started from 1.
        qc = alignmentRes[2][j][s] # an int list of quality score
        if dhs > 0:
            gcindex = GCIndex[dhs-1]
            seq = alignmentRes[1][j][s]
            umi_f, umi_r = len(UMI_F[dhs-1]), len(UMI_R[dhs-1])
            p1Length, p2Length = p1p2frequentLength[j][dhs-1][0], p1p2frequentLength[j][dhs-1][1]
            GCnotIncludeList = GCnotInclude(p1Length,p2Length,umi_f,umi_r,len(originalEnhancerSeq[dhs-1]),gcindex)
            if len(GCnotIncludeList) > 0:
                gcExcludelog[dhs-1] = 'Notice: DHS'+str(dhs)+', gc were not sequenced and removed from analysis:'+str(GCnotIncludeList)
                gcindex = [gc for gc in gcindex if gc not in GCnotIncludeList]
            gcPositions[s] = gcindex
            binary = checkGT(seq[umi_f:len(seq)-umi_r],gcindex)
            checked[s] = binary
            if umi_f > 0 and umi_r == 0:
                umiForwardSeq[s] = seq[:umi_f]
            elif umi_r > 0 and umi_f == 0:
                umiReverseSeq[s] = seq[len(seq)-umi_r:]
            elif umi_f > 0 and umi_r > 0:
                umiForwardSeq[s] = seq[:umi_f]
                umiReverseSeq[s] = seq[len(seq)-umi_r:]
            DHS_name[s] = originalEnhancerInfo.iloc[dhs-1,0]
            Chr[s] = originalEnhancerInfo.iloc[dhs-1,1]
            start[s] = originalEnhancerInfo.iloc[dhs-1,2]
            end[s] = originalEnhancerInfo.iloc[dhs-1,3]
            notGYnumber = binary.count(-1)
            if notGYnumber == 0:
                containNotGY[s] = 0
            else:
                containNotGY[s] = str(notGYnumber) + '/' + str(len(gcindex))
            char = '' # use char for quality score
            for sc in qc:
                char+=chr(sc+33)
            charQC[s] = char
        # column R1R2rawqc contains qc of both R1 and R2 read. '{' curly bracket is the separator for two qc.
        R1R2qcraw[s] = R1rawQC[j][s] + '{' + R2rawQC[j][s]
    sample = [R1fileName[j][:R1fileName[j].index('_R')]]*numberOfRd
    df=pd.DataFrame({'DHS_number':alignmentRes[0][j], 'DHS_name':DHS_name, 'fullSeq_withUMI':alignmentRes[1][j],
                'fullQCwithUMI':charQC, '5primeUMISeq':umiForwardSeq, '3primeUMISeq':umiReverseSeq,
                'checkGT':checked, 'not_GY':containNotGY, 'GCIndex':gcPositions, 'read_id':R1[j]['read_id'],
                'chr':Chr, 'start':start, 'end': end, 'sample':sample})
    df['R1rawSeq'], df['R2rawSeq'], df['R1rawQC'], df['R2rawQC'] = R1_sequence[j], R2_sequence[j], R1rawQC[j], R2rawQC[j]
    df['R1alignment_score'], df['R2alignment_score'] = alignmentRes[3][j], alignmentRes[4][j]
    alignedAll.append(df)
alignedAll = pd.concat(alignedAll)
for i in gcExcludelog:
    if i != '': print(i)
print('sorting aligned sequences by samples and enhancers...')
sampleNames = [R1fileName[j][:R1fileName[j].index('_R')] for j in range(animalNum)]
sortedDfByEnhancer, allEnCountsByEnhancer, goodEnCountsByEnhancer, uniqCountsByEnhancer = [], [], [], []
not_GYCountsByEnhancer, primerMatchCountsByEnhancer = [], []
for e in range(enhancerNumber):
    enNumberAll, enNumberGood, uniqCounts = [0]*animalNum, [0]*animalNum, [0]*animalNum
    GRNum, primerMatchNum = [0]*animalNum, [0]*animalNum
    sortedDF = [0]*animalNum, [0]*animalNum
    sampleDf = []
    for s in range(animalNum):
        subdf = alignedAll[(alignedAll['DHS_number'] == e+1) & (alignedAll['sample'] == sampleNames[s])]
        subdfGood = subdf[subdf['not_GY']==0]
        subdfGR = subdf[subdf['not_GY']!=0]
        subdfPrimerMatch = alignedAll[(alignedAll['DHS_number'] == -e-1) & (alignedAll['sample'] == sampleNames[s])]
        GRNum[s], primerMatchNum[s] = subdfGR.shape[0], subdfPrimerMatch.shape[0]
        sampleDf.append(pd.concat([subdfGood,subdfGR]))
        enNumAll, enNumGood = subdf.shape[0], subdfGood.shape[0]
        enNumberAll[s], enNumberGood[s] = enNumAll, enNumGood
        allQC = subdfGood['fullQCwithUMI'].values
        qcList = []
        for a in range(len(allQC)):
            q = allQC[a]
            qc = []
            for sc in q:
                qc.append(ord(sc)-33)
            qcList.append(qc)
        barcode = []
        l, ll = subdfGood['5primeUMISeq'].values, subdfGood['3primeUMISeq'].values
        for n in range(enNumGood):
            if l[0]=='-' and ll[0] =='-':
                barcode = [i for i in subdfGood['fullSeq_withUMI'].values]
                break
            elif l[0] == '-':
                barcode.append(ll[n])
            elif ll[0] == '-':
                barcode.append(l[n])
            else:
                barcode.append(l[n]+ll[n])
        uniqIndex = np.unique(barcode)
        uniqCounts[s] = len(uniqIndex)
    uniqCountsByEnhancer.append(uniqCounts)
    allEnCountsByEnhancer.append(enNumberAll)
    goodEnCountsByEnhancer.append(enNumberGood)
    not_GYCountsByEnhancer.append(GRNum)
    primerMatchCountsByEnhancer.append(primerMatchNum)
    sortedDfByEnhancer.append(pd.concat(sampleDf))

sortedDfByEnhancer = pd.concat(sortedDfByEnhancer)
sortedDfByEnhancer.to_csv(OUTPUT+'.sorted.all.tsv',sep='\t',index=False)
print('saved sorted file')
print('alignment summary')
for s in range(animalNum):
    subdfNotAligned = alignedAll[(alignedAll['DHS_number'] == -100) & (alignedAll['sample'] == sampleNames[s])]
    notAligned = subdfNotAligned.shape[0]
    sampleRd = numberOfRdEachSample[s]
    am = [allEnCountsByEnhancer[e][s] for e in range(enhancerNumber)]
    uq = [uniqCountsByEnhancer[e][s] for e in range(enhancerNumber)]
    gr = [not_GYCountsByEnhancer[e][s] for e in range(enhancerNumber)]
    pm = [primerMatchCountsByEnhancer[e][s] for e in range(enhancerNumber)]
    grOveram = []
    for i in range(enhancerNumber):
        if am[i] != 0:
            grOveram.append(gr[i]/am[i])
        else:
            grOveram.append(0)
    alignedPer = 100*sum(am)/sampleRd
    notAlignedPer = 100*notAligned/sampleRd
    print('# '+sampleNames[s])
    print('total aligned read number and percentage: '+str(sum(am)) +' ' + str(alignedPer)+'%')
    print('not aligned read number and percentage: '+str(notAligned) +' ' + str(notAlignedPer)+'%')
    print('Counts of DHS: ', enhancerNames)
    print('total reads of each DHS: ',am)
    print('unique read of each DHS: ', uq)
    print('reads contain GR at GC positions: ', gr)
    print('percentage of reads conatain GR: ', grOveram)
    print('reads only matched to primer sequence: ', pm)
```
